# Stable large-scale network architecture and structured individual variability in the awake mouse brain

**DOI:** 10.64898/2026.10.30.762962

**Authors:** Yijuan Zou, Ruilin Tu, Changjiang Zhang, Dandan Hu, Xudong Hou, Wei Wang, Yuan Ji, Yuyuan Jiang, Qian Wang, Yuanlei Yue, Suijuan Zhong, Wencan Wu, Qian Wu, Xiaoqun Wang

## Abstract

Large-scale functional networks provide a scaffold for brain activity, but their population architecture and inter-individual variability remain poorly defined in the awake mouse brain. Here we introduce the Awake Mouse Connectome Project (AMCP), a population-scale multimodal MRI resource comprising 212 sex-balanced awake mice. Using this dataset, we generated high-precision in vivo tissue probability maps, derived a reproducible whole-brain functional network atlas and quantified structured variability in network connectivity across animals. The resulting atlas defined a stable large-scale architecture aligned with structural topology, while inter-individual functional connectivity variability followed organized connectional gradients rather than random dispersion. We further benchmarked the clustering-derived scaffold against conventional group independent component analysis (GICA) and release the GICA network atlas as an accompanying resource. AMCP further provides empirical scaling relationships linking sample size, scan duration and connectome precision. These analyses indicate that mean functional connectivity and functional connectivity variability have distinct sample-size requirements, providing a protocol-specific guidance for studies focused on group-average connectivity and inter-individual variability. Individualized network parcellations derived from the AMCP scaffold improved behavioural prediction relative to group-average features for selected behavioural domains, indicating functional relevance of subject-specific network architecture. Together, AMCP establishes an open dataset, tissue-template resource, clustering-derived and GICA functional atlases, and analysis toolbox for reproducible awake mouse connectomics and for translational studies linking circuit mechanisms to large-scale brain organization.

## Introduction

Large-scale MRI has transformed brain science by enabling non-invasive interrogation of brain organization from microstructure to functional networks^1^. In humans, large collaborative efforts, most notably the Human Connectome Project (HCP)^2,3^, have demonstrated the transformative impact of openly shared, high-quality neuroimaging databases, accelerating discoveries in brain-behavior relationships and disease mechanisms^4–6^. A cornerstone of this progress has been the development of standardized brain atlases and functional network parcellations derived from large resting-state fMRI (rsfMRI) cohorts, including widely used frameworks such as the canonical Yeo^7^, Schaefer^8^, Power^9^, and Gordon^10^ network definitions. These reference systems provide a common coordinate framework for comparing studies, integrating modalities, and translating connectomics findings into clinical and mechanistic insights^11,12^.

Understanding how brain networks give rise to behavior increasingly depends on population-level inference, where normative architecture and structured inter-individual variability can be distinguished from measurement noise^13,14^. In humans, this transition was enabled by large-scale connectome projects that established statistical regimes in which functional networks, individual differences, and brain-behavior associations could be modeled reproducibly^15–17^. However, this population-level transition has not yet occurred in mouse neuroimaging. Functional imaging in mice—the species central to mechanistic systems neuroscience—remains largely constrained to small-cohort feasibility studies. Although awake mouse fMRI provides a crucial bridge between cellular circuit manipulation and systems-level dynamics, the absence of population-scale datasets and quantitative design principles has limited reliable characterization of normative network architecture, hindered interpretation of inter-individual variability, and restricted reproducible brain-behavior mapping in rodents. Consequently, mouse connectomics has lacked the population framework that catalyzed human population connectomics.

Despite these advances, human neuroimaging remains constrained in its ability to probe causal mechanisms, as invasive manipulations are inherently limited and experimental control over biological variables is restricted. The mouse model provides a critical bridge across this translational gap, combining precise genetic tools, ex vivo validation, and scalable experimental design to link neural circuit mechanisms with systems-level brain dynamics^18^. To enhance cross-species comparability, the field has increasingly shifted toward awake imaging paradigms in rodents^17,19–21^. Awake mouse fMRI avoids the widespread suppression of neural and vascular activity induced by anesthesia, and thus provides a functional state more closely aligned with human resting-state^22,23^.

However, the promise of awake mouse fMRI has been limited by a critical lack of large-scale, reliable datasets and, consequently, the absence of a standardized functional brain atlas derived from such data. Most existing mouse functional mapping studies rely on small cohorts, frequently under anesthesia, and commonly employ approaches such as group independent component analysis (GICA), where results can be highly sensitive to subjective parameter selection^24–27^. The field therefore lacks a stable, data-driven functional network atlas and quantitative design principles for interpreting individual variability in the awake mouse brain.

Addressing this gap requires overcoming significant technical barriers associated with awake imaging, particularly stress- and motion-related artifacts. The present work builds upon a systematically optimized framework that enables stable and reproducible data acquisition in awake mice^28–31^. An optimized head–fixation system was combined with a gradual habituation protocol designed to minimize restraint-induced stress, with physiological measures confirming normalization of stress indicators^32^. Residual motion and non-neuronal noise were further reduced through refined head fixation procedures and targeted signal regression strategies, improving the stability and interpretability of BOLD measurements^29,30^. This technical foundation enabled population-scale awake-state functional mapping with low motion and measurable BOLD signal quality.

We therefore introduce AMCP, a standardized population resource for awake mouse connectomics. Across 212 mice, AMCP defines a reproducible network scaffold aligned with structural topology, quantifies structured inter-individual variability, estimates data-quantity requirements for mean functional connectivity (FC) and functional connectivity variability (FCV), and tests individualized parcellations for behavior prediction. The resource includes T2-weighted anatomical MRI, BOLD-EPI rsfMRI, DTI-EPI structural connectivity, probabilistic tissue templates, clustering-derived and GICA-derived functional atlases and the MIKey analysis toolbox.

## Results

### Establishing the statistical regime for awake mouse population connectomics

Population neuroscience requires enough data to separate shared network architecture and structured individual variability from measurement noise. AMCP was designed for this regime by aggregating multimodal MRI from 212 sex-balanced awake mice acquired with an optimized habituation and head-fixation protocol. The sex-balanced design was chosen prospectively to support comparisons of anatomical structure, functional connectivity and structural connectivity between female and male mice. Quality-control analyses confirmed low motion, stable signal-to-noise (SNR) characteristics and detectable functional responses, providing the basis for subsequent analyses of normative architecture and variability.

This AMCP dataset was acquired using a systematic, optimized experimental pipeline for awake mouse imaging^29–33^, as schematically outlined in Fig. 1a. To minimize state-dependent confounds, all behavioral assays for a given animal were completed within a one-week window following its MRI session, ensuring temporal proximity between imaging and behavioral assessment.

**Fig. 1.**
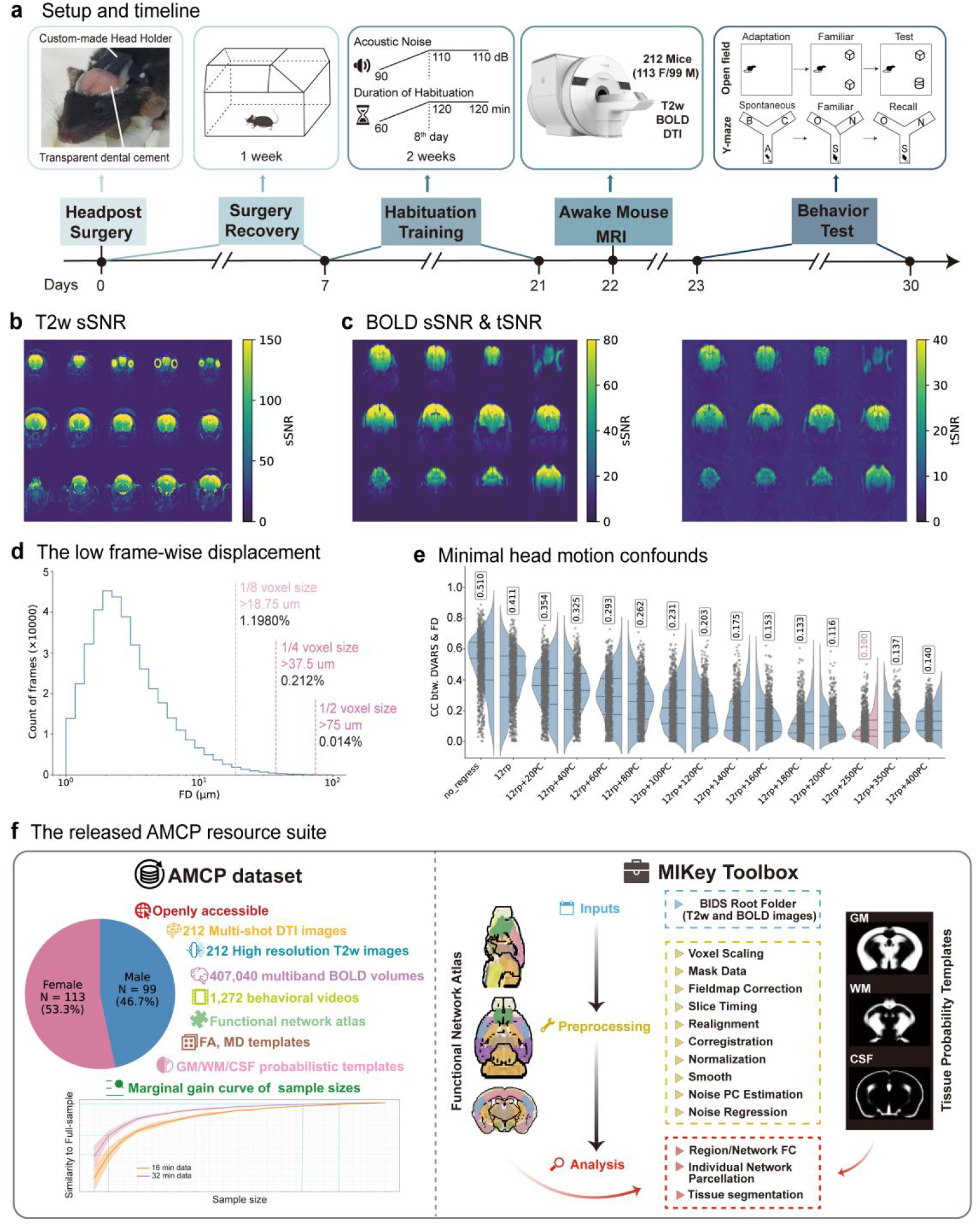
Overview of the AMCP resource, data quality and analysis framework. **a**, Experimental setup and timeline for awake mouse MRI acquisition and behavioral phenotyping. **b**, Representative sSNR map from a T2w anatomical image. **c**, Representative sSNR and tSNR maps from BOLD-EPI data. **d**, FD across all raw BOLD volumes, showing low head motion across the AMCP cohort. **e**, Evaluation of denoising performance across nuisance-regression strategies. The optimized strategy included 6 rp, their first-order temporal derivatives and 250 PCs derived from non-brain voxels. Denoising performance was quantified by the Pearson correlation between FD and DVARS, with lower FD-DVARS coupling indicating more effective reduction of non-neural variance related to motion, physiological fluctuations and scanner/system-related noise. Mean Pearson’s r values are labeled for each strategy. **f**, Overview of the released AMCP resource suite, including the open multimodal database of 212 awake mice, empirical marginal-gain analysis for sample-size estimation, the AMCP clustering-derived and GICA-derived functional network atlases and the MIKey toolbox for end-to-end analysis. . sSNR, spatial signal-to-noise ratio; tSNR, temporal signal-to-noise ratio;T2w, T2-weighted; FD, framewise displacement; rp, rigid-body realignment parameters; PC, principal component; DVARS, root-mean-square temporal derivative of voxel-wise BOLD signal. Source data are provided as a Source Data file

Data quality was rigorously evaluated using multiple quantitative metrics. To visualize spatial signal-to-noise ratio (sSNR) across the brain, we parcellated the whole brain into 1,000 regions. For each region, sSNR was computed as the mean signal intensity divided by the standard deviation of the background noise, generating comprehensive whole-brain sSNR maps. The T2w anatomical data showed complete brain coverage with a cortical sSNR of ∼150 (Fig. 1b). The BOLD-EPI functional data also demonstrated high quality, with cortical sSNR of ∼80 and an average temporal SNR (tSNR) of ∼40 (Fig. 1c). A systematic comparison with six publicly available awake mouse rsfMRI datasets containing C57-background mice further showed that the AMCP BOLD data exhibited high signal quality and low head motion. The AMCP SNR metrics exceeded those of the dataset acquired with a room-temperature surface coil, but were still inferior to those of the dataset acquired with a cryogenic surface coil (Extended Data Fig. 1a,b).

Head motion during scanning was minimal, with only 0.014% of BOLD volumes exhibiting framewise displacement (FD) greater than half the voxel size (75 µm), substantially lower than in other public datasets (Fig. 1d and Extended Data Fig. 2). FD was calculated from rigid-body realignment parameters estimated from the raw BOLD data before denoising or downstream preprocessing. We therefore used FD-DVARS coupling as an additional quality-control measure to quantify residual motion-related and global temporal variance, rather than as a direct measure of head displacement alone^29,30,33^. Before denoising, FD-DVARS coupling remained detectable despite the low absolute FD, because DVARS captures volume-to-volume changes in whole-brain BOLD signal that can arise from subtle motion effects, physiological fluctuations, arousal- or state-related global signal changes and non-brain signal sources. Consistent with this interpretation, the few transient FD spikes were temporally aligned with increases in DVARS and global signal (GS) fluctuations (Supplementary Fig. 1). We therefore optimized nuisance regression by minimizing FD-DVARS coupling across candidate strategies. The optimal model included six rigid-body realignment parameters, their first-order temporal derivatives and 250 principal components derived from non-brain voxels (Fig. 1e), indicating effective reduction of non-neural temporal variance while preserving the low-motion profile of the awake imaging data.

To test whether regression of non-brain principal components preserved biologically meaningful BOLD signal, we performed a complementary visual-stimulation fMRI experiment in three mice using the same awake setup and BOLD-EPI sequence. Robust visual-evoked activation was preserved after regression of 12 motion regressors alone, 12 motion regressors plus 20 non-brain PCs, and 12 motion regressors plus 250 non-brain PCs (Extended Data Fig. 3). The preserved activation pattern involved the expected visual pathway, including the primary visual cortex (V1), superior colliculus (SC) and lateral geniculate nucleus (LGN), indicating that the 250-PCs non-brain regression strategy reduces non-neural variance while retaining stimulus-evoked BOLD responses.

To directly assess physiological adaptation to the awake imaging preparation, we performed an additional validation experiment in an independent cohort of 19 mice that underwent the same head-post implantation surgery, 14-day habituation protocol and MRI scanning procedure. Body weight and tail-vein plasma corticosterone were measured longitudinally before habituation, during training and after MRI scanning (Supplementary Fig. 2). Plasma corticosterone was elevated during the early phase of habituation but returned to a level comparable to the pre-training baseline by day 7 (D7 versus D-1: paired t-test, t = 0.803, Holm-adjusted P > 0.05), indicating substantial adaptation to restraint and scanner-noise exposure. Corticosterone levels showed a further downward trend with continued habituation and were significantly lower than the pre-training baseline after MRI scanning (post-MRI versus D-1: paired t-test, t = 2.446, Holm-adjusted P = 0.049). This lower post-MRI level may partly reflect the fact that the pre-training baseline sample was collected during the animals’ first tail-vein blood draw, which could have induced an acute handling-related stress response. Body weight remained broadly stable across the 14-day habituation period. A small decrease was observed after MRI scanning (post-MRI versus pre-MRI: paired t-test, t = 5.378, Holm-adjusted P = 0.005), consistent with urination and defecation during the scan. Together, these data indicate that the habituation protocol substantially reduced stress-related physiological responses before awake fMRI acquisition.

The AMCP main cohort (N = 212) substantially exceeds existing public awake mouse datasets, which include no more than 15 C57-background animals (Extended Data Fig. 2b). This scale enabled construction of a population-derived functional atlas and empirical sample-size analyses; the raw data, derived resources and toolbox are publicly available (Fig. 1f).

### Constructing high-precision probabilistic tissue templates

Using the high-quality in vivo T2-weighted images from AMCP, we constructed tissue probability maps (i.e., gray matter/white matter/cerebrospinal fluid/other tissue/air, GM/WM/CSF/OT/AIR) with sharper tissue boundaries, more anatomically plausible segmentations, more stable tissue-volume estimates and stronger cross-dataset/cross-strain generalizability than the commonly used Keigo template^34^ (Fig. 2a,b).

**Fig. 2.**
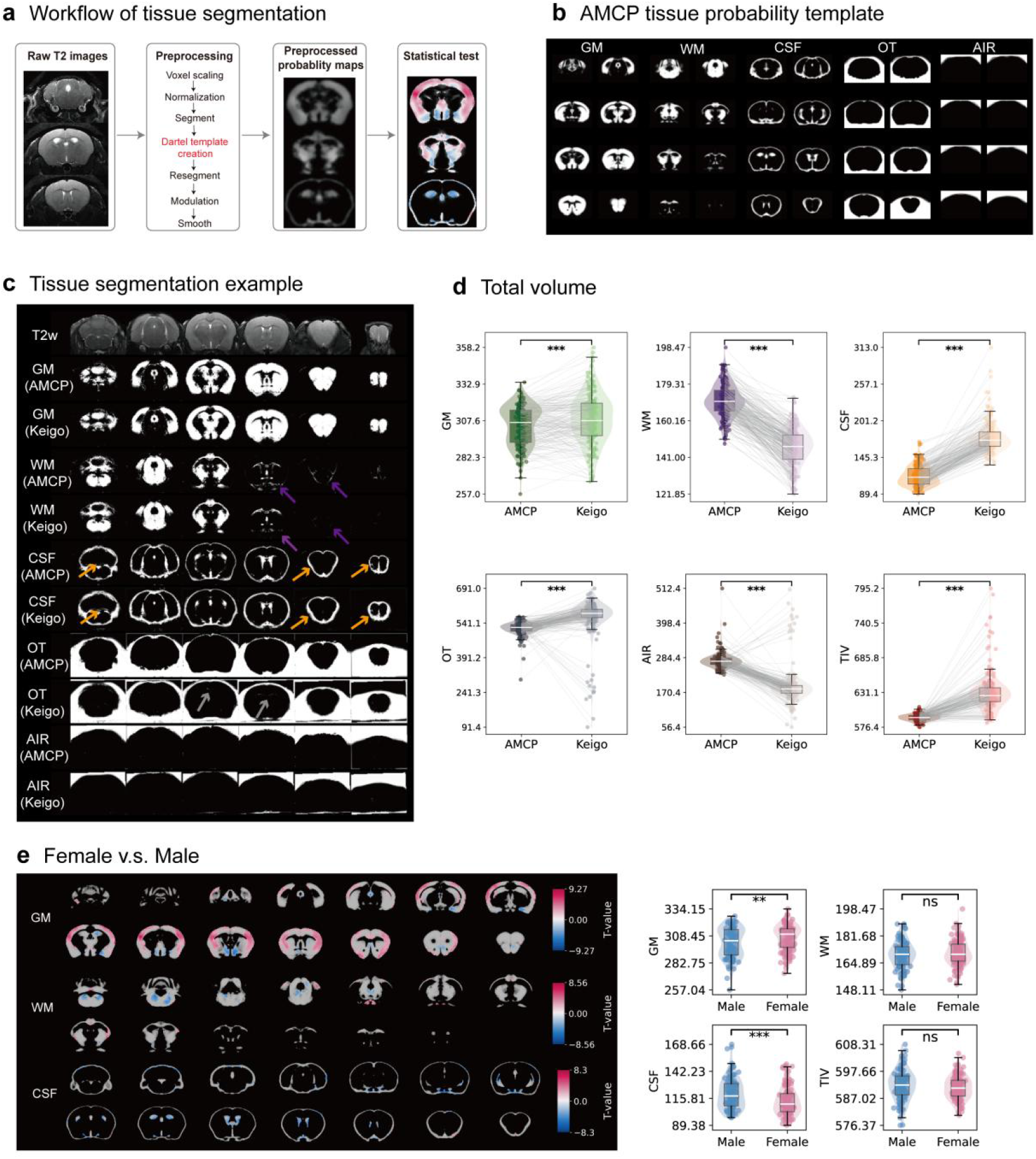
Construction and validation of AMCP probabilistic tissue priors. **a**, Workflow for constructing AMCP tissue probability maps. Initial tissue segmentation was performed using Keigo probabilistic priors, followed by population-template refinement with SPM DARTEL. The resulting AMCP tissue priors were used for subject-level segmentation and downstream tissue-composition analyses. **b**, AMCP probabilistic maps for GM, WM, CSF, OT and air. **c**, Representative comparison of tissue segmentation using AMCP and Keigo priors, overlaid on a T2w image. Arrows indicate regions where AMCP priors provided more anatomically plausible tissue boundaries than Keigo priors, including WM tracts (purple), CSF spaces (orange) and OT boundaries (gray). **d**, Quantitative comparison of tissue-volume estimates derived from AMCP and Keigo priors across all 212 mice. Each point represents one mouse. Template-wise comparisons were performed using paired two-sided t-tests; ***FDR corrected P < 0.001. **e**, Sex differences in tissue composition estimated using AMCP priors. Left, voxel-wise t-maps showing differences in GM, WM and CSF between female (n = 113) and male (n = 99) mice, thresholded at voxel-wise FDR-corrected P < 0.05. Right, group comparisons of total GM, WM, CSF and TIV. Each point represents one mouse. Sex comparisons were performed using two-sided two-sample t-tests; **FDR-corrected P < 0.01; ***FDR-corrected P < 0.001. GM, gray matter; WM, white matter; CSF, cerebrospinal fluid; OT, other tissue; TIV, total intracranial volume; FDR, false discovery rate. Source data are provided as a Source Data file.

Direct comparison of segmentation outcomes revealed a clear improvement in segmentation quality using the AMCP template (Fig. 2c). For example, in ventral forebrain regions where DTI data show a well–defined WM skeleton (Extended Data Fig. 4e) and high fractional anisotropy (Extended Data Fig. 4f), indicating dense white matter tracts, the Keigo template consistently failed to correctly classify these fibers as WM (Fig. 2c, purple arrow). In contrast, the AMCP template more accurately delineated these structures. Furthermore, the AMCP template produced sharper boundaries for CSF compartments, including structures such as the cerebral aqueduct, while the Keigo template tended to over-segment the surrounding subarachnoid CSF space (Fig. 2c, orange arrow). In comparable regions, the Keigo template also showed additional local misclassifications in adjacent tissue (Fig. 2c, gray arrow). Beyond the representative examples shown in Fig. 2c, quantitative analysis across all 212 mice showed that Keigo-based segmentation produced broader and more outlier-prone tissue-volume distributions than AMCP-based segmentation (Fig. 2d). Paired comparisons further confirmed systematic differences in estimated tissue volumes between the two priors across GM, WM, CSF, OT, AIR and total intracranial volume (TIV) (two-sided paired t-tests, n = 212; FDR-corrected across the six comparisons, all P < 0.001).

The AMCP template also generalized beyond the primary cohort. It performed well in an independent C57BL/6 dataset (Extended Data Fig. 5a) and produced anatomically plausible segmentations in C3H/He, DBA/2 and BALB/cBy mice (Extended Data Fig. 5b-d), despite being constructed only from C57BL/6 animals. This cross-strain performance exceeded that of the Keigo template, which was built from a multi-strain composite including these strains^34^.

Together, these analyses establish the AMCP tissue probability maps as a more anatomically plausible and stable resource for automated tissue segmentation. We then used AMCP-derived segmentations to compare tissue composition between female (N = 113) and male (N = 99) mice (Fig. 2e). Females showed larger cortical GM but smaller subcortical GM, lower WM volume in the interbrain and hindbrain, and smaller CSF volumes. At the global level, females had higher total GM volume and lower total CSF volume, whereas total WM volume and TIV did not differ significantly (Fig. 2e; two-sided two-sample t-tests, FDR-corrected P < 0.01 for GM and P < 0.001 for CSF). Inspection of mice with high versus low whole-brain GM volume confirmed that the observed bimodality was not driven by segmentation error (Supplementary Fig. 3).

### Defining the normative functional network scaffold of the awake mouse brain

We next defined a group-level functional network scaffold for the awake mouse brain. Following the Yeo framework^7^, we clustered voxels by similarity in connectivity profiles and evaluated split-half stability, while adapting the implementation to whole-brain mouse data. Specifically, we computed voxel-to-region FC matrices for 149,112 voxels across multiple initial region definitions, including Allen Mouse Brain Atlas (AMBA)-based parcellations (214, 428, 604 and 1,184 regions) and uniform parcellations (750, 1,000, 1,200 and 1,500 regions) (Fig. 3a). This stability framework allowed us to compare anatomical and spatially uniform initializations under the same criterion before selecting the final atlas solution.

**Fig. 3.**
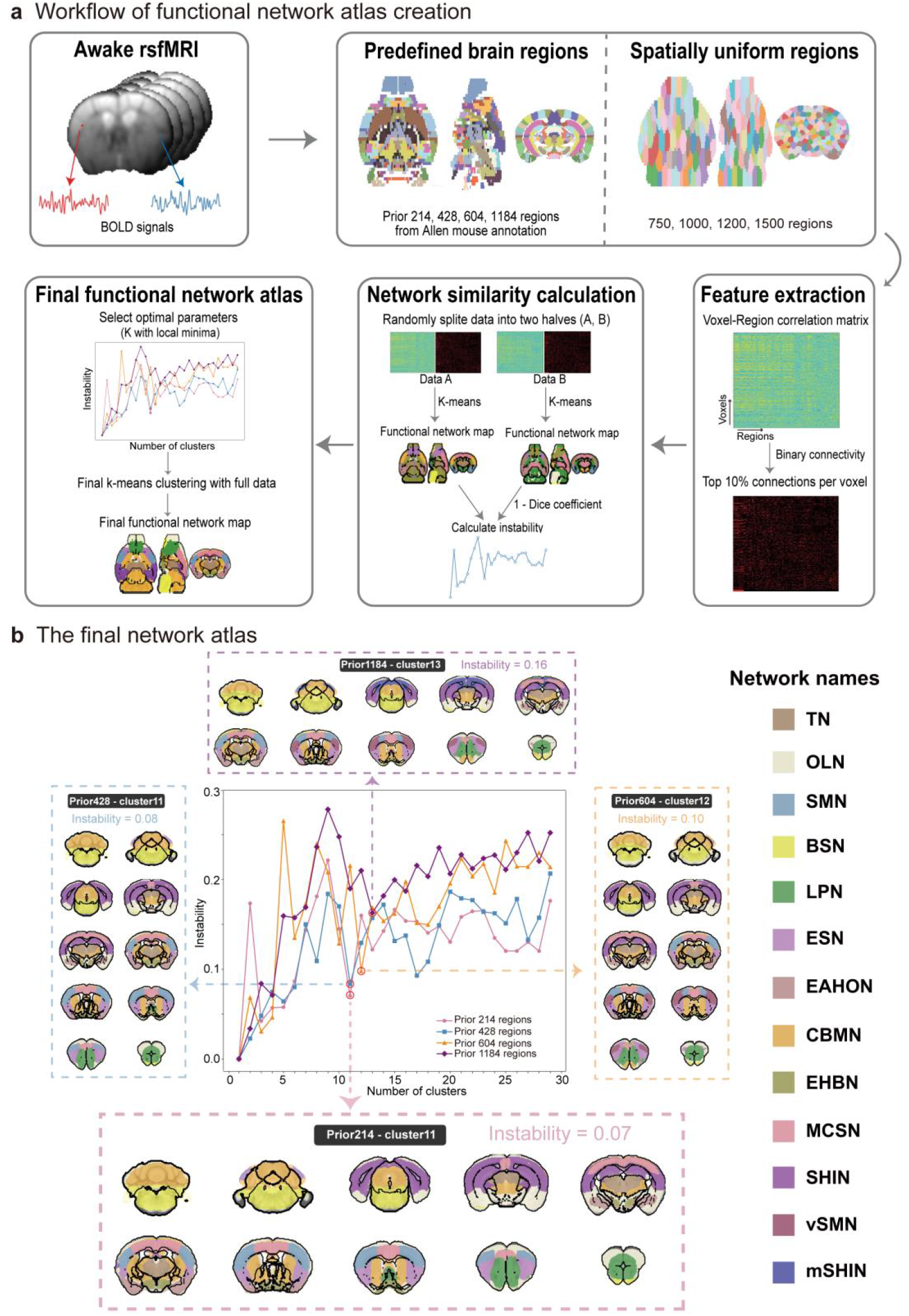
Derivation of the group-level AMCP functional network atlas. **a**, Workflow for atlas construction. Whole-brain voxel-to-region correlation matrices were computed for all rsfMRI runs (212 mice × 4 runs) and split into two independent subject-level halves. Within each half, voxels were clustered according to their connectivity profiles using k-means clustering across candidate region definitions and network numbers. Split-half reproducibility was quantified using the Dice coefficient, with 1 - Dice used as the instability metric. Each parameter combination was repeated ten times, and the solution at the elbow of the instability curve was selected. The final atlas was generated by applying k-means clustering with the selected parameters to the full dataset. The pipeline was adapted from Yeo et al. **b**, Spatial topography of the AMCP 11-network atlas. The selected solution was derived using a 214-region AMBA prior and corresponded to a stable elbow point on the instability curve. The 11 networks were TN, OLN, SMN, BSN, LPN, ESN, EAHON, CBMN, EHBN, MCSN and SHIN. AMBA, Allen Mouse Brain Atlas; TN, thalamus network; OLN, olfactory-limbic network; SMN, sensorimotor network; BSN, brainstem network; LPN, limbic-prefrontal network; ESN, extended salience network; EAHON, extended amygdala-hypothalamic-olfactory network; CBMN, cerebellar-basal ganglia-motor network; EHBN, extended hypothalamic-basal forebrain network; MCSN, medial cortical-sensorimotor network; SHIN, sensory-hippocampal-integration network. Source data are provided as a Source Data file.

Importantly, the split-half analysis used for atlas construction was performed at the subject level. In each repetition, animals were randomly divided into two independent groups, and all data from a given animal were assigned to only one half. Thus, the resulting stability estimates reflect agreement between independently derived solutions rather than any reuse of data across halves. Consistent with the original Yeo algorithm^7^, clustering stability decreased as the number of estimated networks increased, reflecting the expansion of the solution space and the increasing complexity of the clustering problem (Fig. 3b and Extended Data Fig. 6). Across region-definition schemes, although the exact location of the inflection point on the instability curve varied modestly, a clear and reproducible optimum emerged (Fig. 3b and Extended Data Fig. 6). Notably, Allen atlas-based parcellations consistently yielded lower instability than uniform parcellation schemes (Fig. 3b and Extended Data Fig. 6). Together with the consideration that voxels within Allen-defined anatomical regions are more likely to exhibit coherent signal fluctuations, we therefore adopted the Allen-prior scheme for constructing the final functional atlas.

We ultimately focused on the 11-network solution derived from the prior parcellation of 214 anatomical regions. This solution was selected because it corresponded to a distinct inflection point on the instability curve, representing a trade-off between relatively low instability and a functionally interpretable coarse-grained partition of the brain (Fig. 3b). Its robustness was further supported by cross-validation using a finer prior parcellation of 428 regions, which identified an almost identical inflection point at 11 networks. Moreover, the spatial topography of the 11 networks derived from the 428-region parcellation was highly similar to that derived from the 214-region parcellation (Fig. 3b), indicating that the large-scale network organization was not strongly dependent on this particular prior definition. The convergence of these solutions suggests that the 11-network architecture captures a stable large-scale property of the awake mouse functional connectome under the present analytical framework. Network nomenclature for the AMCP 11-network atlas was guided by the regional composition of each network based on Allen CCFv3 anatomical labels (Supplementary Fig. 4).

In line with the common practice of providing multiple scalable solutions (for example, the Yeo 7- and 17-network atlases)^7^, we also generated a complementary fine-grained 17-network atlas from a secondary optimal parameter set (Supplementary Fig. 5). While preserving the global architecture of the 11-network atlas, this solution reveals finer functional subdivisions, including separation of the Sensory-Hippocampal-Integration Network (SHIN) into cortical and hippocampal modules and division of the Brainstem Network (BSN) and Sensorimotor Network (SMN) into dorsal and ventral subnetworks. This additional release allows users to select analysis granularity according to their scientific objectives.

### Validation of AMCP 11 functional network atlas

We performed multi-faceted validation analyses to assess the robustness and biological relevance of the derived AMCP 11-network solution. First, we examined sensitivity to preprocessing choices. The spatial organization of the 11 networks remained highly similar across different nuisance-regression strategies, including pipelines with or without GS and CSF regressors (Fig. 4a and Extended Data Fig. 7a). This result indicates that the major large-scale clustered network architecture is not strongly dependent on the inclusion of these debated regressors, supporting the robustness of the atlas across common preprocessing variations.

**Fig. 4.**
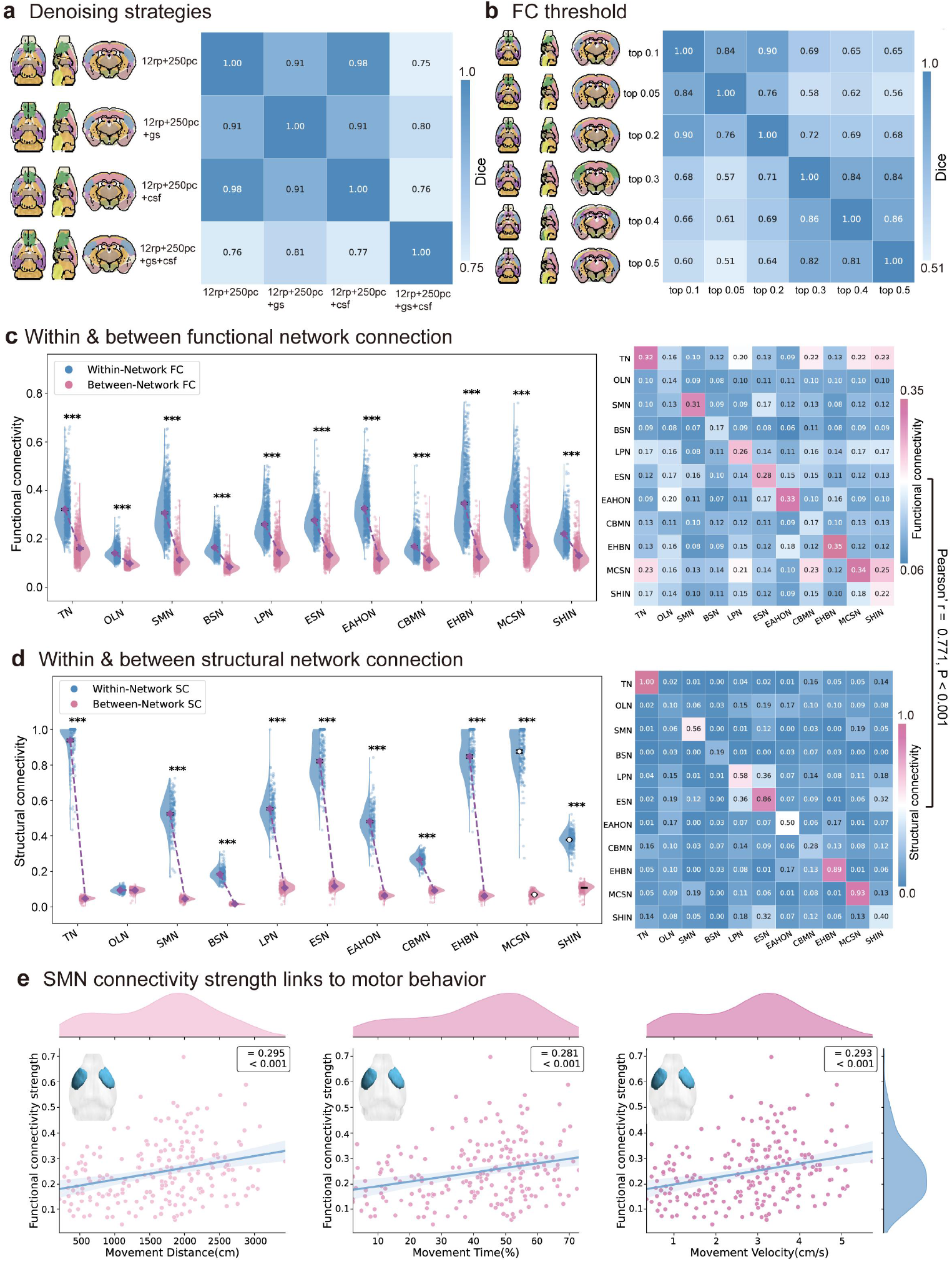
Validation of the AMCP 11-network functional atlas. **a**, Robustness of the atlas to nuisance-regression strategy. **b**, Robustness of network topography to the FC threshold used to retain voxel-to-region connections before clustering. **c**, Within- and between-network FC. Left, mean within-network FC was significantly higher than mean between-network FC across all 212 mice. Each point represents one mouse; violin plots show distributions; black points and error bars indicate mean ± SEM. Significance was assessed using paired two-sided t-tests with FDR correction across networks. Right, group-averaged FC matrix. **d**, Within- and between-network SC derived from DTI. Plotting and statistical conventions are as in c. **e**, Association between total SMN FC and three locomotion-related behavioral measures, assessed using Pearson correlation. ***FDR-adjusted P < 0.001. FC, functional connectivity; SC, structural connectivity; SMN, sensorimotor network. Source data are provided as a Source Data file.

Second, we evaluated the impact of the functional-connectivity threshold used to sparsify the voxel-to-region correlation matrix prior to clustering (the main analysis retained only the top 10% of strongest connections^7^). Network similarity remained high (Dice coefficient ∼0.8-0.9) when the retained connections were restricted to a relatively selective subset of strong edges (i.e., the top 5% to 20% of connections), indicating that the core large-scale network topology is well captured within this range (Fig. 4b and Extended Data Fig. 7b). In contrast, applying more inclusive thresholds that retained a larger proportion of weaker edges (e.g., the top 30% to 50%) reduced similarity to the reference atlas (Dice coefficient ∼0.6-0.7, Fig. 4b). We interpret this pattern not as instability of the core atlas, but as evidence that the strongest correlations provide the most stable basis for defining large-scale functional communities, whereas progressively weaker edges introduce greater variability and reduce clustering specificity. Thus, retaining the top 10% of connections represents a practical balance between capturing stable community structure and excluding the majority of weaker, less reliable correlations, consistent with common practice in rsfMRI studies^7,35^. Together, these analyses indicate that the major spatial topography of the AMCP 11-network atlas is preserved across preprocessing choices and remains stable across selective connectivity thresholds, as also illustrated by the slice-wise visualizations in Extended Data Fig. 7.

To further evaluate the biological validity of the atlas, we examined its correspondence with structural connectivity (SC) and behavior. We first tested the fundamental property of stronger within-network than between-network connectivity. Analysis confirmed that within-network FC was significantly stronger than between-network FC (Fig. 4c). Critically, this key feature of functional modularity was also reflected in the structural connectome: within-network SC, derived from DTI, was significantly stronger than between-network SC (Fig. 4d), supporting the plausibility of the functional network definitions at the network level. The group-averaged FC and SC matrices showed a significant edge-wise association at the network level (Pearson’s r = 0.771, P < 0.001), providing additional evidence for correspondence between functional and structural organization without implying one-to-one coupling at the individual-edge level. This observation is broadly consistent with prior studies reporting structured correspondence between functional and anatomical organization in the mouse brain^16,36,37^.

We next asked whether a functionally interpretable group-level network measure was related to behavior. We found that the total FC strength of the SMN—its aggregate connectivity with all other networks—was significantly correlated with three core locomotion metrics: total movement distance, movement time, and average velocity (Fig. 4e). This analysis was intended as a direct functional validation of the group-level atlas, showing that the network scaffold defined by the AMCP 11-network solution captures behaviorally meaningful variation at the level of a biologically interpretable system.

Collectively, these analyses support the AMCP 11-network solution as a robust large-scale awake-mouse functional atlas within the present framework, while also indicating that local boundary placement in lower-tSNR regions should be interpreted more cautiously than the core network architecture.

### Benchmarking the clustering-derived atlas against conventional GICA

Because GICA is still widely used to identify functional networks in mouse rsfMRI^24–27^, we benchmarked the clustering-based atlas against a conventional GICA framework (Fig. 5). We first evaluated GICA solutions across multiple model orders using ICASSO stability and repeated-run spatial correspondence, rather than selecting the dimensionality by visual inspection alone. This analysis showed that component stability was not sufficient to guarantee reproducible network definition. Lower-order solutions merged anatomically distinct patterns into broad components, whereas higher-order decompositions increasingly split coherent networks and showed reduced run-to-run correspondence (Fig. 5a-e). The 20-component solution therefore represented a practical compromise, preserving anatomically interpretable large-scale components while maintaining comparatively stable spatial structure. Consistent with this choice, the 18-component solution showed broadly similar patterns but reduced anatomical specificity in several components, including mixed hemispheric or ventricular-adjacent signals (Supplementary Fig. 6).

**Fig. 5.**
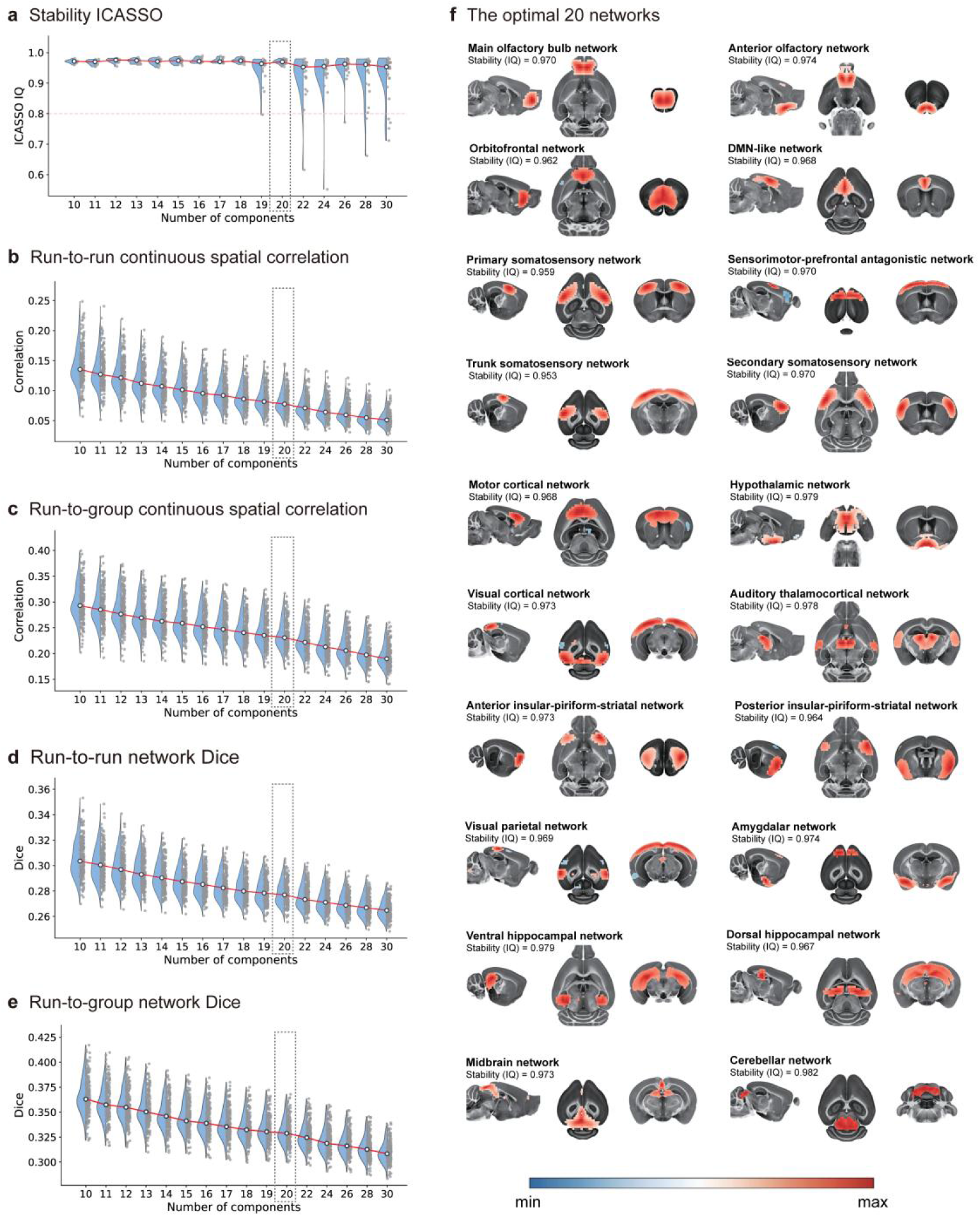
Stability-based evaluation of GICA-derived functional components. **a**, ICASSO stability across GICA model orders. Each point represents one independent component, and the 20-component solution is highlighted as the representative GICA model order used for comparison with the clustering-derived AMCP atlas. **b**, Run-to-run spatial correspondence across GICA model orders, quantified as Pearson correlation between subject-specific component maps from different runs without thresholding. **c**, Run-to-group spatial correspondence, quantified as Pearson correlation between each run-specific component map and the corresponding group-level component map without thresholding. **d**, Run-to-run spatial overlap, quantified using the Dice coefficient after binarizing subject-specific run-level component maps at the 80th percentile of their intensity distribution. **e**, Run-to-group spatial overlap, quantified using the same 80th-percentile binarization procedure. **f**, Spatial maps of the 20-component GICA solution. Component maps were converted to Z scores and thresholded at |Z| > 1.96 for visualization. Red and blue indicate positive and negative component loadings, respectively. GICA, group independent component analysis; ICASSO, independent component analysis reliability analysis. Source data are provided as a Source Data file.

Despite this optimization, the reproducibility of the GICA-derived networks remained limited. Across repeated awake rsfMRI runs, the 20-component GICA solution showed substantial variability in component overlap, particularly after threshold-based network binarization, with low run-to-run Dice similarity (approximately 0.24; Fig. 5d). Continuous spatial correlations also declined as model order increased, indicating that the instability was not solely a consequence of thresholding (Fig. 5b-e). By contrast, the clustering-derived atlas and its individualized network maps showed markedly higher consistency across repeated runs (Fig. 7b,c). These results indicate that, in the present awake mouse dataset, clustering provided a more reproducible group-level scaffold than conventional GICA for downstream functional network definition and individual-level parcellation.

### Structured inter-individual variability in mouse functional networks

Having established the group-level scaffold, we next quantified how individual brains diverge from it. Individualized parcellation preserved the global network architecture while revealing subject-specific variation in network boundaries and connectivity strength. These individualized features improved prediction for selected behavioral domains relative to group-average features, indicating that variability in awake mouse functional networks is structured and behaviorally informative rather than purely stochastic measurement noise.

We next examined the spatial organization of inter-individual functional connectivity variability (FCV) in the awake mouse brain, following the connectional-variability framework used to characterize individual functional variability and its structural correlates in the human connectome^5^. The SMN exhibited the highest within-network FCV, whereas the MCSN showed the highest total FCV, combining within- and between-network variability (Fig. 6a,c,d). This pattern extends prior awake-mouse work on individual variability and fingerprinting under a different analytical framework. Bergmann et al. (2020)^38^, using a smaller repeated-measurement cohort and a coarser cortical module scheme, reported relatively homogeneous variability across sensory and association systems. By contrast, our larger whole-brain network-level analysis revealed a structured spatial gradient of FCV. We therefore interpret the difference between the two studies as likely reflecting differences in sample size, parcellation granularity, spatial coverage and variability metrics, rather than as a direct contradiction. In addition, the spatial pattern of FCV was significantly correlated with structural communicability variability (SCV) derived from DTI (Fig. 6b,e), indicating network-level correspondence while remaining agnostic about precise edge-wise FC-SC coupling.

**Fig. 6.**
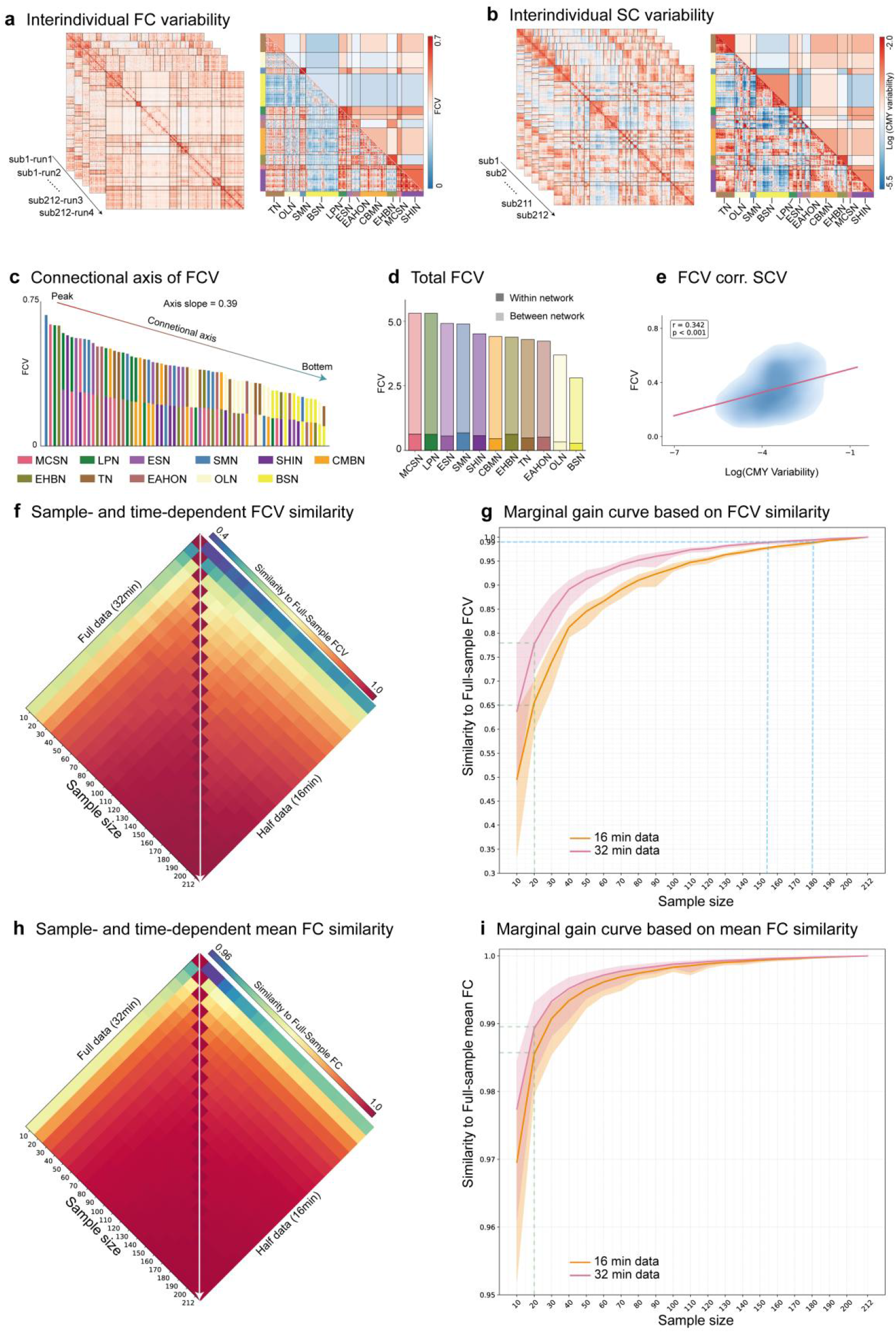
Network-level FCV architecture and data-driven sample-size estimation in awake mouse fMRI. **a**, Inter-individual FCV. Region-level functional connectomes from all imaging runs were used to estimate FCV. The resulting FCV matrix is shown at the region level in the lower triangle and summarized at the AMCP 11-network level in the upper triangle. **b**, SCV. Regional structural connectomes were used to compute communicability, and inter-individual variability in structural communicability is shown at the region level in the lower triangle and at the network level in the upper triangle. **c**, Connectional axis of network-level FCV. The 66 network-level FCV terms were ranked from highest to lowest. Single-color bars indicate within-network terms, and two-color bars indicate between-network terms; colors denote the networks involved. The axis slope was estimated by linear regression on the ranked FCV values and summarizes heterogeneity across the connectome. **d**, Total within- and between-network FCV for each network. **e**, Association between FCV and SCV across region-level connectome edges, assessed using Pearson correlation. **f**, Sample-and scan-duration-dependent similarity to the full-sample FCV matrix. Similarity was computed between subset-derived FCV matrices and the full-sample FCV matrix generated using all four runs (Runs 1-4, 32 min; left) or half of the data (Runs 1-2, 16 min; right). **g**, Marginal-gain curve for sample-size estimation based on FCV. Similarity to the full-sample FCV matrix is plotted as a function of sample size and scan duration. **h**, Sample- and scan-duration-dependent similarity to the full-sample mean FC matrix, computed as in **f**. **i**, Marginal-gain curve for sample-size estimation based on mean FC, computed as in **g**. FC, functional connectivity; FCV, functional connectivity variability; SCV, structural communicability variability; CMY, communicability. Source data are provided as a Source Data file.

We then summarized the spatial organization of FCV as a population-level connectional axis by ranking the 66 network-level FCV terms from highest to lowest (Fig. 6c). FCV declined from high-variability within-network terms and edges involving SMN, MCSN and LPN toward lower-variability terms involving the brainstem network (BSN). In keeping with the connectional-axis framework^5^, a linear slope fitted to the ranked FCV values was used as a compact summary of how strongly variability was differentiated across the connectome. A steeper slope indicates a larger separation between high- and low-variability network terms, whereas a flatter slope indicates a more uniform FCV distribution. Female and male mice showed broadly similar axes, with only a small slope difference in the present dataset (female: 0.388; male: 0.391; Supplementary Fig. 7). Because this analysis was not preregistered, the sex-related difference should be interpreted as hypothesis-generating.

To assess whether the observed FCV architecture contained a stable between-subject component, we quantified repeated-run reliability for the 66 network-level FC terms across the four runs acquired from each animal. Single-run reliability was moderate to high overall, with mean ICC(3,1) values of 0.736 for within-network terms and 0.742 for between-network terms, where ICC(3,1) denotes the reliability of a single run in preserving between-subject differences relative to within-subject/run-level fluctuation (Supplementary Fig. 8a,c). Averaged-measure reliability was substantially higher, with mean ICC(3,k) values of 0.917 for within-network terms and 0.920 for between-network terms; ICC(3,k) denotes the reliability of the measure averaged across repeated runs (Supplementary Fig. 8b,f). Variance decomposition showed that both between-subject and within-subject/error components contributed to network-level FC variability (Supplementary Fig. 8d,e). Thus, the FCV architecture reported here includes a substantial reproducible component, while still retaining measurable within-subject fluctuation that may reflect residual state-dependent or measurement-related effects.

To test whether FCV architecture and scaling were driven primarily by gradual state changes across the scanning session, we repeated the analysis separately for Runs 1-2 and Runs 3-4. The two network-level FCV matrices were highly similar (Pearson’r = 0.868, P < 0.001; Extended Data Fig. 9a-c), and their sample-size marginal-gain curves were nearly superimposable (Extended Data Fig. 9d,e). Together with repeated-measures reliability analyses (Supplementary Fig. 8), these results indicate that the main FCV architecture is not dominated by monotonic scan-time drift, although residual arousal, stress or state-dependent effects cannot be fully excluded without concurrent physiological measurements.

### Scaling relationships for connectome precision in awake mouse connectomics

We next quantified how sample size and scan duration affected the stability of connectome estimates in AMCP using a marginal-gain framework (Fig. 6f-i). For each subsample, we computed the similarity between the subsample-derived matrix and the corresponding full-sample AMCP estimate. Similarity increased nonlinearly with data quantity, with rapid early gains followed by progressively smaller improvements. These curves therefore provide protocol-specific scaling relationships between scan duration, cohort size and the stability of network-level measurements.

These marginal-gain curves address two related but distinct experimental design questions. The FCV curve (Fig. 6g) is most relevant for studies focused on inter-individual variability in functional connectivity, including FCV architecture, axes of variability, individual-difference mapping, or genotype-, treatment- or phenotype-associated changes in connectivity heterogeneity. Such analyses depend on estimating how connectivity varies across animals and therefore place stronger demands on cohort size. By contrast, the mean FC curve (Fig. 6i) is most relevant for studies focused on the group-average connectivity pattern, such as a normative mean connectome or average network-level FC. This curve quantifies how closely a smaller cohort’s mean FC matrix approximates the full-sample mean FC matrix, but does not by itself determine power for detecting between-group effects.

The data requirement was more pronounced for FCV than for mean FC. Similarity between subsampled FCV matrices and the full-sample FCV matrix increased with sample size, but approached the 0.99 similarity criterion only at relatively large cohort sizes (Fig. 6f,g). Under the AMCP acquisition and processing protocol, approximately 154 mice were needed when 32 min of rsfMRI data were available per mouse, compared with approximately 180 mice when 16 min were used. At n approximately 20, a cohort size common in rodent rsfMRI studies, FCV similarity was approximately 0.78 with 32 min of data and 0.65 with 16 min of data. These results indicate that analyses targeting population-level variability or variability axes are likely to require larger samples than analyses targeting group-average connectivity, at least under the present protocol and similarity criterion.

Because many rodent rsfMRI studies use smaller cohorts to estimate group-average FC, we next examined the marginal-gain curve for mean FC. In contrast to FCV, mean FC approached the full-sample estimate with fewer animals (Fig. 6h,i). With 20 mice, the subsampled mean FC matrix reached a similarity of approximately 0.986 using 16 min of data and approximately 0.99 using 32 min. Thus, under AMCP acquisition conditions, group-average FC was less sample-size dependent than FCV. This distinction suggests that studies focused on inter-individual variability, variability axes or heterogeneity-related phenotypes should be planned using the more conservative FCV curve rather than the mean FC curve.

Finally, we asked whether the AMCP-derived marginal-gain curves provided a useful reference for independent awake mouse rsfMRI datasets. Using the same analytical framework, we computed FCV and mean FC matrices for an independent AMCP validation cohort (AMCP-VC19; n = 19) and two publicly available awake mouse fMRI datasets, then quantified their similarity to the corresponding full-sample AMCP estimates (Extended Data Fig. 10). In AMCP-VC19, which was acquired using the same protocol as the main AMCP cohort, both the 16-min and 32-min estimates fell slightly below the similarity values expected from subsampling the main AMCP dataset at matched sample size and scan duration (Extended Data Fig. 10a,b). This modest reduction is expected in part because the marginal-gain curves were derived from subsamples drawn from the same animals used to define the full-sample reference, whereas AMCP-VC19 was fully independent.

The two public datasets, which differed from AMCP in EPI sequence parameters and data-quality profiles, showed lower similarity to the AMCP full-sample estimates than the AMCP subsampling curves at matched sample sizes. The Gutierrez-Barragan dataset (n = 10), which had lower data quality in our quality-control analyses, fell below the AMCP marginal-gain range for both FCV similarity (Pearson’s r = 0.183, P < 0.001) and mean FC similarity (Pearson’s r = 0.676, P < 0.001). The ds002551 dataset (n = 6), despite its higher data-quality profile, also fell below the AMCP range for FCV similarity (Pearson’s r = 0.145, P < 0.001) and mean FC similarity (Pearson’s r = 0.737, P < 0.001). These observations indicate that acquisition parameters, data quality and cohort-specific factors, potentially including housing and habituation procedures, can influence how closely an independent dataset matches the AMCP reference. The AMCP marginal-gain curves should therefore be interpreted as protocol-specific design references rather than universal sample-size thresholds. They estimate the stability of sample-derived connectome matrices relative to the AMCP full-sample reference, but do not replace power analyses for detecting between-group effects, which also depend on effect size, within-group variance and multiple-comparison burden. For studies that differ from AMCP in acquisition protocol, animal preparation or data quality, sample-size targets should be interpreted conservatively, and modestly larger cohorts may be warranted.

### Individualized functional network parcellation based on iterative optimization

After defining the group-level scaffold, we derived individualized functional networks using a modified iterative parcellation algorithm adapted from Wang et al.^39^. To improve convergence in high-dimensional mouse fMRI data, voxel time series were first averaged within AMBA prior-label units. Network assignments were then iteratively updated by correlating each unit with dynamically estimated network reference signals until assignment changes fell below a preset threshold or a maximum iteration number was reached (Fig. 7a). Network-level FC matrices were computed from the final individualized parcellations.

**Fig. 7.**
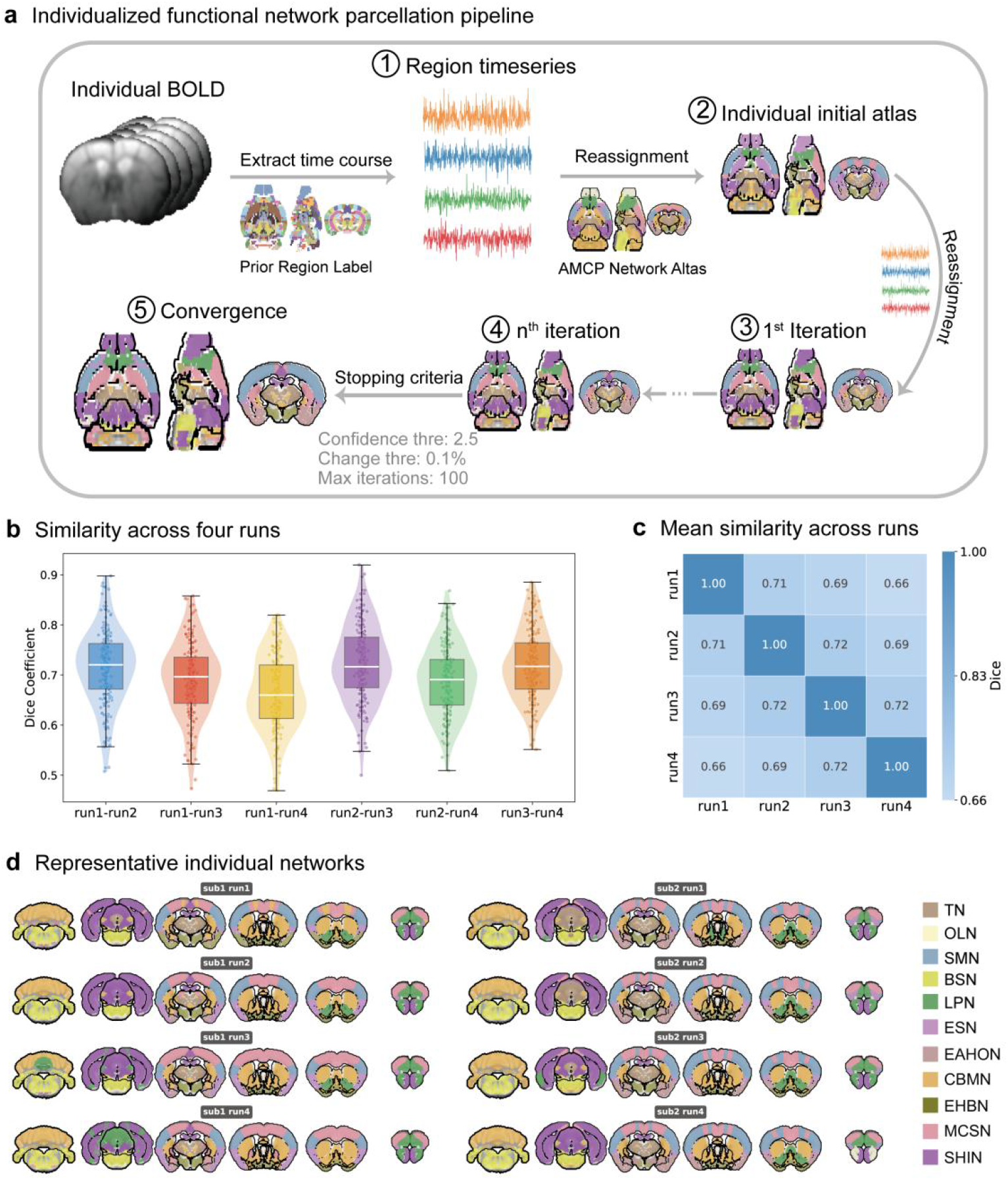
Individualized functional network parcellation in the awake mouse brain. **a**, Workflow of the iterative individualized parcellation algorithm, adapted from Wang et al. Initial units were defined using downsampling strategies, network reference signals were generated from the group-level atlas, and units were iteratively reassigned to the network with the highest temporal correlation until convergence. **b**, Within-subject similarity of individualized parcellation maps across four repeated scanning runs. c, Mean inter-run similarity of individualized network maps across all subjects, summarizing test-retest stability across repeated measurements. **d**, Representative individualized functional network maps from two mice across repeated runs, illustrating subject-specific and run-to-run variation in network topography. Source data are provided as a Source Data file.

Across four consecutive rsfMRI runs, individualized maps showed higher similarity between adjacent runs than between temporally distant runs (Fig. 7b,c), consistent with both stable individual organization and gradual state-dependent modulation during prolonged awake scanning. A comparable level of reproducibility was observed in the independent AMCP validation cohort of 19 mice, with consistent within-subject Dice similarity across repeated runs (mean Dice = ∼0.65, compared with ∼0.7 in the main AMCP cohort; Supplementary Fig. 9). Representative subjects illustrated localized boundary shifts around motor-related networks while preserving the overall scaffold (Fig. 7d), in line with the high FCV observed in MCSN and SMN (Fig. 6c,d).

### Individualized network features enhance behavioral prediction

Using behavioral phenotyping from all 212 mice (Supplementary Fig. 10 and Supplementary Table 1), we tested whether individualized network architecture improves behavior prediction relative to group-average features. Each behavioral outcome was modeled independently using network-level FC features from individualized parcellations (Indv-Net) or the group-average atlas (Avg-Net), with leakage-safe feature selection, ridge regression with nested cross-validation and permutation testing (Fig. 8a).

**Fig. 8.**
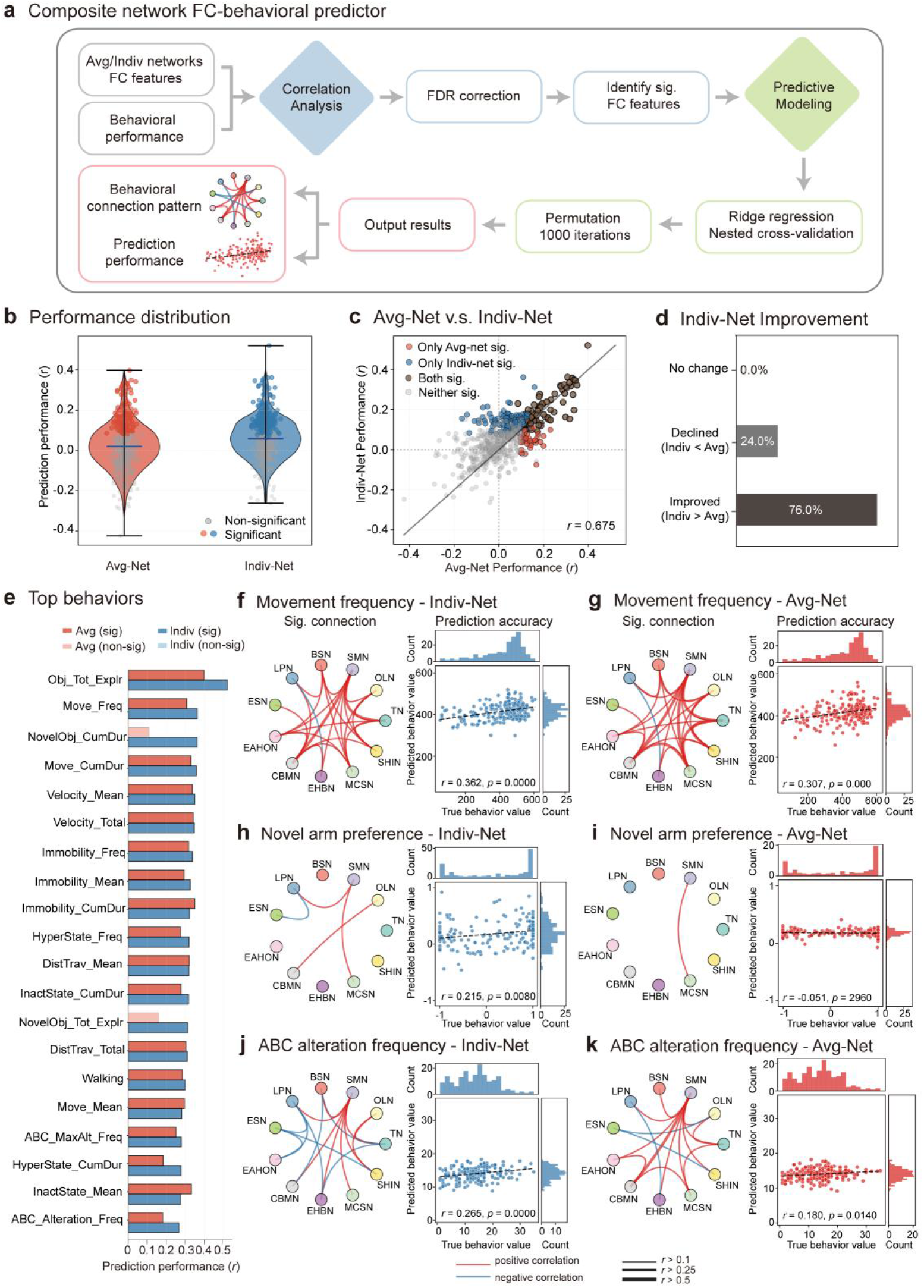
Behavioral prediction using individualized and group-average network features. **a**, Computational workflow for predicting behavior from network-level FC features derived from either the group-average atlas (Avg-Net) or individualized parcellations (Indv-Net). For each behavioral outcome, behavior-associated connections were selected by mass-univariate screening, and prediction was performed using ridge regression with nested cross-validation. Prediction significance was assessed by permutation testing. **b-d**, Comparison of prediction performance between Avg-Net and Indv-Net features. **b**, Distribution of prediction accuracy, quantified as Pearson correlation between observed and predicted behavioral scores. **c**, Number of behavioral outcomes with significant prediction performance for each feature set (permutation test, P < 0.05). **d**, Proportion of behavioral outcomes for which Indv-Net features improved, reduced or did not change prediction accuracy relative to Avg-Net features. **e**, Top 20 behavioral outcomes ranked by prediction accuracy using Indv-Net features. **f-k**, Representative prediction examples for movement frequency (**f**,**g**), novel arm preference (**h**,**i**) and ABC arm alternation frequency (**j**,**k**). For each behavior, significant network connections and prediction correlations are shown for Indv-Net features (left column) and Avg-Net features (right column). FC, functional connectivity; Avg-Net, group-average atlas-based network features; Indv-Net, individualized parcellation-based network features. Source data are provided as a Source Data file.

Indv-Net features produced an overall shift toward higher prediction accuracy (Fig. 8b), a larger number of significant behavioral predictions (Fig. 8c), and more outcomes with improved than reduced performance relative to Avg-Net features (Fig. 8d). The top-ranking behavioral outcomes are shown in Fig. 8e.

Performance varied by behavioral domain. Indv-Net and Avg-Net models showed overlapping connection patterns (Fig. 8f-k), suggesting that individualized maps refine rather than replace major group-level signatures. Both feature sets predicted basic motor metrics such as movement frequency and mean velocity well (Fig. 8f,g and Supplementary Fig. 11a,b), whereas Indv-Net showed an advantage for higher-order measures such as novel arm preference and ABC arm alternation frequency (Fig. 8h-k). For some behaviors, including head scanning, Avg-Net performed better (Supplementary Fig. 11i,j), indicating that certain behavioral variance is captured more parsimoniously by stable population-typical features.

Thus, individualized and group-average features capture complementary aspects of behaviorally relevant network organization: individualized parcellations improve sensitivity to subject-specific heterogeneity, whereas group-average features remain useful for behaviors anchored in more conserved circuits.

### Infrastructure enabling reproducible population connectomics in awake mice

AMCP also provides practical infrastructure for reproducible awake mouse connectomics: tissue probability maps, a preprocessing framework, the AMCP functional atlas and the MIKey open-source toolbox for individualized parcellation (Fig. 9). Together with public raw and processed data, these resources translate the population framework into reusable tools for cross-laboratory analysis.

**Fig. 9.**
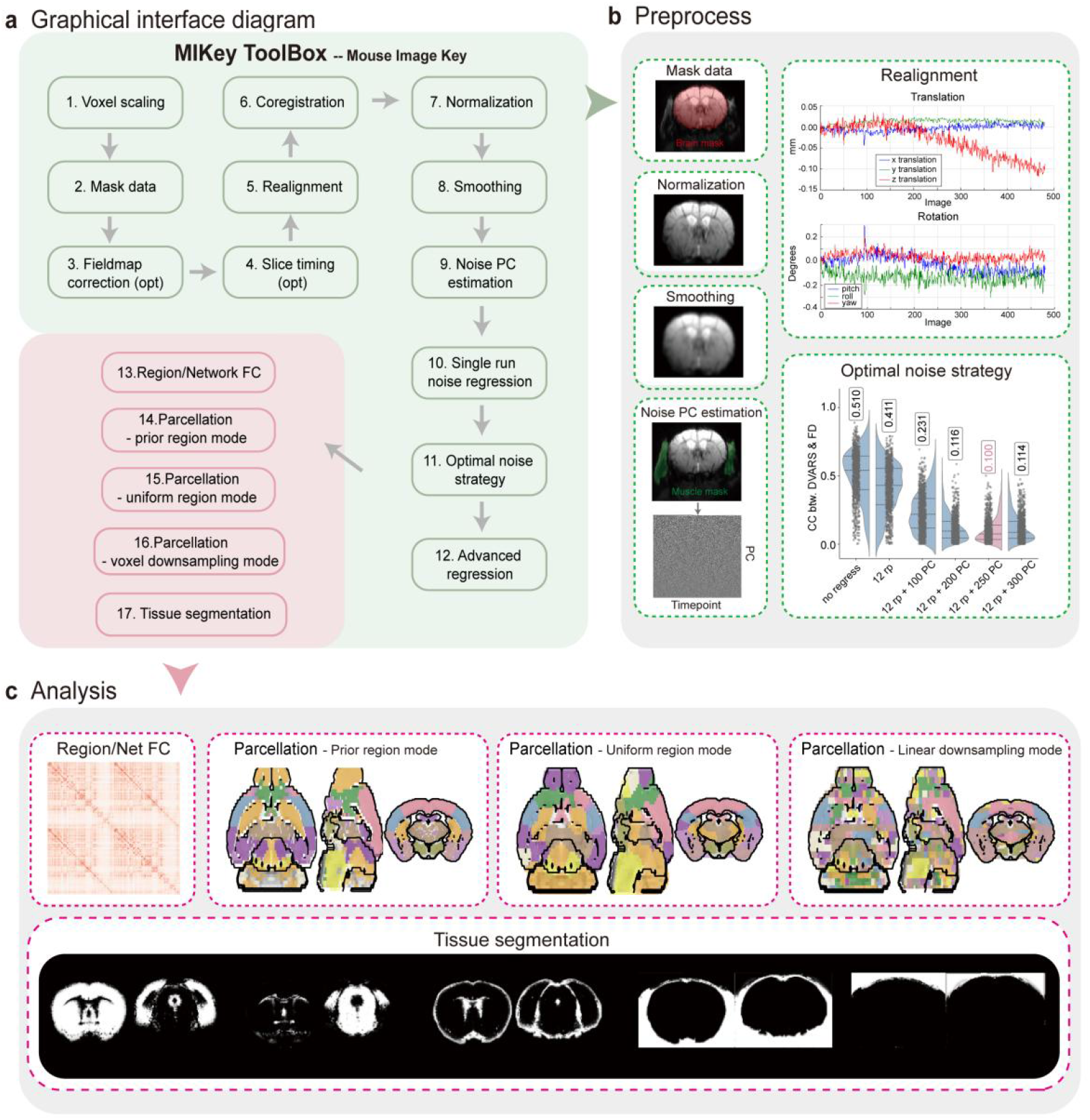
The MIKey Toolbox: an integrated platform for mouse fMRI from preprocessing to individualized parcellation with embedded AMCP resources. **a**, Graphical interface of MIKey, a MATLAB/SPM12-based open-source toolbox that integrates AMCP resources, including tissue probability maps and functional network atlases, with workflows for mouse fMRI preprocessing and analysis. **b**, Preprocessing module for awake mouse fMRI, including data organization, spatial preprocessing and nuisance regression strategies designed to reduce motion-related and non-neural variance. **c**, Post-processing and individualized-analysis module, including extraction of regional or network-level time series, computation of FC matrices, automated tissue segmentation using AMCP tissue priors, and individualized functional parcellation at the whole-brain or user-defined ROI level. The individualized parcellation module also includes a voxel-downsampling mode, in which setting the downsampling factor to 1 enables voxel-level parcellation for datasets with sufficient tSNR. FC, functional connectivity; GM, gray matter; WM, white matter; CSF, cerebrospinal fluid; OT, other tissue; ROI, region of interest.

To maximize the utility and impact of the AMCP dataset and its derived resources, we developed the MIKey Toolbox, a comprehensive open-source MATLAB-based software platform. We note that MIKey is not intended to replace existing rodent fMRI platforms such as RABIES^40^. Rather, the two tools emphasize partially different use cases: RABIES provides a standardized framework for rodent fMRI preprocessing, confound correction, and quality control, whereas MIKey was developed as a complementary MATLAB/SPM-based workflow that integrates the AMCP-specific templates, atlas resources, and individualized parcellation framework into a user-friendly end-to-end analysis environment. In parallel with earlier multi-centre mouse rsfMRI comparisons and ongoing open-science efforts such as AwakeRodent, this positioning is intended to improve interoperability while also highlighting the specific analytical value of AMCP-derived resources.

The preprocessing module of MIKey guides users from data organization to denoised time series, including slice-timing correction, motion realignment, spatial normalization to Allen CCFv3 space and nuisance regression (Fig. 9b). Following the artifact-reduction criterion used in AMCP, the toolbox can vary the number of non-brain PCs in strategies such as 6 rp + 6 Δrp + N PCs, allowing users to adapt denoising to dataset-specific motion and non-brain signal characteristics.

The post-processing module implements the individualized parcellation framework (Fig. 9c). In addition to the prior-label mode used in the main analysis, MIKey supports spatial k-means unit definition and linear downsampling, including voxel-wise units for datasets with sufficiently high tSNR. This allows users to choose a unit-definition strategy according to data quality and scientific objective. MIKey also supports network parcellation within user-defined regions of interest, enabling focused analyses of subsystem organization in structures such as hippocampal or thalamic subdivisions.

By packaging AMCP templates, atlases, preprocessing steps and individualized parcellation into a graphical interface (Supplementary Fig. 12), MIKey lowers the barrier to applying AMCP resources and facilitates replication and extension of awake mouse connectomics analyses.

## Discussion

AMCP establishes a population framework for awake mouse connectomics by integrating a large awake rsfMRI cohort, a normative network scaffold, structured individual variability, empirical data-quantity estimates and reusable analysis resources. This framework moves mouse fMRI beyond small-cohort feasibility studies toward a regime in which group architecture, individual differences and behaviorally relevant network features can be evaluated reproducibly.

A prerequisite for this framework is imaging the mouse brain in a state that is closer to the one used in human neuroimaging. Anesthesia shifts the brain into a state fundamentally different from physiological wakefulness and alters multiple processes that shape the BOLD signal^22^. Rather than preserving the awake relationship between neural activity and hemodynamics, anesthesia markedly changes neurovascular coupling. In mouse visual cortex, hemodynamic responses during wakefulness were reported to be substantially larger and faster than those measured under either isoflurane or urethane anesthesia, with these differences attributed primarily to altered neurovascular coupling rather than to comparably large changes in spiking activity^41^. At the systems level, awake and anesthetized mice also exhibit distinct large-scale resting-state organization, including stronger interregional communication, functional anti-coordination between visual-auditory and default-mode network regions, and state-dependent reconfiguration of rsfMRI dynamics in the awake brain^42^. Because human neuroimaging is performed predominantly in the awake state, awake mouse fMRI provides a more appropriate framework for mechanistic and translational comparisons across species.

Methodological choice remains a central issue for mouse functional network mapping. Human neuroimaging offers multiple parcellation strategies, and atlas choice can alter individual-difference associations^11^. In mouse fMRI, GICA remains common^24–27^, but model order and component stability are often difficult to evaluate systematically in small datasets. Our direct benchmark showed that, in the present awake dataset, the clustering-derived atlas had higher repeated-run reproducibility than conventional GICA (Fig. 5). This does not imply that ICA is intrinsically unsuitable. Rather, it supports clustering as a stable operating solution for the AMCP scaffold, while the released GICA solutions and the public AMCP dataset provide complementary references against which future methods can be compared.

Several lower-signal and boundary-adjacent regions require more cautious interpretation than the core large-scale network structure. Voxels with lower signal intensity and tSNR, particularly at cortical margins and in some ventral regions (Extended Data Fig. 8), tended to be assigned to the olfactory-limbic network (OLN). Reduced signal at these locations may reflect partial-volume effects, susceptibility-related signal loss at the tissue-CSF interface, receive-coil sensitivity profiles and intrinsic tissue properties such as lower cellular density in superficial cortical layers. These factors make OLN boundaries less certain than the central scaffold.

However, lower tSNR alone does not imply that these voxels should be excluded from atlas construction. In the 20-component GICA solution, voxels assigned to the OLN in the clustering-based atlas were not absorbed into diffuse noise-like components, but were represented by anatomically plausible and highly stable components, including a main olfactory bulb network (ICASSO IQ = 0.970, Fig. 5f) and an amygdalar network (ICASSO IQ = 0.974, Fig. 5f). This independent decomposition suggests that at least part of the OLN captures coherent olfactory-limbic organization, although the clustering solution groups these lower-signal and boundary-adjacent territories into a broader network. We therefore retained low-tSNR voxels to preserve whole-brain anatomical completeness and avoid an arbitrary post hoc exclusion threshold. Excluding them could systematically remove ventral, superficial or intrinsically low-signal territories and produce an atlas that no longer covers the whole brain. This concern parallels the caution applied to the human Yeo-7 Limbic Network, where orbitofrontal and ventral temporal regions are also affected by signal loss^43,44^. The OLN should therefore be treated as a lower-confidence but anatomically informative network, rather than as evidence that low-tSNR territories can be ignored.

The individualized parcellation framework is intentionally group-guided. Prior-defined anatomical units and a group-derived template improve convergence and cross-subject interpretability, but they limit fully unconstrained voxel-level boundary refinement. Future work should compare this strategy with matched bottom-up methods to test whether greater individual freedom improves behavioral prediction or boundary recovery.

A central biological implication of AMCP is that inter-individual variability in mouse functional networks is structured, measurable and behaviorally informative, rather than being incidental noise. These differences emerge within the constraints of a shared network scaffold, with local variation in network boundaries and connectivity contributing to behavioral prediction. In this respect, AMCP extends prior awake-mouse work on individual variability and fingerprinting by shifting the emphasis from the presence of individual functional signatures to their whole-brain network-level organization, cross-modal plausibility and behavioral relevance. This brings mouse connectomics closer to human population neuroscience, where individual differences in functional-connectivity architecture and cortical network topography are reproducible and predictive of cognition and behavior^45,46^, while preserving the advantage of future circuit-level perturbation and causal validation.

A key qualification is that observed inter-individual variability contains both stable between-subject differences and within-subject fluctuation. We did not acquire concurrent physiological or behavioral state measures during rsfMRI, such as respiration, heart rate or pupil diameter. Nevertheless, scan-half FCV similarity, half-scan scaling curves and repeated-measures reliability analyses all indicate a substantial stable component. We therefore interpret FCV as structured observed variability with an important subject-specific component, while acknowledging residual state and habituation effects.

The population scale of AMCP also enabled empirical estimation of data requirements for awake mouse connectomics. Previous human fMRI studies have shown that the amount of data required for stable estimation depends strongly on the analytical target. Stable estimation of large resting-state effective-connectivity effects, task-fMRI brain-behaviour correlations and individual-difference measures often requires sample sizes well beyond the conventional small-cohort regime. For example, Ma et al.^47^ reported that large DCM-based effective-connectivity effects approached a plateau at approximately 70 participants, whereas Grady et al.^48^ found that roughly 80 or more participants were needed to obtain stable estimates of task-fMRI brain-behaviour correlation magnitude. Awake mouse fMRI has lacked an analogous empirical foundation, largely because available public datasets have been too small to characterize how connectome estimates approach stability as a function of sample size and scan duration. By establishing the AMCP cohort, we were able to extend this type of marginal-gain analysis to awake mouse rsfMRI.

A central implication of this analysis is that there is no single sample-size threshold for mouse connectomics. Instead, the relevant benchmark depends on the estimand. The FCV curve is most appropriate for studies that aim to quantify inter-individual variability, variability axes or heterogeneity associated with genotype, treatment or phenotype. These questions require stable estimation of how connectivity varies across animals and therefore impose substantially higher data requirements. By contrast, the mean FC curve is more appropriate for studies focused on the group-average connectome or on group-level differences in mean connectivity. Because averaging across animals suppresses individual-specific variation, mean FC approached the full-sample reference with fewer animals than FCV. Thus, the FCV and mean FC curves should be viewed as design tools for different scientific questions, rather than as alternative estimates of a single universal threshold. The curves also clarify that longer scans can improve precision when sample size is constrained, especially for FCV or other variability-sensitive measures, but they are not a substitute for larger cohorts when the goal is to characterize population heterogeneity. They complement formal power analysis by quantifying the reliability of the connectomic quantity that enters downstream hypothesis testing.

The projection of independent awake mouse rsfMRI datasets onto the AMCP-derived curves provides a useful contextual comparison, but it also defines the limits of their transferability. The independent AMCP validation cohort fell slightly below the similarity expected from AMCP subsampling at matched data quantities, and the two public datasets fell further below the AMCP marginal-gain range. These observations indicate that the curves capture protocol-specific relationships between data quantity and connectome precision, rather than externally validated universal thresholds. Absolute values will vary with scanner hardware, coil sensitivity, preprocessing choices, motion control, network definition and biological effect size. The main contribution is therefore a quantitative framework for deciding whether a dataset is better suited for estimating mean connectivity, population-level variability or both, while requiring conservative interpretation when acquisition or data-quality conditions differ from AMCP.

The sex-balanced AMCP design enabled exploratory analyses of sexual dimorphism. We observed sex-associated differences in cortical GM, subcortical GM, WM and CSF volumes (Fig. 2e), together with localized diffusion differences (Extended Data Fig. 4f). We did not use Keigo-based segmentations as a downstream biological benchmark because visible tissue misclassifications in that framework could propagate segmentation error into biological inference. The small sex-related difference in connectional-axis slope (Supplementary Fig. 7d,e) should be treated as hypothesis-generating. The analysis was not preregistered, and independent sex-balanced awake mouse datasets are not yet large enough to support replication.

Several limitations define the boundary of the present resource. The awake head-fixed preparation remains a controlled state distinct from freely behaving animals. The 9.4T scanner and 4-channel surface coil produced lower SNR in ventral regions, which affects confidence in ventral and boundary-sensitive networks. The atlas was derived with one clustering framework, and its correspondence with cytoarchitectonic or molecular boundaries requires further validation. The optimized non-brain PC regression is intentionally aggressive; although visual-stimulation and preprocessing-sensitivity analyses support preservation of expected BOLD responses and large-scale network topology, FD-DVARS minimization is an artifact-reduction criterion and cannot by itself establish neural validity. Finally, because AMCP is Allen-space based whereas many rodent fMRI workflows use DSURQE, we will release DSURQE-space tissue maps and functional atlases to support interoperability.

Future extensions should test generalizability across strains, developmental stages, aging, disease models and multi-center acquisitions. AMCP also creates a basis for integrating awake functional networks with DTI, AMBA anatomy and mouse transcriptomic resources, enabling hypotheses about molecular and circuit-level mechanisms of macroscale network organization to be tested experimentally. AMCP should therefore be viewed as both a substantial resource and an initial step. Its most robust contributions are the large awake mouse dataset, tissue probability maps, functional atlas, individualized parcellation framework and empirical design curves. Numerical scaling thresholds and sex-related FCV observations remain protocol-specific and hypothesis-generating until validated across broader datasets. By enabling reproducible connectomics in a species amenable to circuit manipulation, AMCP provides a practical basis for linking cellular mechanisms to large-scale brain network organization.

## Supporting information

All Extended Figures

All Supplementary Figures

## Materials and Methods

### Animals

In total, 212 C57BL/6N mice, 8 weeks of age at the start of the experiments, were used as the main AMCP dataset. An additional validation cohort of 19 C57BL/6N mice, including 11 males, was acquired independently under the same experimental conditions. A smaller cohort of mice from three additional strains (BALB/cBy, C3H/He, and DBA/2) for template validation were also acquired and maintained under identical conditions. All mice were first purchased at 7 weeks of age from Beijing Vital River Laboratory Animal Technology Co., Ltd, and then acclimated to the laboratory environment for one week prior to experimentation. They were group-housed by sex in individually ventilated cages, with no more than six animals per cage, ensuring sufficient nesting material and activity space. The housing conditions included a 12-hour light/dark cycle, controlled temperature and humidity, and ad libitum access to food and water. All procedures were approved by the Institutional Animal Care and Use Committee of the Beijing Normal University.

### Headpost surgery

A custom-made MRI-compatible head holder was implanted over the mouse skull for head fixation during awake imaging (Fig. 1a). Under isoflurane anesthesia (induction at 5% and maintenance at 1-1.5%), the mouse head was fixed on a stereotaxic apparatus, and body temperature was maintained using a heating pad. The fur on the head was shaved, and the skin was disinfected alternately with alcohol and iodophor, while eye ointment was applied to protect the eyes. An incision was made along the midline of the scalp to expose the skull, and then the scalp over the skull as well as muscle covering the skull posterior to the lambda point were removed. The periosteum was removed using 3% hydrogen peroxide to reduce susceptibility artifacts. Afterwards, the exposed skull was cleaned with saline. Any fluid on the skull was then absorbed with dust-free paper, and a small amount of tissue adhesive was applied to bond the surrounding scalp to the exposed skull. Also, a thin layer of self-etch adhesive (3M ESPE Adper Easy one) was evenly spread over the skull, followed by light curing. To ensure long-term stable fixation, the light-curing flowable dental resin was employed and cured with blue light for 20 seconds to secure the head holder to the skull. Finally, we optimized the previous setup^29,32,33^ by applying a thick smooth layer of transparent dental cement over the remaining exposed skull area, which facilitates the observation of hemorrhage and tissue proliferation around the mouse head. To prevent postoperative infection, the mouse was injected with meloxicam 0.5 mg kg^−1^ (Baoding sunlight herb medicament, CN) subcutaneously for seven consecutive days post-surgery.

### Habituation

After one week of recovery from head holder surgery, the mice were habituated to acclimate immobilization and scanning sound for 14 days to minimize head movement and stress during awake fMRI scanning (Fig. 1a). Guided by an established habituation paradigm, the mice were secured on the animal bed and exposed to recorded MRI scanning noises. Over the first 7 days, MRI noise intensity and duration were gradually increased. From days 8 to 14, the mice experienced the maximum sound intensity (110 dB) and duration (120 min). The detailed habituation schedule is shown in Table 1. No rewards were given throughout the habituation.

**Table 1.** Summary of habituation schedules for awake mouse fMRI.

|  | Day 1 | Day 2 | Day 3-4 | Day 5-6 | Day 7 | Day 8-14 |
|---|---|---|---|---|---|---|
| Duration of habituation | 60 min | 75 min | 90 min | 100 min | 110 min | 120 min |
| Acoustic noise | - | 90 db | 95 db | 100 db | 105 db | 110 db |

Note: “-” means absence.

### MRI data acquisition

All MRI scans were performed at Changping Laboratory on a 9.4 Tesla scanner (uMR, United Imaging Life Science Instrument, Wuhan, China) with an 86-mm diameter volume transmission coil and a 4-channel mouse head surface receiver coil. Prior to scanning, each mouse was secured in a custom–made MRI–compatible animal bed, and a thick layer of Kwik–cast was smoothly applied over the skull to minimize distortion caused by air. High resolution T2–weighted structural imaging was performed with the following parameters: TR = 3000 ms, TE = 47.36 ms, slice thickness = 0.5 mm, field of view = 20 × 20 mm^2^, matrix size = 208 × 208, flip angle = 90°, RARE factor = 15, averages = 2, 30 coronal slices (voxel size = 0.1 × 0.1 × 0.5 mm³). For functional MRI, we first acquired B0 field map (phase and magnitude images) and a pair of gradient–echo echo–planar imaging (EPI) images with opposite phase–encoding directions (anterior-posterior and posterior-anterior; 5 volumes each) to correct for susceptibility–induced geometric distortions and signal loss. Subsequently, four resting-state fMRI runs were acquired using a multi-band gradient-echo EPI sequence (multi-band acceleration factor = 2), and the imaging parameters: TR = 1000 ms, TE = 15.9 ms, slice thickness = 0.5 mm, field of view = 20 mm × 20 mm^2^, matrix size = 132 × 132, bandwidth = 3100 Hz/pixel, flip angle = 55°, 28 coronal slices (voxel size = 0.15 × 0.15 × 0.5 mm). The first 5 volumes of the fMRI run were acquired as dummy scans to allow for signal stabilization and each run lasted about 8.09 min with 480 useful volumes. Finally, DTI-EPI data were acquired using a multi-shot diffusion-weighted EPI sequence to minimize the geometric distortions. The following parameters were applied: TR = 4000 ms, TE = 15 ms, slice thickness = 0.2 mm, field of view = 19 × 11 mm^2^, matrix size = 96 × 55, flip angle = 55°, 68 coronal slices (voxel size 0.2 × 0.2 × 0.2 mm^3^). The diffusion scheme comprised 33 volumes: one non-diffusion-weighted volume (b = 0 s/mm²) and 32 diffusion–weighted directions (16 each at b = 800 and 1600 s/mm²). Total acquisition time for the DTI scan was 32.55 minutes.

### Corticosterone validation

To validate physiological adaptation to the awake imaging preparation, an independent cohort of 19 mice underwent the same head-post implantation surgery, 14-day habituation protocol and MRI scanning procedure as used in the main AMCP cohort. Body weight was recorded longitudinally across the training and scanning period. Repeated tail-tip blood samples were collected from the same mice 1 day before habituation (D-1), on habituation days 1, 7 and 14, and after the first MRI scanning session for plasma corticosterone measurement. Except for the post-MRI sample, all blood samples were collected between 11:00 and 14:00 to minimize circadian effects. For D1, D7 and D14, blood was collected immediately after the habituation session. Each sampling procedure was completed within 2 min to reduce handling-induced acute stress.

Before baseline sampling at D-1, mice were habituated to experimenter handling for 10 min per day for 3 consecutive days and were then gently restrained by hand for blood collection. During habituation and after MRI scanning, mice were fixed on the MRI scanning bed for blood collection. The tail was disinfected with 75% ethanol, and approximately 0.5-1 mm of the tail tip was clipped using sterile scissors. Blood was collected into sodium-heparin tubes (HY-17567A, MedChemExpress), centrifuged at 10,000 rpm for 10 min at 4°C, and plasma was stored at −80°C until analysis. Plasma corticosterone was quantified using an ELISA kit (HB493-Mu, Shanghai Hengyuan Biological Technology) according to the manufacturer’s instructions, with absorbance measured at 450 nm using a multimode microplate reader (Cytation C10, Agilent Technologies). Pairwise comparisons between time points were performed using two-sided paired t-tests, with Holm-Bonferroni correction applied across planned comparisons.

### Behavior testing

All mice underwent a battery of four freely moving behavioral tests, performed within one week following MRI acquisition to ensure state proximity. The assays included: the open field test, the novel object recognition test, the spontaneous alternation test in the Y-maze, and the novel arm preference test in the Y-maze.

The Y-maze apparatus consisted of three identical arms (each 25 cm long × 6 cm wide × 14 cm high) arranged at 120-degree angles. A single protocol, comprising three sequential phases, was used to assess both spontaneous alternation and novel arm preference. In the spontaneous phase, mice explored all three accessible arms (A, B, and C) freely for 10 minutes to assess baseline alternation behavior. For the familiar phase, one arm was blocked, restricting exploration to the remaining two arms for 5 minutes. After a 120-minute inter-trial interval, during the recall/novelty phase, the blockade was removed, allowing the mouse to explore all three arms for 5 minutes to assess memory and preference for the previously unavailable (novel) arm.

The open field test was conducted in a 40 cm × 40 cm arena. The protocol involved three phases: a 10-minute adaptation phase in the empty arena, a 5-minute familiarization phase with two identical objects placed symmetrically, and a final 5-minute testing phase conducted after a 1-hour interval, during which one familiar object was replaced with a novel object distinct in visual and tactile properties.

Between trials, all apparatuses and objects were cleaned with 70% ethanol and dried to remove olfactory cues. Behavioral sessions were video recorded, and animal trajectories were automatically tracked using EthoVision XT (<u>Noldus;</u> https://www.noldus.com.cn/ethovision-xt/) (Supplementary Fig. 10a,b). Extracted metrics included total distance traveled, movement velocity, arm entries in the Y-maze, and object exploration time, defined as the mouse’s head being within 2 cm of an object with clear orientation or contact. In addition, LabGym^49^ was employed for AI-based identification and quantification of specific ethological patterns, such as grooming, rearing, head scanning (Supplementary Fig. 10c). This process yielded 1047 primary behavioral measures per subject. We additionally derived metrics that characterize relative exploration behaviors, such as (i) proportion of time exploring a novel object relative to total object exploration, and (ii) relative frequency of novel object exploration. In total, this resulted in 1054 behavioral metrics. A complete listing of these metrics is provided in Supplementary Table 1.

### BOLD data preprocessing

BOLD data were preprocessed using an established pipeline optimized for awake mouse fMRI^29,32,33,50^. To ensure compatibility with SPM12, BOLD and T2-weighted images were isotropically rescaled by a factor of 20. BOLD images were first corrected for susceptibility-induced distortions along the phase-encoding direction using FSL topup. For each subject, whole-brain masks for BOLD and T2-weighted images were generated using MouseBrainExtractor followed by manual refinement in ITK-SNAP. Subsequent preprocessing included field-map correction, rigid-body realignment, and co-registration of each subject’s high-resolution T2w image to the mean BOLD-EPI image. The co-registered T2w image was nonlinearly normalized to the Allen CCFv3 space using an Allen-referenced MRI-compatible volumetric template, and the resulting deformation fields were applied to all native-space EPI images. Data were then spatially smoothed with a Gaussian kernel of 6 mm FWHM in the 20-fold rescaled space, corresponding to 0.3 mm FWHM in native mouse space, approximately twice the in-plane BOLD voxel size.

Framewise displacement (FD) was calculated from the six SPM realignment parameters (6 rp) estimated from the raw BOLD data before denoising or downstream preprocessing^51^. For each volume, temporal derivatives of the three translational and three rotational parameters were computed. Rotational parameters were converted to displacement by multiplying by an assumed mouse brain radius of 5 mm, following Power et al.^51^ FD was then defined as the Euclidean norm of the six differential displacement terms:

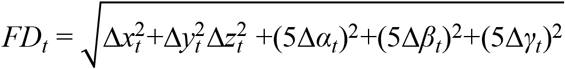

The same preprocessing, registration and FD estimation procedures were applied to the AMCP dataset and all publicly available comparison datasets. No animals, runs or volumes were excluded from these quality-control comparisons.

To mitigate non-neural physiological and motion-related confounds inherent to awake imaging, nuisance regression included the six rigid-body motion parameters and their first-order temporal derivatives, together with principal components derived from a non-brain tissue mask dominated by extracranial signal sources such as musculature. DVARS was computed as the root-mean-square temporal derivative of voxel-wise BOLD signal across brain voxels^51^. Specifically, for each time point, the signal difference between the current and preceding volume was calculated at every brain voxel, squared, averaged across voxels and square-root transformed. DVARS was recomputed after each denoising strategy, and FD-DVARS coupling was quantified as the Pearson correlation between the raw-data FD time series and the DVARS time series from the corresponding denoised data. The number of non-brain principal components was selected by minimizing FD-DVARS coupling across candidate denoising strategies. This criterion was used as an artifact-reduction objective, not as standalone evidence that neural BOLD variance was preserved. The optimized strategy used in the main analysis regressed 250 non-brain PCs in addition to the 12 motion-related regressors (6 rp + 6 Delta rp + 250 PCs; Fig. 1e). Thus, for each run with 480 retained volumes, the nuisance model included 262 regressors, leaving a nominal 218 residual temporal degrees of freedom before subsequent detrending and band-pass filtering; the effective degrees of freedom after filtering are lower and depend on temporal autocorrelation. The residual time series were then linearly detrended and band-pass filtered within 0.01-0.1 Hz. Biological preservation after this regression strategy was evaluated separately using visual-stimulation fMRI and atlas-robustness analyses.

Functional data quality was quantified using temporal and spatial signal-to-noise ratios. Temporal SNR was computed voxel-wise as the temporal mean divided by the temporal standard deviation of each voxel’s signal. Spatial SNR was estimated using a region-based approach. A background noise ROI was defined from voxels outside the brain mask with the lowest 0.5% signal intensity. The brain was parcellated into 1,000 approximately equal-volume spatial regions, and for each time point the mean signal within each region was divided by the standard deviation of the background noise ROI. Region-wise sSNR values were assigned back to constituent voxels, averaged across the session and spatially smoothed to generate whole-brain sSNR maps. Cross-dataset comparisons of sSNR and tSNR were corrected for voxel-size differences because larger voxels tend to yield higher SNR at the expense of spatial resolution. Specifically, measured SNR values were normalized by the square root of voxel volume:

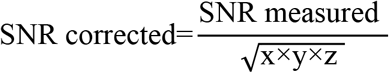

where x×y×z denotes voxel volume in mm^3^. This correction was applied uniformly across all datasets to facilitate comparison across different spatial resolutions.

### Visual-stimulation fMRI validation experiment

To validate whether non-brain PC regression preserved neural BOLD responses, we performed an independent visual-stimulation fMRI experiment in three awake mice using the same head-fixation setup, BOLD-EPI sequence and preprocessing framework as in the main AMCP dataset. Bilateral visual stimulation was delivered using a 437-nm laser through optical fibers positioned approximately 1 cm from each eye. The stimulation paradigm consisted of 20-s stimulation blocks followed by 40-s inter-stimulation intervals, repeated for 30 trials. Stimulation was delivered at 10 Hz with a 20% duty cycle and 1 mW light power. The data were analyzed under three denoising strategies: “12 rp” regressors only, “12 rp + 20 PCs”, and “12 rp + 250 PCs”. Voxel-wise stimulation-evoked activation maps were thresholded at FDR-corrected P < 0.05 with a minimum cluster size of 20 voxels. Activation patterns were compared across denoising strategies to determine whether non-brain PC regression preserved responses in the expected visual pathway, including the primary visual cortex, superior colliculus and lateral geniculate nucleus, while reducing nonspecific activation outside this circuit.

### Construction of AMCP tissue probability maps

For each high-resolution T2-weighted image in the standard Allen CCFv3 space, an initial tissue segmentation was performed using the Keigo 5-component probabilistic template, which delineates GM, WM, CSF, OT and air. The resulting segmentation maps from all mice were then employed to generate a study-specific, high-fidelity template through the SPM12 DARTEL (Diffeomorphic Anatomical Registration Through Exponentiated Lie Algebra) tool. This process yielded the AMCP Tissue Probability Maps. We intentionally constructed these tissue priors from in vivo data acquired in the same cohort used for the awake rsfMRI and DTI analyses. Although ex vivo MRI can provide higher anatomical resolution^52^, our goal here was to establish a subject-matched multimodal in vivo reference framework so that tissue segmentation, functional atlas construction, structural-connectivity estimation, and downstream structure-function analyses could all be performed within the same animals and the same imaging space. Finally, to maximize anatomical precision, each individual’s T2w image was resegmented using these newly created AMCP tissue priors. In the tissue-template analysis, the direct probabilistic-prior comparison was between the Keigo and AMCP frameworks, because both provide probabilistic tissue priors. DSURQE was not used as an equivalent benchmark here, as it provides anatomical labels rather than directly comparable probabilistic GM/WM/CSF priors. The AMCP tissue priors were therefore generated and evaluated within the same Allen-based coordinate system used throughout the study, ensuring consistency between tissue segmentation and all downstream structural and functional analyses.

### DTI data processing

The DTI data preprocessing pipeline was implemented as follows. Prior to tensor estimation, non-brain tissue was manually removed via skull stripping using ITK-SNAP. Subsequent preprocessing, executed within the MRtrix3 framework (https://www.mrtrix.org/), included Marchenko-Pastur principal component analysis (MP–PCA) denoising^53^, Gibbs unringing^54^, combined head motion and eddy current correction^55^, and B1 bias field correction^56^ using the *dwidenoise*, *mrdegibbs*, *dwifslpreproc*, and *dwibiascorrect* functions, respectively. From the preprocessed data, fractional anisotropy (FA), mean diffusivity (MD), axial diffusivity (AD), and radial diffusivity (RD) maps were derived using MRtrix3’s dwi2tensor and *tensor2metric* functions^57^.

For spatial normalization, the precomputed nonlinear transformation field from each subject’s T2–weighted image to the Allen CCFv3 standard space was applied to all individual microstructural maps (FA, MD, AD, RD). Population–averaged templates for each metric were subsequently generated from the normalized maps of all mice using the *antsMultivariateTemplateConstruction2* script from Advanced Normalization Tools (ANTs). A group mean white matter skeleton was created by applying an FA threshold of 0.2 to the group mean FA template via FSL’s Tract–Based Spatial Statistics (TBSS) pipeline (*tbss_3_postreg*)^58^. Voxel–wise analysis on the skeletonized data was performed using FSL’s *randomise* tool with 1000 permutations to assess sexual dimorphism, with significance evaluated via Threshold–Free Cluster Enhancement (TFCE) at a family–wise error (FWE) corrected threshold of p < 0.05.

Whole–brain probabilistic tractography was performed to construct structural connectomes. Fiber orientation distributions were estimated in each voxel using FSL’s *BedpostX*^59^. For each subject, 5000 streamlines were initiated per voxel in the diffusion space. To minimize false–positive connections, a consistency threshold was applied, excluding voxels traversed by fewer than 2 streamlines. This produced an individual voxel–wise structural connectivity (SC) matrix. The AMCP 11–network atlas and the AMBA (e.g., 428 regions) were then warped from standard space into each subject’s native diffusion space to derive network–level and region–level SC matrices, respectively.

### Construction of group-level functional network atlas

To address the lack of a comprehensive whole-brain functional network atlas for awake mice, we adapted the connectivity-profile clustering logic introduced by Yeo et al.^7^ and optimized it for high-dimensional whole-brain mouse fMRI data (Fig. 3a). This adaptation used k-means clustering rather than the von Mises-Fisher mixture-model clustering used in the original Yeo framework following Lashkari et al.^60^; the resulting networks should therefore be interpreted as stable k-means partitions of mouse whole-brain connectivity profiles.

The procedure consisted of the following key stages. First, to define the feature space for clustering, we generated multiple prior parcellations. These included anatomical partitions based on the AMBA at several levels of granularity (214, 428, 604, and 1184 regions) and a series of spatially uniform parcellations (750, 1000, 1200, and 1500 regions). The Allen-based schemes were selected to sample coarse-to-fine anatomical scales within the atlas hierarchy, with 428 representing the bilateral version of the 214-region set, whereas the uniform schemes were included to parallel the equal-sized regional partitioning strategy used in the original Yeo framework. For each scanning run, we computed a voxel-to-region functional connectivity matrix, correlating the time series of each of the 149,112 whole-brain voxels with the average time series of every region within a given parcellation. To focus on the strongest associations, each correlation matrix was sparsified by retaining only the top 10% of connections.

Next, we performed a systematic parameter search to identify the most stable clustering solution. For each combination of region count (R) and target network number (N), subjects were randomly divided into two non-overlapping subsets, and all runs from a given subject were retained within the same subset; this subject-level split-half procedure was repeated 10 times. A binarized voxel-to-region connectivity matrix was constructed independently for each subset, and k-means clustering was applied separately to the two subset-specific matrices to group voxels into N candidate networks. The consistency between the resulting partitions was quantified using the Dice similarity coefficient, with 1 minus Dice defined as the instability metric.

The optimal parameter set was identified at the inflection point of the instability curve, corresponding to a local minimum that balanced granularity and robustness. Using these optimized parameters, k-means clustering was then performed on the full, unsplit dataset to generate the final group-level functional network atlas.

### Validation of AMCP functional network atlas

Having identified the optimal functional network solution based on minimal split-half instability, we conducted a series of validation analyses to comprehensively assess its robustness. Validation proceeded along two primary axes: methodological stability and biological reliability.

First, we evaluated the stability of the parcellation against common preprocessing variations. To test sensitivity to denoising strategy, we applied the optimal clustering parameters to datasets processed with three alternative nuisance regression models: one including GS regression, one including CSF signal regression, and one combining both GS and CSF regressors. The spatial similarity between networks derived from these strategies and our primary solution was quantified using the Dice coefficient. In parallel, we assessed the impact of the correlation threshold used to sparsify the voxel-to-region connectivity matrix prior to clustering. The optimal clustering was repeated on matrices binarized at thresholds ranging from 5% to 50% (retaining the top 5%, 20%, 30%, 40%, and 50% of edges). Network similarity across these thresholds was again quantified using the Dice coefficient. High Dice values across these tests would indicate that the derived network architecture is robust to specific methodological choices.

Second, we validated the biological reliability of the atlas by examining its alignment with underlying structural connectivity and behavioral output. We first confirmed the fundamental property of functional modularity: for each of the 11 networks, within-network FC strength was calculated and compared to its average between-network FC strength (i.e., its mean connectivity to all other networks) using paired-sample t-tests across all subjects. We then tested whether this functional modularity was supported by anatomy. A group-level SC matrix was generated from DTI-based tractography and normalized to control for global individual differences. Within-network and between-network SC strengths were computed for each network and compared via paired-sample t-tests. Finally, to link the atlas to organism-level function, we examined its behavioral relevance. We calculated the total FC strength of the SMN-defined as its average connectivity with all other networks-and tested its correlation with three core locomotion metrics extracted from the Y-maze spontaneous exploration phase: total movement distance, movement time, and average velocity.

### Group-based ICA

Group-level independent component analysis (GICA) was performed using the GIFT toolbox v4.0 to evaluate large-scale functional components derived from a conventional ICA framework. We examined a range of model orders and assessed component stability and spatial reproducibility across repeated scanning runs. For each model order, the Infomax algorithm was repeated 20 times with different initializations and bootstrapped data subsets using the ICASSO framework, and component stability was quantified for each independent component. Subject-specific spatial maps and time courses for each scanning run were then estimated using spatiotemporal regression.

To evaluate spatial reproducibility of GICA-derived components, we used both continuous-map correlation and threshold-based spatial overlap metrics. For continuous spatial correspondence, run-to-run similarity was quantified by Pearson correlation between subject-specific component maps from different runs without thresholding. Run-to-group similarity was quantified by Pearson correlation between each run-specific component map and the corresponding group-level component map, also without thresholding. For threshold-based overlap analysis, subject-specific run-level spatial maps were binarized at the 80th percentile of their intensity distribution. Pairwise run-to-run spatial overlap was then quantified using the Dice coefficient. Run-to-group Dice overlap was computed using the same 80th-percentile binarization procedure. These analyses were performed across GICA model orders to identify a representative solution with stable, spatially coherent components.

The 20-component solution was selected as the representative GICA model order for comparison with the clustering-derived AMCP atlas because it provided a practical balance between component stability, spatial reproducibility and anatomical interpretability. Lower-dimensional solutions yielded less differentiated components, whereas higher-dimensional solutions increasingly showed spatial fragmentation, unilateral maps or reduced ICASSO stability, including components with IQ values below 0.8. For visualization, component maps from the 20-component solution were converted to Z scores and thresholded at ∣Z∣>1.96. The thresholded maps were used only for display of spatial topography, whereas the continuous spatial-correlation analyses were performed on unthresholded component maps.

### Quantification of inter-individual variability in functional and structural connectivity

We quantified inter-individual variability in both functional (FCV) and structural connectivity (SCV) within a unified analytical framework adapted from established methodologies^5^. This approach ensured a consistent and comparable assessment of population heterogeneity across modalities.

For FCV analysis, regional BOLD time series were extracted using the AMBA (428 regions). A symmetric 428 × 428 functional connectivity matrix was computed per run via Pearson correlation and Fisher’s z-transformation for each mouse. The within-mouse and between-mouse variance components for each edge were estimated using a linear mixed-effects model, from which the intra-class correlation coefficient (ICC) was derived to formalize FCV as the proportion of total variance attributable to inter-individual differences, ICC = σ²_between / (σ²_between + σ²_within), where σ²_between and σ²_within denote the between- and within-subject variance, respectively. This metric therefore quantifies adjusted between-mouse FCV while controlling for intra-mouse fluctuations.

The SCV analysis followed a parallel computational framework. First, a consistency-based threshold was applied to each individual’s streamline-count matrix to reduce noise, retaining edges with low cross-subject coefficient of variation. For each thresholded matrix, communicability was computed to incorporate direct and indirect pathways, and SCV for each edge was defined as the mean absolute deviation of its communicability value across all subjects.

Finally, to facilitate a direct systems-level comparison, both edge-wise FCV and SCV matrices were summarized using our AMCP 11-network atlas. Regional values were mapped and averaged within and between networks, producing two analogous 11 × 11 symmetric matrices for FCV and SCV, where diagonal elements represent within-network variability and off-diagonal elements represent between-network variability.

### Repeated-measures reliability analysis of network-level FC terms

To quantify measurement reliability, we used the four repeated resting-state runs available for each subject. For each run, the ROI-level functional connectivity matrix was first summarized into 66 network-level FC terms based on the fixed AMCP 11-network assignment, including 11 within-network terms and 55 between-network terms. Within-network terms were defined as the mean FC across all ROI pairs belonging to the same network, whereas between-network terms were defined as the mean FC across all ROI pairs linking two different networks. Reliability across runs was quantified using intraclass correlation coefficients, reported as ICC(3,1) for single-run reliability and ICC(3,k) for the reliability of the averaged measure across repeated runs. To further characterize the variance structure of these measures, we also estimated between-subject and within-subject/error variance components for each network-level term using the same repeated-measures ANOVA framework.

### Scan-half stability analysis of FCV architecture and scaling

To test whether the main FCV architecture and scaling behavior were sensitive to gradual scan-time state changes, we repeated the FCV analysis separately for the first half (Runs 1-2) and second half (Runs 3-4) of the rsfMRI data. For each scan half, ROI-level FC matrices were computed using the same preprocessing and FCV-estimation framework as in the main analysis, and then summarized into network-level FCV matrices. Concordance between the two scan halves was quantified by correlating the corresponding network-level FCV matrices. We further repeated the sample-size analysis independently for the two scan halves by computing subset-derived FCV matrices and quantifying their similarity to the corresponding half-scan reference FCV matrix. This yielded separate sample- and time-dependent similarity profiles and marginal gain curves for Runs 1-2 and Runs 3-4, allowing direct comparison of scaling behavior across early and late portions of the scanning session.

### Individualized functional network parcellation

To derive individualized functional networks, we implemented a modified iterative parcellation algorithm based on the framework by Wang et al.^39^. This adaptation was designed to address the convergence challenges inherent in high-dimensional, voxel-wise mouse fMRI data. The procedure consisted of two main phases: data reduction/initialization and iterative optimization. To ensure stable algorithmic convergence, the high-dimensional whole-brain voxel space was first reduced to a set of representative functional units. In our primary analysis, this was achieved using a prior-label strategy, where voxel time series were averaged within each of the 214 anatomical regions defined by the AMBA, with each region’s average signal constituting one initial unit.

The network assignment for each unit was then refined through an iterative optimization process. In each iteration, a reference time course was first generated for every network. This reference was computed as a dynamically weighted combination of the current average time series of all units tentatively assigned to that network (the “core” signal) and the initial, atlas-based reference signal. Subsequently, each unit’s time series was correlated with all network reference signals, and the unit was tentatively reassigned to the network showing the highest correlation. A confidence value for this reassignment was calculated as the ratio of the largest to the second-largest correlation. The algorithm iterated until convergence criteria were met: specifically, until the percentage of units changing network assignment fell below a set threshold (change_thr = 0.1%) and the mean confidence ratio across all units exceeded a threshold (conf_thr = 2.5), or until a maximum number of iterations (i.e., 100) was reached.

The final output of this process was a stable, individualized functional parcellation for each subject. Network-level functional connectivity matrices were subsequently computed from the average time series of units assigned to the same network. This flexible framework also accommodates alternative unit-definition strategies, such as k-means clustering based on spatial coordinates or direct voxel-wise analysis for data with sufficiently high temporal signal-to-noise ratio.

### Composite network FC-behavior predictor

The composite network FC-behavior Predictor was constructed to link network functional connectivity to behavior. For each mouse, across four scan runs, network time-series data were extracted based on a group-level average atlas (AMCP 11-Network Atlas) and an individualized network parcellation, respectively. FC matrices were then generated by computing Pearson’s correlation for the time-series data from each run, applying Fisher’s Z-transform to the correlation coefficients, and then averaged across runs to yield one 11×11 FC matrix per subject. This yielded two types of FC matrices per subject: one reflecting the group-level atlas and another based on individual parcellation.

The upper-triangular elements of each symmetric FC matrix were retained, yielding 55 network-to-network edges per subject. Each behavioral outcome was modeled independently. To avoid information leakage, mass-univariate feature screening, FDR correction and feature scaling were performed within each outer training fold only, using training subjects alone.

Predictive modeling used ridge regression with nested cross-validation. The outer 10-fold split was performed at the subject level, so all runs from a mouse contributed only to either the training or test set. Within each outer training fold, a 5-fold inner cross-validation selected the regularization parameter lambda from a predefined logarithmic range. The final model, trained with the fold-specific feature set and selected lambda, was applied once to the held-out test subjects. Prediction performance was quantified as the correlation between predicted and observed values across all outer folds. Statistical significance was assessed using 1,000 label-permutation tests.

## Data and code availability

All data generated in this study, including neuroimaging and behavioral video data from 212 fully awake, sex-balanced C57BL mice, the AMCP atlas, templates, and MIKey toolbox, have been deposited in the Science Data Bank (DOI: 10.57760/sciencedb.35040) and are accessible through the AMCP resource website (http://amcp.org.cn/), which serves as a portal for data access, visualization, and download. All imaging and behavioral datasets are organized according to the standard Brain Imaging Data Structure (BIDS) format to facilitate community reuse and interoperability across analysis platforms. All resources will be publicly released without restriction upon acceptance of the manuscript. Source data are provided with this paper and contain all numerical values underlying the figures presented in the main manuscript.

For quality comparison, this study utilized the following publicly available awake C57-mouse fMRI datasets: ds003027 (https://openneuro.org/datasets/ds00 3027), ds003468 (https://openneuro.org/datasets/ds003468), ds001653 (https://openneuro.org/datasets/ds001653), ds002551 (https://openneuro.org/datasets/ds002551), ds001720 (https://openneuro.org/datasets/ds001720), and Gutierrez-Barragan datas et (https://data.mendeley.com/datasets/np2fx99hn6/2).

In addition to the Allen-space resources used in the present study (https://github.com/wangxiaoqun-lab/MIKey/tree/main/share_temp/AMCP_atlas/), we will also release DSURQE-space versions of the AMCP-derived tissue probability maps and functional network atlas to support interoperability with DSURQE-based preprocessing workflows (https://github.com/wangxiaoqun-lab/MIKey/tree/main/share_temp/AMCP_atlas/DS URQE_40_40_40).

## Acknowledgements

This work was supported by the National Natural Science Foundation of China (NSFC) (82501714, 32192411), the Science and Technology Innovation 2030-Brain Science and Brain-inspired Intelligence Project of China (2021ZD0200102, 2022ZD0211900, and 2022ZD0207700), CAS Project for Young Scientists in Basic Research (YSBR-097), the New Cornerstone Science Foundation, Changping Laboratory, the Fundamental Research Funds for the Central Universities.

## Author Contributions

X.W. conceived and supervised the project and designed the experiments. Y.Z., C.Z., X.H., W.W., Y.J., and Y.Y.J. performed the animal surgery under the guidance of Q.W. and Y.Y. R.T. carried out the animal training. R.T. and Y.Z. performed the MRI experiments. C.Z. and R.T. conducted the behavioral experiments. C.Z. and R.T. were responsible for animal husbandry. Y.Z. and R.T. analyzed the MRI and behavioral data. D.H. collected all data for the AMCP validation cohort, including MRI data, tail-vein corticosterone measurements, and animal body-weight records. X.W., Y.Z., Q.W., and S.Z. discussed and interpreted the results. Y.Z., and X.W. wrote the manuscript. All authors reviewed, edited, and approved the final manuscript.

---

**Supplementary Table 1.** 

## References

1. Finn, E. S., Poldrack, R. A. & Shine, J. M. Functional neuroimaging as a catalyst for integrated neuroscience. Nature 623, 263–273 (2023).

2. Van Essen, D. C. et al. The Human Connectome Project: a data acquisition perspective. Neuroimage 62, 2222–2231 (2012).

3. Glasser, M. F. et al. The Human Connectome Project’s neuroimaging approach. Nat Neurosci 19, 1175–1187 (2016).

4. Tong, C. et al. Multimodal analysis demonstrating the shaping of functional gradients in the marmoset brain. Nat Commun 13, 6584 (2022).

5. Yang, H. et al. Connectional axis of individual functional variability: Patterns, structural correlates, and relevance for development and cognition. Proc. Natl. Acad. Sci. U.S.A. 122, e2420228122 (2025).

6. Huang, S. et al. TransBrain: a computational framework for translating brain-wide phenotypes between humans and mice. Nat Methods 23, 426–437 (2026).

7. Yeo, B. T. T. et al. The organization of the human cerebral cortex estimated by intrinsic functional connectivity. Journal of neurophysiology 106, 1125–1165 (2011).

8. Schaefer, A. et al. Local-Global Parcellation of the Human Cerebral Cortex from Intrinsic Functional Connectivity MRI. Cereb Cortex 28, 3095–3114 (2018).

9. Power, J. D. et al. Functional network organization of the human brain. Neuron 72, 665–678 (2011).

10. Gordon, E. M. et al. Generation and Evaluation of a Cortical Area Parcellation from Resting-State Correlations. Cereb Cortex 26, 288–303 (2016).

11. Bryce, N. V. et al. Brain parcellation selection: An overlooked decision point with meaningful effects on individual differences in resting-state functional connectivity. Neuroimage 243, 118487 (2021).

12. Milham, M. P. et al. Assessment of the impact of shared brain imaging data on the scientific literature. Nat Commun 9, 2818 (2018).

13. Bressler, S. L. & Menon, V. Large-scale brain networks in cognition: emerging methods and principles. Trends Cogn Sci 14, 277–290 (2010).

14. Suárez, L. E., Markello, R. D., Betzel, R. F. & Misic, B. Linking Structure and Function in Macroscale Brain Networks. Trends Cogn Sci 24, 302–315 (2020).

15. Bassett, D. S. & Sporns, O. Network neuroscience. Nat Neurosci 20, 353–364 (2017).

16. Pagani, M., Gutierrez-Barragan, D., de Guzman, A. E., Xu, T. & Gozzi, A. Mapping and comparing fMRI connectivity networks across species. Commun Biol 6, 1238 (2023).

17. Gutierrez-Barragan, D., Ramirez, J. S. B., Panzeri, S., Xu, T. & Gozzi, A. Evolutionarily conserved fMRI network dynamics in the mouse, macaque, and human brain. Nat Commun 15, 8518 (2024).

18. Ganeshan, K. & Chawla, A. Warming the mouse to model human diseases. Nat Rev Endocrinol 13, 458–465 (2017).

19. Liang, Z. et al. Mapping the functional network of medial prefrontal cortex by combining optogenetics and fMRI in awake rats. NeuroImage 117, 114–123 (2015).

20. Shim, H.-J. et al. Mouse fMRI under ketamine and xylazine anesthesia: Robust contralateral somatosensory cortex activation in response to forepaw stimulation. NeuroImage 177, 30–44 (2018).

21. Franceschini, M. A. et al. The effect of different anesthetics on neurovascular coupling. NeuroImage 51, 1367–1377 (2010).

22. Gao, Y.-R. et al. Time to wake up: Studying neurovascular coupling and brain-wide circuit function in the un-anesthetized animal. Neuroimage 153, 382–398 (2017).

23. Hike, D. et al. High-resolution awake mouse fMRI at 14 tesla. Elife 13, RP95528 (2025).

24. Gezginer, I. et al. Concurrent optoacoustic tomography and magnetic resonance imaging of resting-state functional connectivity in the mouse brain. Nat Commun 15, 10791 (2024).

25. Grandjean, J., Zerbi, V., Balsters, J. H., Wenderoth, N. & Rudin, M. Structural Basis of Large-Scale Functional Connectivity in the Mouse. J Neurosci 37, 8092–8101 (2017).

26. Grandjean, J. et al. Common functional networks in the mouse brain revealed by multi-centre resting-state fMRI analysis. NeuroImage 205, 116278 (2020).

27. Zerbi, V., Grandjean, J., Rudin, M. & Wenderoth, N. Mapping the mouse brain with rs-fMRI: An optimized pipeline for functional network identification. Neuroimage 123, 11–21 (2015).

28. Chen, X. et al. Sensory evoked fMRI paradigms in awake mice. Neuroimage 204, 116242 (2020).

29. Zou, Y. et al. Cell-type-specific optogenetic fMRI on basal forebrain reveals functional network basis of behavioral preference. Neuron 112, 1342–1357.e6 (2024).

30. Yu, Y. et al. Sleep fMRI with simultaneous electrophysiology at 9.4 T in male mice. Nat Commun 14, 1651 (2023).

31. Han, Z. et al. Awake and behaving mouse fMRI during Go/No-Go task. Neuroimage 188, 733–742 (2019).

32. Xu, W. et al. A systematically optimized awake mouse fMRI paradigm. *bioRxiv* 2022.11.16.516376 (2022).

33. Tong, C. et al. Norepinephrine-mediated arousal fluctuations drive inverted U-shaped functional connectivity dynamics. Nat Commun 16, 11318 (2025).

34. Hikishima, K. et al. In vivo microscopic voxel-based morphometry with a brain template to characterize strain-specific structures in the mouse brain. Sci Rep 7, 85 (2017).

35. Dong, H.-M., Margulies, D. S., Zuo, X.-N. & Holmes, A. J. Shifting gradients of macroscale cortical organization mark the transition from childhood to adolescence. Proc Natl Acad Sci U S A 118, e2024448118 (2021).

36. Bergmann, E., Zur, G., Bershadsky, G. & Kahn, I. The Organization of Mouse and Human Cortico-Hippocampal Networks Estimated by Intrinsic Functional Connectivity. Cereb Cortex 26, 4497–4512 (2016).

37. Melozzi, F. et al. Individual structural features constrain the mouse functional connectome. Proc. Natl. Acad. Sci. U.S.A. 116, 26961–26969 (2019).

38. Bergmann, E., Gofman, X., Kavushansky, A. & Kahn, I. Individual variability in functional connectivity architecture of the mouse brain. Commun Biol 3, 738 (2020).

39. Wang, D. et al. Parcellating cortical functional networks in individuals. Nat Neurosci 18, 1853–1860 (2015).

40. Desrosiers-Grégoire, G., Devenyi, G. A., Grandjean, J. & Chakravarty, M. M. A standardized image processing and data quality platform for rodent fMRI. Nat Commun 15, 6708 (2024).

41. Pisauro, M. A., Dhruv, N. T., Carandini, M. & Benucci, A. Fast Hemodynamic Responses in the Visual Cortex of the Awake Mouse. J. Neurosci. 33, 18343–18351 (2013).

42. Ferenczi, E. A. et al. Prefrontal cortical regulation of brainwide circuit dynamics and reward-related behavior. Science 351, aac9698 (2016).

43. Huang, C.-C., Rolls, E. T., Feng, J. & Lin, C.-P. An extended Human Connectome Project multimodal parcellation atlas of the human cortex and subcortical areas. Brain Struct Funct 227, 763–778 (2022).

44. Wig, G. S., Laumann, T. O. & Petersen, S. E. An approach for parcellating human cortical areas using resting-state correlations. Neuroimage 93 **Pt** **2**, 276–291 (2014).

45. Finn, E. S. et al. Functional connectome fingerprinting: identifying individuals using patterns of brain connectivity. Nat Neurosci 18, 1664–1671 (2015).

46. Mueller, S. et al. Individual Variability in Functional Connectivity Architecture of the Human Brain. Neuron 77, 586–595 (2013).

47. Ma, L. et al. Effect of scanning duration and sample size on reliability in resting state fMRI dynamic causal modeling analysis. NeuroImage 292, 120604 (2024).

48. Grady, C. L., Rieck, J. R., Nichol, D., Rodrigue, K. M. & Kennedy, K. M. Influence of sample size and analytic approach on stability and interpretation of brain-behavior correlations in task-related fMRI data. Human Brain Mapping 42, 204–219 (2021).

49. Hu, Y., et al. *LabGym*: Quantification of user-defined animal behaviors using learning-based holistic assessment. Cell Reports Methods 3, 100415 (2023).

50. Zhang, C. et al. Integrating optogenetic fMRI and spatial transcriptomics to reveal circuit-specific gene signatures in fronto- and hippo-thalamic networks. Nat Commun 10.1038/s41467-026-71923-w (2026) doi:10.1038/s41467-026-71923-w.

51. Power, J. D., Barnes, K. A., Snyder, A. Z., Schlaggar, B. L. & Petersen, S. E. Spurious but systematic correlations in functional connectivity MRI networks arise from subject motion. NeuroImage 59, 2142–2154 (2012).

52. Lerch, J. P. et al. Wanted dead or alive? The tradeoff between in-vivo versus ex-vivo MR brain imaging in the mouse. Front. Neuroinform. 6, (2012).

53. Veraart, J. et al. Denoising of diffusion MRI using random matrix theory. Neuroimage 142, 394–406 (2016).

54. Kellner, E., Dhital, B., Kiselev, V. G. & Reisert, M. Gibbs-ringing artifact removal based on local subvoxel-shifts. Magn Reson Med 76, 1574–1581 (2016).

55. Andersson, J. L. R., Graham, M. S., Zsoldos, E. & Sotiropoulos, S. N. Incorporating outlier detection and replacement into a non-parametric framework for movement and distortion correction of diffusion MR images. NeuroImage 141, 556–572 (2016).

56. Tustison, N. J. et al. N4ITK: improved N3 bias correction. IEEE Trans Med Imaging 29, 1310–1320 (2010).

57. Tournier, J.-D. et al. MRtrix3: A fast, flexible and open software framework for medical image processing and visualisation. Neuroimage 202, 116137 (2019).

58. Smith, S. M. & Nichols, T. E. Threshold-free cluster enhancement: addressing problems of smoothing, threshold dependence and localisation in cluster inference. Neuroimage 44, 83–98 (2009).

59. Behrens, T. E. J., Berg, H. J., Jbabdi, S., Rushworth, M. F. S. & Woolrich, M. W. Probabilistic diffusion tractography with multiple fibre orientations: What can we gain? Neuroimage 34, 144–155 (2007).

60. Lashkari, D., Vul, E., Kanwisher, N. & Golland, P. Discovering structure in the space of fMRI selectivity profiles. NeuroImage 50, 1085–1098 (2010).

