## Supplementary material for "Stable large-scale network architecture and structured individual variability in the awake mouse brain": All Extended Figures

### Extended Data Fig. 1

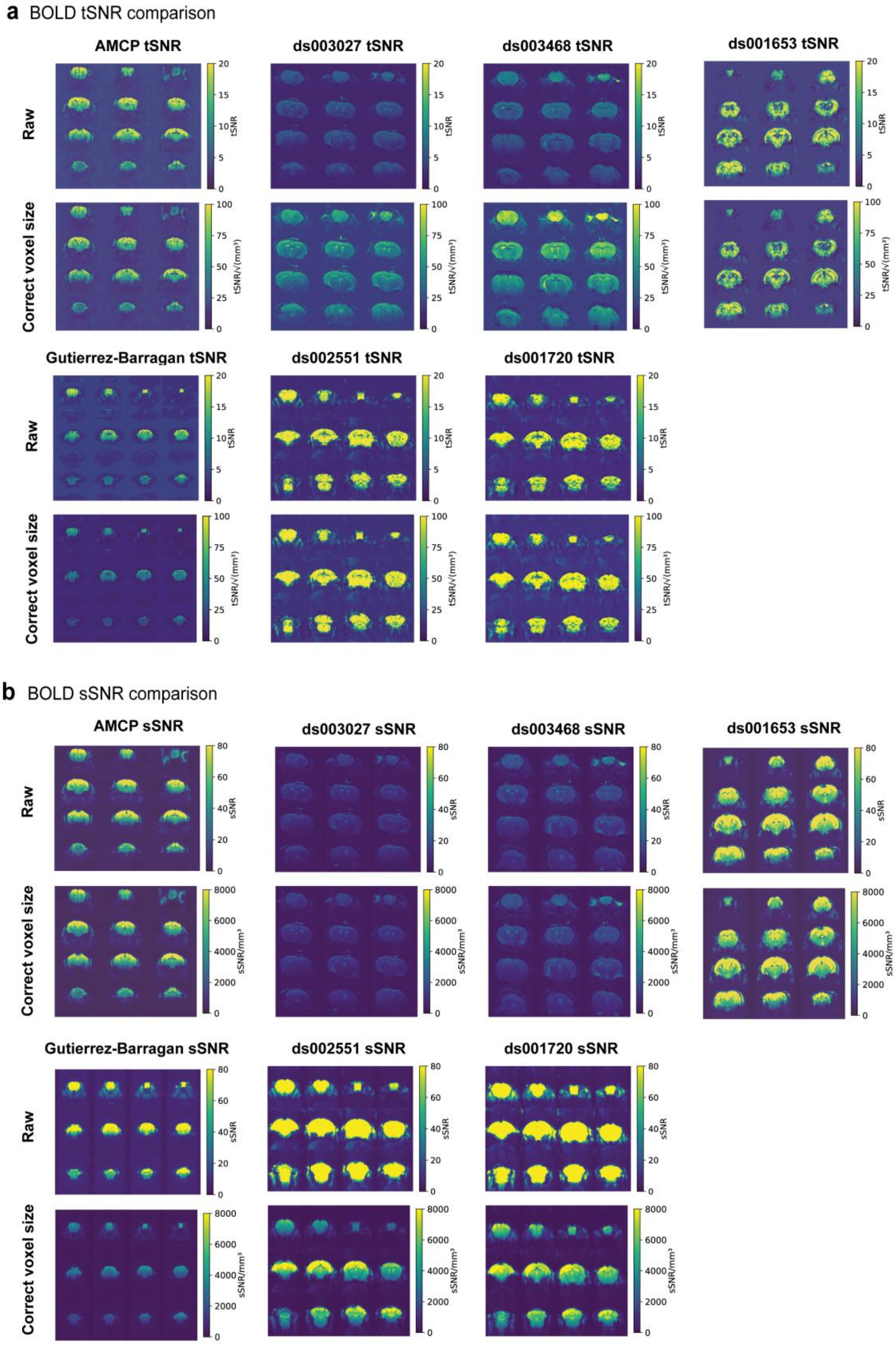


**Extended Data Fig. 1 | Cross-dataset benchmarking of BOLD signal quality.**
**a**, tSNR maps for raw BOLD data from AMCP and three publicly available awake mouse resting-state fMRI datasets. The top row shows uncorrected tSNR, and the

bottom row shows tSNR after voxel-size correction to account for differences in spatial resolution across datasets. **b**, sSNR maps for the same datasets. The top row shows uncorrected sSNR, and the bottom row shows voxel-size-corrected sSNR. Voxel-size correction was performed by normalizing measured SNR values by the square root of voxel volume. These comparisons show that the AMCP cohort maintains high BOLD signal quality relative to existing public awake mouse fMRI datasets. tSNR, temporal signal-to-noise ratio; sSNR, spatial signal-to-noise ratio; SNR, signal-to-noise ratio.

### Extended Data Fig. 2

**a** Frame-wise displacement across datasets

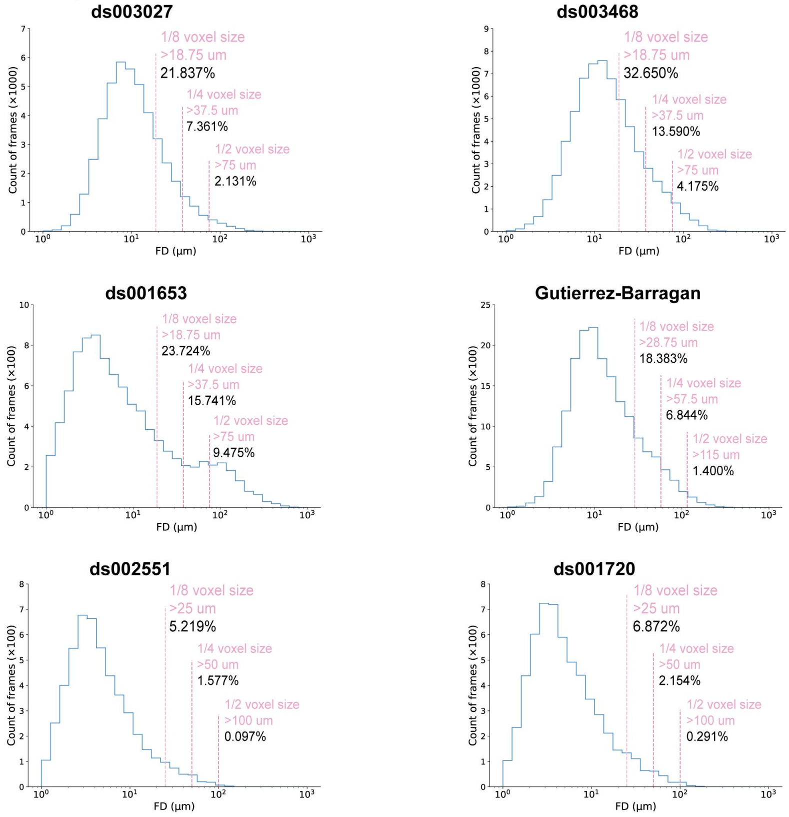


**b** Acquisition parameters and data scale across datasets

| | Device | Surface Coil | TR (ms) | TE (ms) | Resolution (mm3) | Slices Orientation | GE/ SE | fMRI Volumes | N (sex) |
|---|---|---|---|---|---|---|---|---|---|
| AMCP | uMR 9.4T | 4-ch-RT | 1000 | 15.9 | 0.15*0.15*0.5 | 28/Coronal | GE | 1920*212 | 212 (99Male/113Female) |
| ds003027 | Bruker 9.4T | single loop | 2500 | 18.398 | 0.15*0.15*0.45 | 30/Coronal | SE | 7733*6 | 6 (-) |
| ds003468 | Bruker 9.4T | single loop | 2500 | 18.398 | 0.15*0.15*0.45 | 30/Coronal | SE | 5354*13 | 13 (Male) |
| ds001653 | Bruker 11.7T | 4-ch-Cryo | 1000 | 11.7 | 0.19*0.15*0.4 | 28/Coronal | GE | 1080*9 | 9 (Female) |
| Gutierren-Barragan | Bruker 7.0T | 4-ch-RT | 1000 | 15 | 0.23*0.23*0.6 | 18/Coronal | GE | 1920*10 | 10 (Male) |
| ds002551 | Bruker 7.0T | 2-ch-Cryo | 1500 | 15 | 0.2*0.2*0.75 | 18/Coronal | GE | 800*6 | 6 (Male) |
| ds001720 | Bruker 7.0T | 2-ch-Cryo | 1500 | 15 | 0.2*0.2*0.75 | 18/Coronal | GE | 390*15 | 15 (Female) |

- means not reported; ch: channels; RT: room temperatrue; Cryo: cryogenic.

**Extended Data Fig. 2 | Head motion, acquisition parameters and dataset scale.**
**a**, FD across all BOLD volumes in AMCP and three publicly available awake mouse resting-state fMRI datasets. Volumes exceeding half the voxel size are indicated as high-motion volumes. AMCP showed a markedly low proportion of high-motion volumes, indicating minimal head motion during awake fMRI acquisition. **b**, Comparison of BOLD-EPI acquisition parameters and dataset scale across AMCP and the three public datasets, including spatial resolution, repetition time, number of animals and total number of imaging volumes. AMCP provides comparable or higher spatial and temporal resolution than existing public awake mouse fMRI datasets while

substantially increasing sample size and total data volume. FD, framewise displacement.

### Extended Data Fig. 3

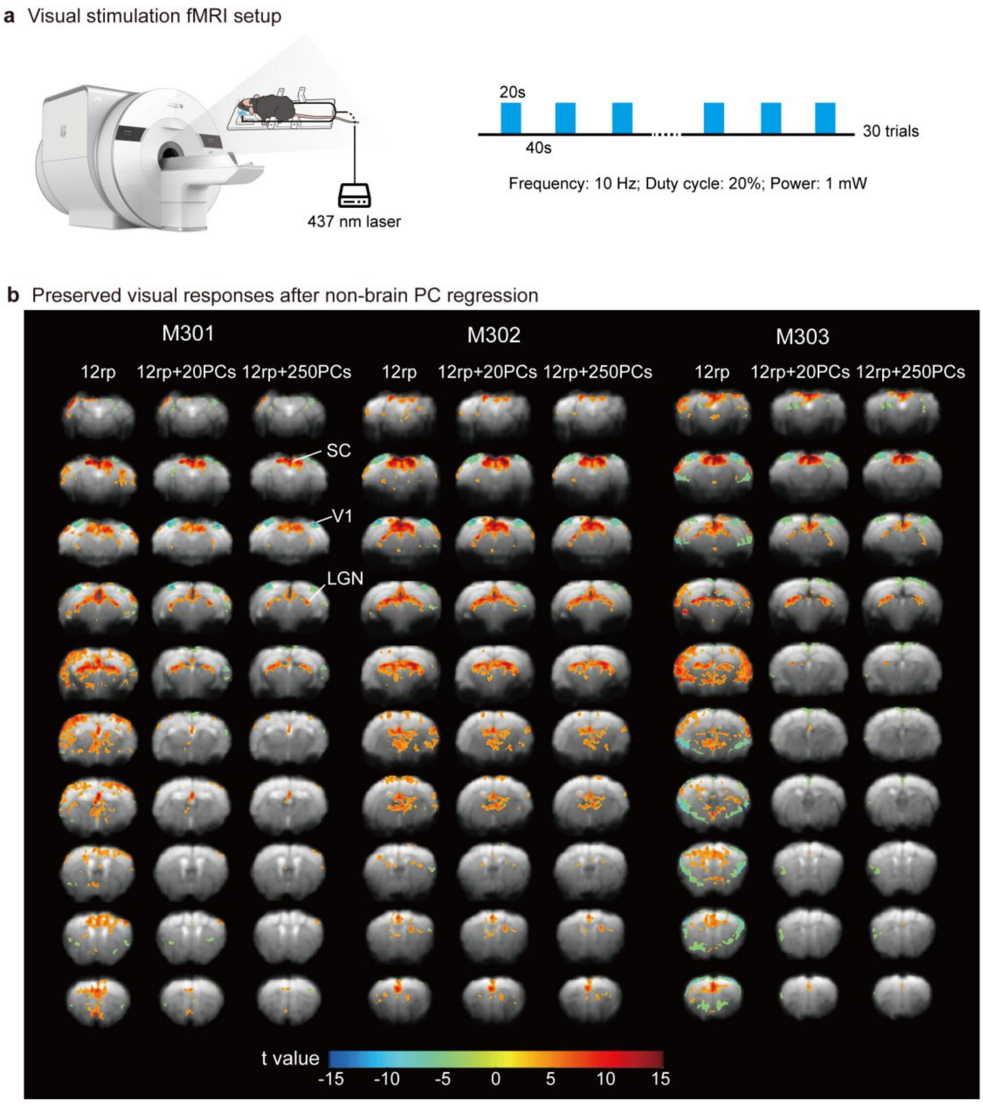


**Extended Data Fig. 3 | Visual-evoked fMRI responses are preserved after regression of non-brain principal components.**

**a**, Visual-stimulation fMRI setup and block-design paradigm. Awake mice were scanned using the same head-fixation setup and BOLD-EPI sequence as in the main AMCP dataset. Bilateral visual stimulation was delivered with a 437-nm laser through optical fibers positioned approximately 1 cm from each eye. Stimulation blocks lasted 20 s and were followed by 40-s inter-stimulation intervals, repeated for 30 trials. Stimulation was delivered at 10 Hz with a 20% duty cycle and 1 mW light power. **b**, Whole-brain visual-evoked activation maps from three mice after denoising with 12 motion-related regressors only, 12 motion-related regressors plus 20 non-brain PCs, or 12 motion-related regressors plus 250 non-brain PCs. Voxel-wise t-maps were thresholded at FDR-corrected $P < 0.05$ with a minimum cluster size of 20 voxels.

Activation in the expected visual pathway, including the primary visual cortex, superior colliculus and lateral geniculate nucleus, was preserved after 250-PC regression, whereas nonspecific activation outside this pathway was reduced. M301-M303 indicate individual mice. rp, rigid-body realignment parameters; PC, principal component; SC, superior colliculus; V1, primary visual cortex; LGN, lateral geniculate nucleus.

### Extended Data Fig. 4

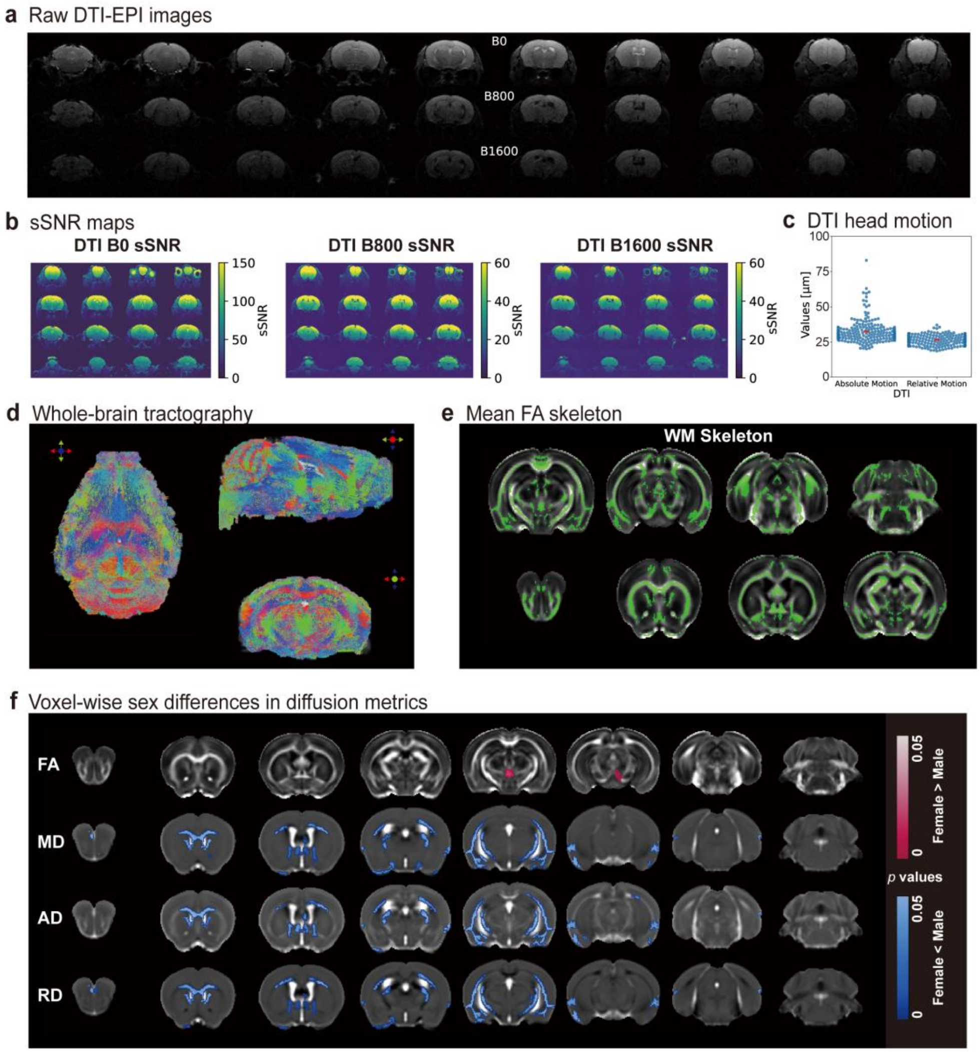


**Extended Data Fig. 4 | DTI data quality, population diffusion templates and sex differences in diffusion metrics.**

**a**, Representative raw DTI-EPI images from the same slice acquired at b = 0, 800 and 1,600 s $mm^{-2}$, showing minimal geometric distortion and artifacts. **b**, Whole-brain sSNR maps for the b = 0, 800 and 1,600 s $mm^{-2}$ volumes. **c**, Framewise displacement across all DTI volumes, indicating low head motion during diffusion imaging. **d**, Representative whole-brain deterministic tractography from one mouse. **e**, Population-averaged white-matter skeleton derived from tract-based spatial statistics, representing major white-matter pathways shared across the cohort. **f**, Voxel-wise sex differences in diffusion metrics between female (n = 113) and male (n = 99) mice. Statistical maps show two-sample t-tests for FA, MD, AD and RD, thresholded at cluster-level FDR-corrected $P < 0.05$ and overlaid on the corresponding AMCP population diffusion templates. sSNR, spatial signal-to-noise ratio; FA, fractional anisotropy; MD, mean diffusivity; AD, axial diffusivity; RD, radial diffusivity.

### Extended Data Fig. 5

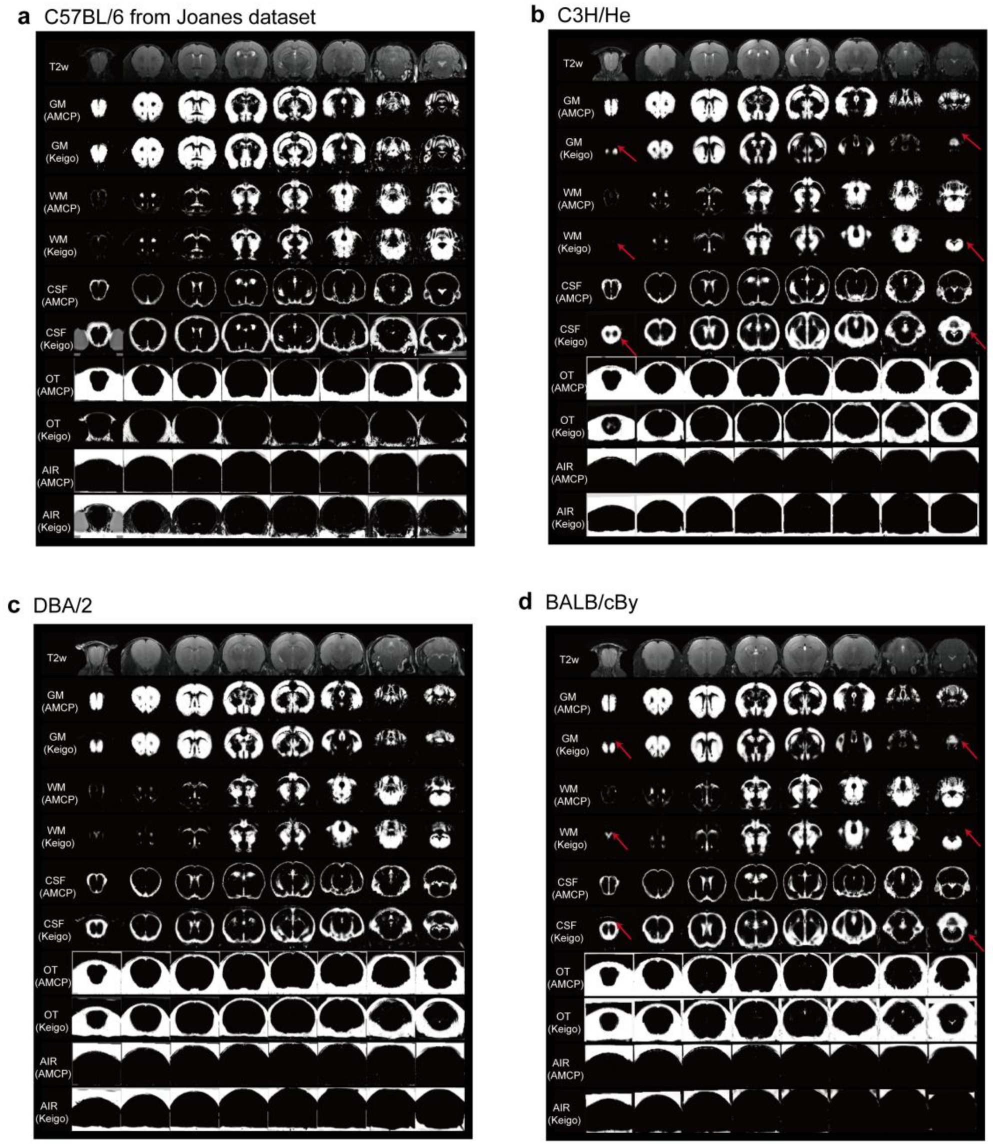


**Extended Data Fig. 5 | Cross-dataset and cross-strain validation of the AMCP tissue probability template.**

**a**, Segmentation of an independent C57BL/6 T2-weighted image using the AMCP and Keigo tissue priors. The mouse was not included in the AMCP cohort. Tissue maps are shown for GM, WM, CSF, OT and air, illustrating the generalizability of the AMCP priors within the same strain. **b-d**, Cross-strain tissue segmentation in C3H/He, DBA/2 and BALB/cBy mice. T2w images from each strain were segmented using the AMCP and Keigo priors. Red arrows indicate representative regions where the Keigo priors produced anatomically implausible tissue assignments. The AMCP priors retained anatomically plausible tissue boundaries across strains, despite being derived from C57BL/6 mice. GM, gray matter; WM, white matter; CSF, cerebrospinal fluid; OT, other tissue.

### Extended Data Fig. 6

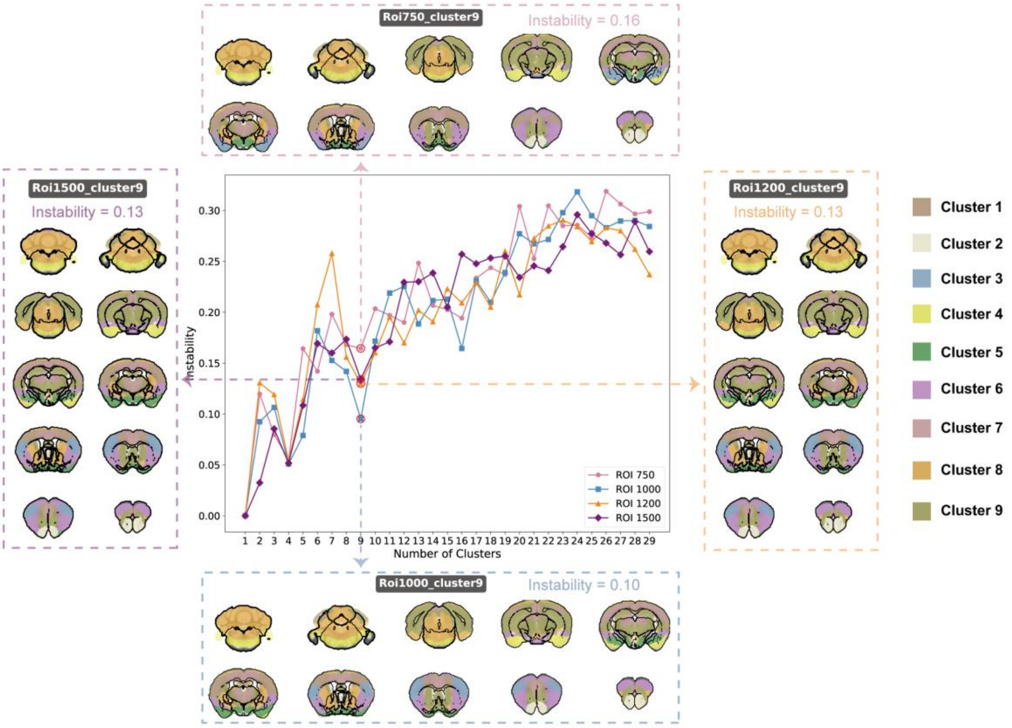


**Extended Data Fig. 6 | Functional network construction using uniform whole-brain parcellations.**

Whole-brain voxels were first grouped into uniformly sized regions using k-means clustering, and voxel-to-region connectivity matrices were then clustered to derive functional networks. The instability curves identified a 9-network solution as the stable operating point across uniform parcellations with 750, 1,000, 1,200 and 1,500 regions. The resulting 9-network maps showed consistent spatial topography across region counts and substantial overlap with the AMCP 11-network atlas derived from Allen CCF-based prior regions (Fig. 3). Differences between the uniform-region and Allen-prior solutions included assignment of the olfactory bulb with the ESN rather than limbic cortex, and a more spatially restricted SMN in the uniform-region solution. Allen CCF, Allen Common Coordinate Framework; ESN, extended salience network; SMN, sensorimotor network.

### Extended Data Fig. 7

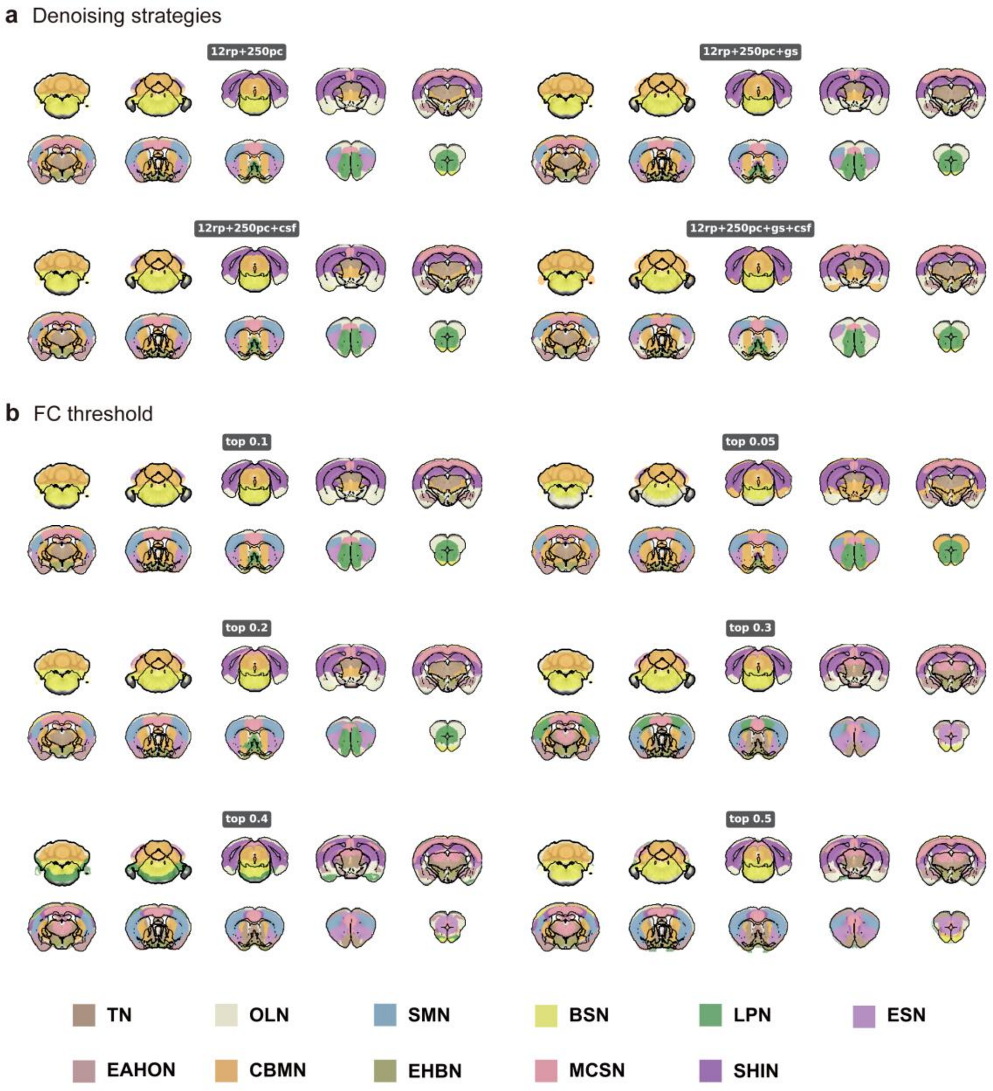


**Extended Data Fig. 7 | Robustness of AMCP network topography to threshold and preprocessing choices.**

**a**, Preprocessing sensitivity of the atlas. Representative slices show functional network maps derived using different nuisance-regression strategies. **b**, Threshold sensitivity of the AMCP 11-network atlas. Representative slices show functional network maps derived using different connectivity thresholds for retaining voxel-to-region connections before clustering. The large-scale network topography was broadly preserved across thresholds, with only modest local boundary differences. The major network organization remained stable across preprocessing variants, indicating that the principal atlas structure was not driven by a single denoising choice.

### Extended Data Fig. 8

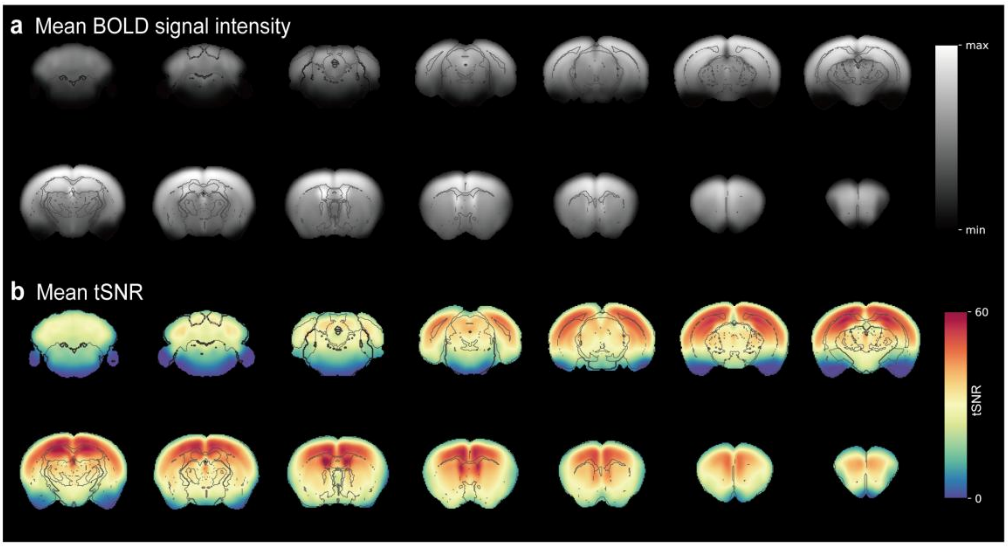


**Extended Data Fig. 8 | Characterization of BOLD signal intensity and tSNR in the common coordinate space.**

**a**,**b**, Mean BOLD signal intensity (**a**) and mean tSNR (**b**) across all subjects after normalization to the common coordinate space. Both maps show reduced signal intensity and lower tSNR at the lateral cortical margins and in ventral brain regions, consistent with common technical features of BOLD fMRI, including air-tissue susceptibility effects and receive-coil sensitivity profiles. These spatial signal properties may contribute to the clustering of some lower-quality boundary and ventral voxels into the OLN; results involving this network should therefore be interpreted with appropriate caution. tSNR, temporal signal-to-noise ratio; OLN, olfactory-limbic network.

### Extended Data Fig. 9

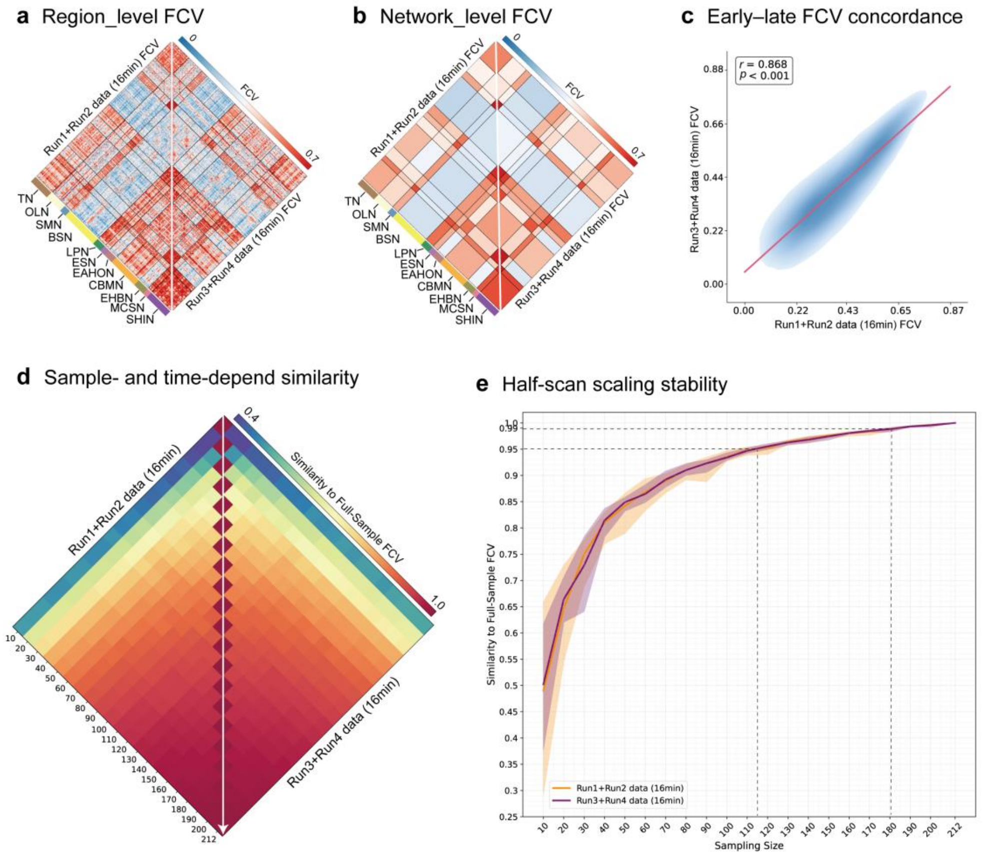


**Extended Data Fig. 9 | Stability of FCV architecture across scan halves.**
**a**,**b**, Region- and network-level FCV matrices computed separately from the first half (Runs 1-2; a) and second half (Runs 3-4; **b**) of the rsfMRI data. **c**, Concordance between early- and late-half FCV matrices, quantified by Pearson correlation across network-level FCV terms. **d**, Sample-size-dependent similarity between subset-derived FCV matrices and the corresponding half-scan reference matrix, computed separately for Runs 1-2 and Runs 3-4. **e**, Half-scan marginal-gain curves for FCV estimation. The close overlap between curves indicates that the sample-size dependence of FCV estimation was stable across early and late portions of the scanning session. Shaded areas indicate the minimum-to-maximum range across samplings, and solid lines indicate the mean. FCV, functional connectivity variability.

### Extended Data Fig. 10

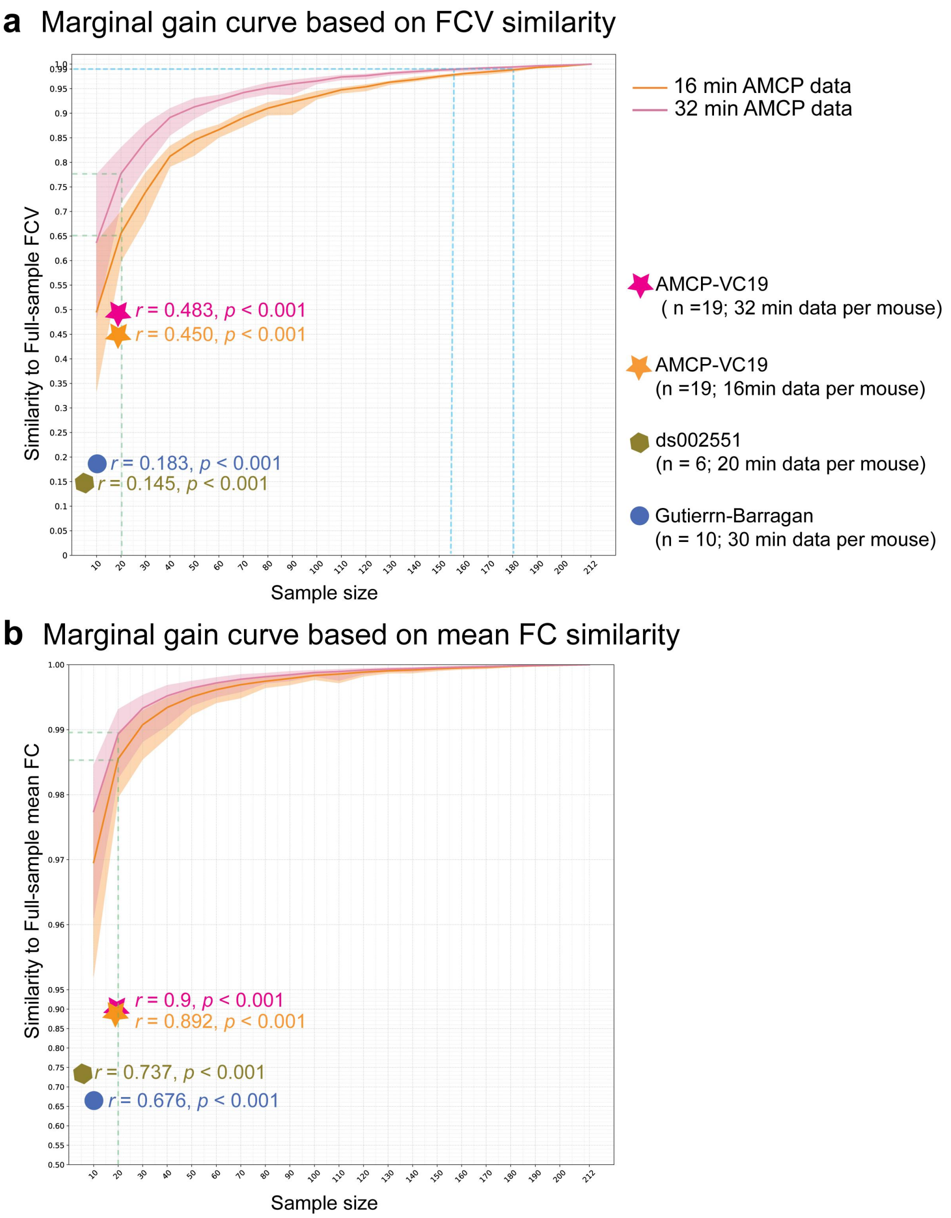


**Extended Data Fig. 10 | External comparison of independent datasets with sample-size marginal-gain curves.**
**a**, External comparison based on FCV similarity. Curves show the relationship between sample size and similarity to the full-sample FCV matrix from the AMCP main cohort, estimated using either 32 min or 16 min of resting-state fMRI data per mouse. Colored markers show independent datasets overlaid on the AMCP-derived curves, including m (AMCP-VC19; n = 19) and two publicly available awake mouse fMRI datasets. **b**, External comparison based on mean FC similarity. Curves show

similarity to the full-sample mean FC matrix from the AMCP main cohort as a function of sample size and scan duration, with the same datasets shown as colored markers. For each marker, Pearson's r denotes the matrix similarity between the independent dataset and the corresponding full-sample matrix from the AMCP main cohort. These projections provide contextual comparisons with the AMCP reference, but they should not be interpreted as external validation of universal sample-size thresholds because independent cohorts can fall below AMCP subsampling expectations when acquisition parameters, data quality or cohort characteristics differ. Shaded areas indicate the minimum-to-maximum range across samplings, and solid lines indicate the mean. FC, functional connectivity; FCV, functional connectivity variability; AMCP-VC19, independent AMCP validation cohort of 19 mice.
