## Supplementary material for "Stable large-scale network architecture and structured individual variability in the awake mouse brain": All Supplementary Figures

### Supplementary Fig. 1

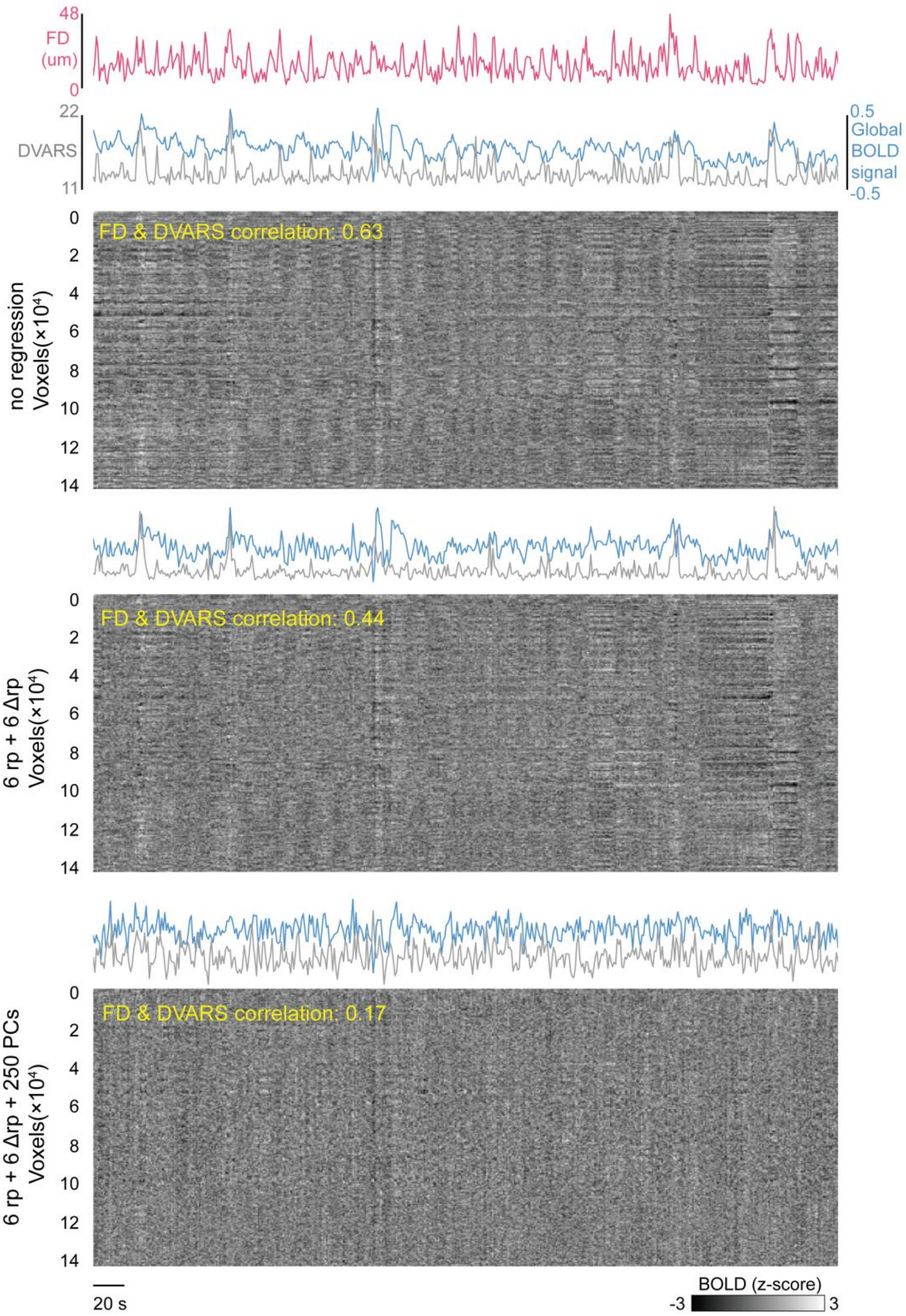


**Supplementary Fig. 1 | Effect of nuisance regression on motion-related and global BOLD fluctuations.**

Representative time-series visualization from one awake mouse rsfMRI run. Top, FD, DVARS and global BOLD signal time courses from the raw data. Middle and bottom, voxel-wise BOLD signal time series shown as intensity plots before regression, after regression of 12 motion-related regressors (6 rp + 6 Δ rp), and after regression of 12 motion-related regressors plus 250 non-brain principal components (6 rp + 6 Δ rp + 250 PCs). Each row represents one brain voxel and each column represents one time point; BOLD signals are shown as z scores. FD-DVARS correlations are labeled for each preprocessing stage. Regression of non-brain PCs markedly reduced global structured fluctuations and FD-DVARS coupling, indicating effective removal of motion-related and non-neural variance while retaining residual brain BOLD time series for downstream analyses. FD, framewise displacement; DVARS, root-mean-square temporal derivative of voxel-wise BOLD signal; rp, rigid-body realignment parameter; PC, principal component.

### Supplementary Fig. 2

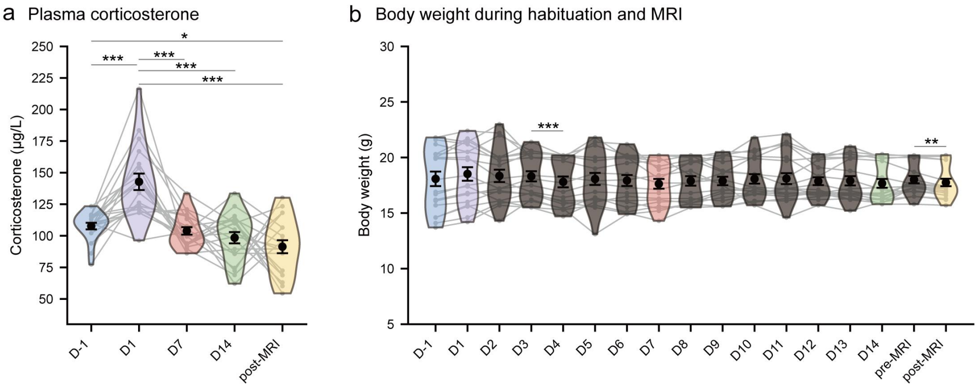


**Supplementary Fig. 2 | Plasma corticosterone and body weight during awake MRI habituation and scanning.**

**a**, Plasma corticosterone levels measured from tail-vein blood samples before habituation (D-1), during habituation training (D1, D7 and D14) and after MRI scanning (post-MRI). **b**, Body weight measured longitudinally across the same experimental period, including baseline (D-1), habituation days (D1 – D14) and immediately before and after MRI scanning. Violin plots show distributions across mice; gray lines connect repeated measurements from the same mouse; black points and error bars indicate mean ± SEM. Statistical significance was assessed using two-sided paired t-tests with Holm-Bonferroni correction across all pairwise comparisons within each outcome. Only significant corrected comparisons are shown. Adjusted $P < 0.05$, adjusted $P < 0.01$, adjusted $P < 0.001$. $n = 19$ mice.

#### Supplementary Fig. 3

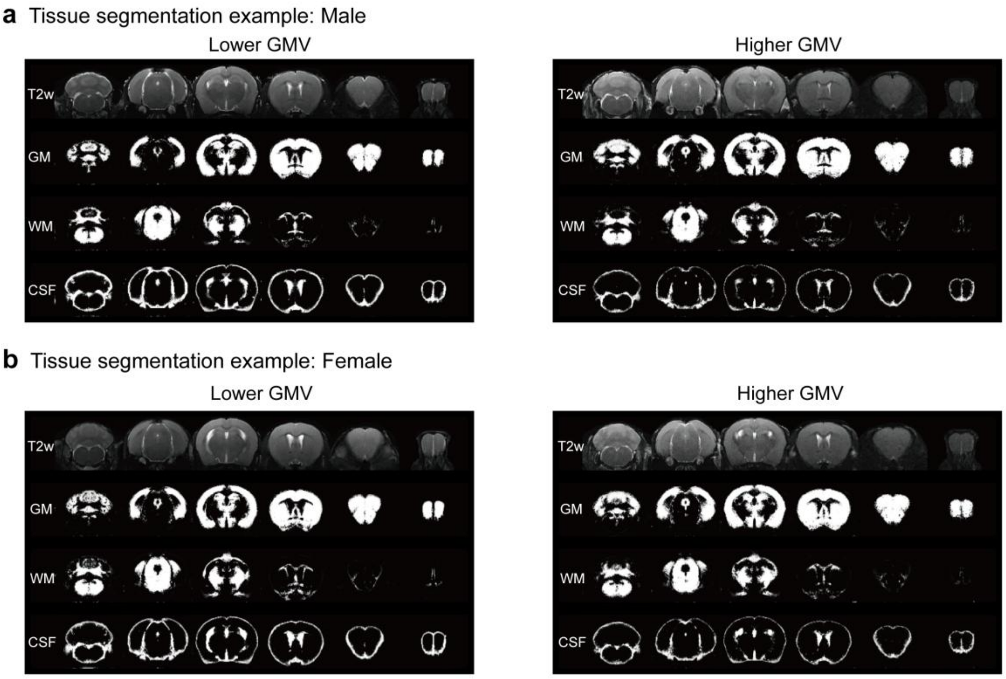


**Supplementary Fig. 3 | Representative segmentation examples across the range of whole-brain GM volume.**

**a**, T2-weighted images and corresponding AMCP-derived GM, WM and CSF segmentations from two male mice with relatively low and high whole-brain GM volume. **b**, T2-weighted images and corresponding segmentations from two female mice with relatively low and high whole-brain GM volume. These representative cases illustrate that animals at the lower and higher ends of the GM-volume distribution showed anatomically plausible tissue boundaries, supporting that the observed inter-individual variation in GM, WM and CSF volumes was not driven by gross segmentation failure. GM, gray matter; WM, white matter; CSF, cerebrospinal fluid.

### Supplementary Fig. 4

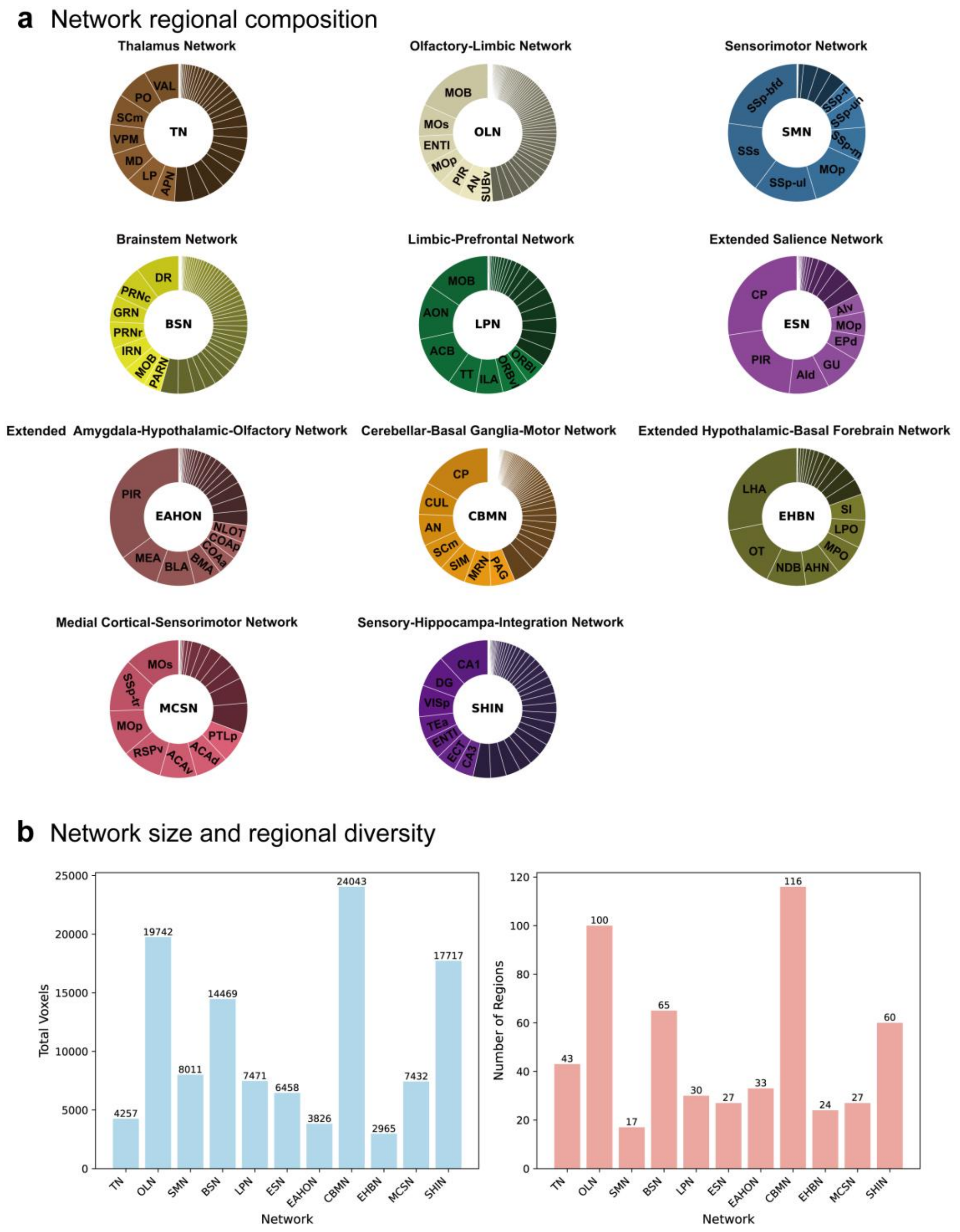


**Supplementary Fig. 4 | Regional composition of the AMCP 11-network atlas.**
**a**, Regional composition of each AMCP functional network based on Allen CCFv3 anatomical labels. Pie charts show the relative voxel contribution of each anatomical region within each network. **b**, Quantitative summary of network composition. Left, total voxel count for each of the 11 networks. Right, number of distinct Allen CCFv3 regions represented within each network. These analyses were used to guide anatomical interpretation and network nomenclature.

### Supplementary Fig. 5

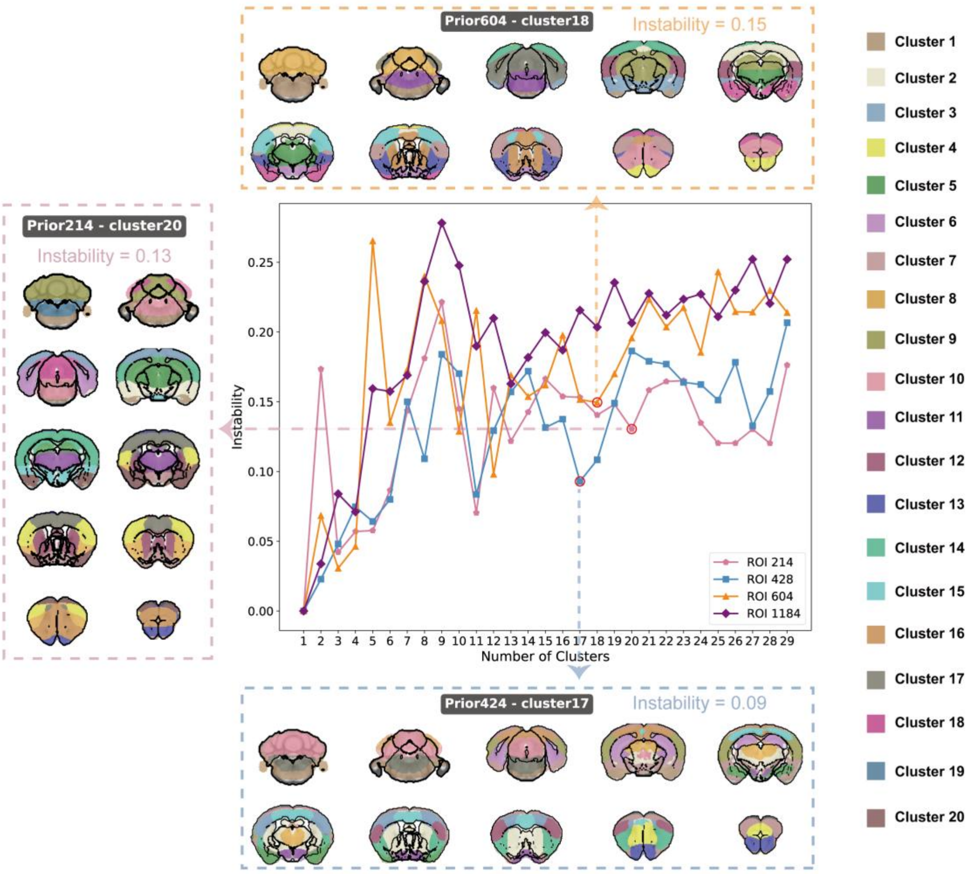


**Supplementary Fig. 5 | Fine-grained functional network solution derived from Allen-prior regions.**

Alternative higher-resolution functional network solution identified from the second prominent elbow of the instability curve in Fig. 3b. This finer-grained solution preserves the global organization of the primary AMCP 11-network atlas while revealing additional subdivisions. Notable refinements include separation of the SHIN into cortical and hippocampal components, subdivision of the BSN into dorsal and ventral components, and subdivision of the SMN into dorsal and ventral components. This atlas provides a complementary higher-granularity network reference for studies requiring finer functional subdivision. SHIN, sensory-hippocampal-integration network; BSN, brainstem network; SMN, sensorimotor network.

### Supplementary Fig. 6

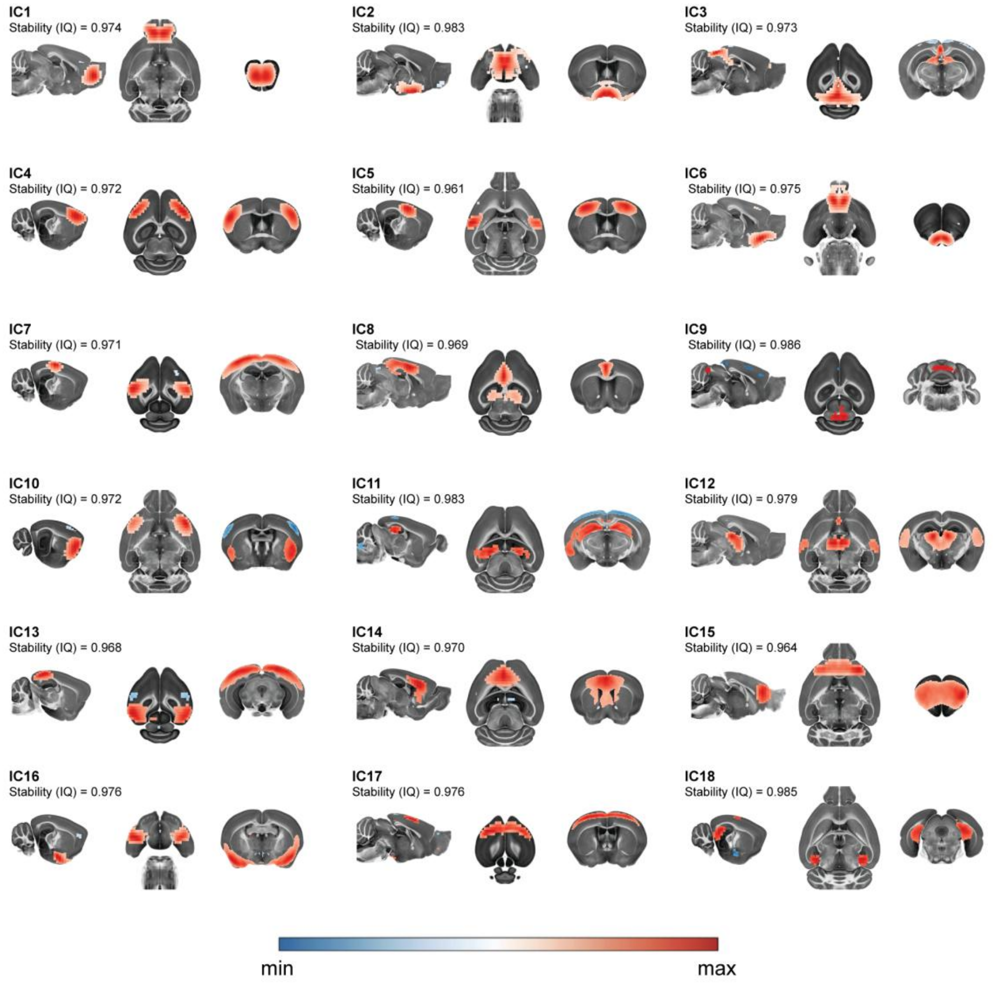


**Supplementary Fig. 6 | Spatial maps of the 18-component GICA solution.**
Spatial maps of the 18 GICA-derived components are shown as a lower-dimensional comparison to the 20-component solution in Fig. 5. Component maps were converted to Z scores and thresholded at $|Z| > 1.96$ for visualization. Most components showed spatial patterns broadly consistent with the 20-component solution, but the 18-component solution showed reduced anatomical specificity in selected components, including hemispheric asymmetry in some maps and mixed anatomical loading across functionally distinct regions. For example, IC11 showed marked left-right asymmetry, and IC14 included both motor-related regions and gray-matter regions near the ventral ventricles. In addition, a ventral hippocampus-dominant component was not clearly separated in this model order. These observations support the use of the 20-component solution as the representative GICA model order in the main analysis. GICA, group independent component analysis; IC, independent component; IQ, ICASSO stability index.

### Supplementary Fig. 7

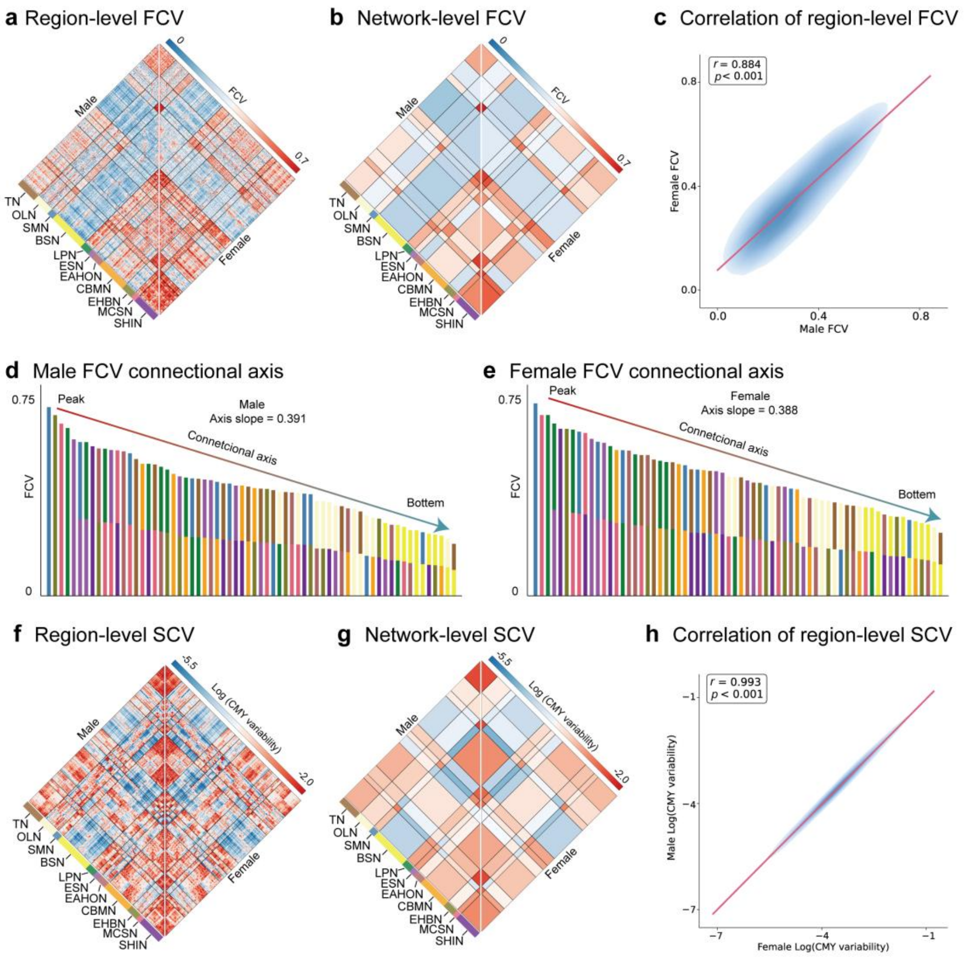


**Supplementary Fig. 7 | Sex-wise comparison of functional and structural connectivity variability.**

**a**-**b**, Inter-individual FCV matrices shown at the region level using 214 regions (**a**) and at the AMCP 11-network level (**b**). The lower triangle shows female mice (n = 113), and the upper triangle shows male mice (n = 99). **c**, Spatial correspondence of region-level FCV between sexes, quantified by Pearson correlation. **d**,**e**, Sex-wise connectional axes of network-level FCV. Network-level FCV terms were ranked from highest to lowest separately in male (**d**) and female (**e**) mice, and the axis slope was estimated by linear regression on the ranked values. **f**,**g**, Inter-individual SCV matrices shown at the region level (**f**) and network level (**g**), displayed as in **a** and **b**. **h**, Spatial correspondence of region-level SCV between sexes, quantified by Pearson correlation. SCV showed very high similarity between female and male mice, whereas FCV showed a slightly lower but still strong sex-wise correspondence. FCV, functional connectivity variability; SCV, structural communicability variability.

### Supplementary Fig. 8

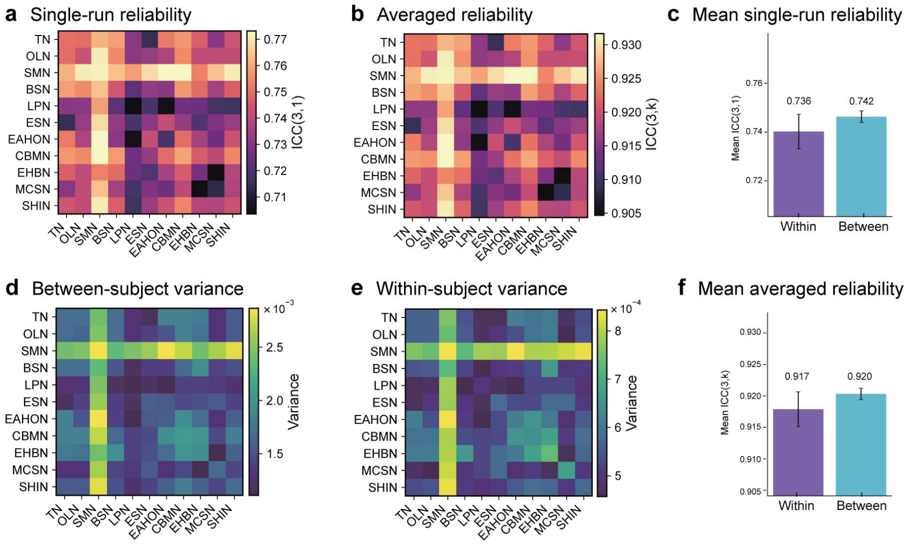


**Supplementary Fig. 8 | Repeated-run reliability and variance components of network-level functional connectivity.**

**a**, Single-run reliability of the 66 network-level FC terms, comprising 11 within-network and 55 between-network terms, quantified using ICC(3,1). **b**, Averaged-measure reliability of the same 66 FC terms across repeated runs, quantified using ICC(3,k). **c**, Mean single-run reliability for within- and between-network FC terms. **d**,**e**, Between-subject variance (**d**) and within-subject/error variance (**e**) estimated from repeated-measures ANOVA for each network-level FC term. **f**, Mean averaged-measure reliability for within- and between-network FC terms. FC, functional connectivity; ICC, intraclass correlation coefficient.

### Supplementary Fig. 9

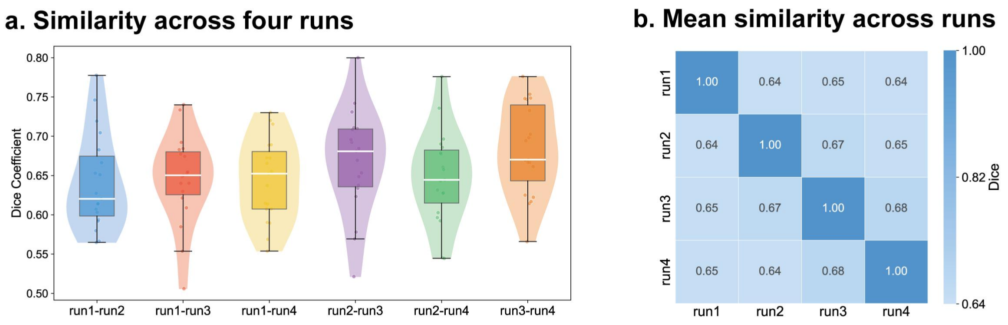


**Supplementary Fig. 9 | Reproducibility of individualized functional network parcellation in the independent AMCP validation cohort.**

**a**, Within-subject Dice similarity of individualized parcellation maps across all pairwise combinations of four repeated rsfMRI runs. Each point represents one mouse, boxes indicate the interquartile range and median, and violins show the distribution across subjects. **b**, Mean within-subject Dice similarity matrix across repeated runs, summarizing test-retest stability of individualized network assignments in the validation cohort. Higher Dice values indicate greater spatial overlap between individualized parcellation maps. rsfMRI, resting-state functional MRI.

#### Supplementary Fig. 10

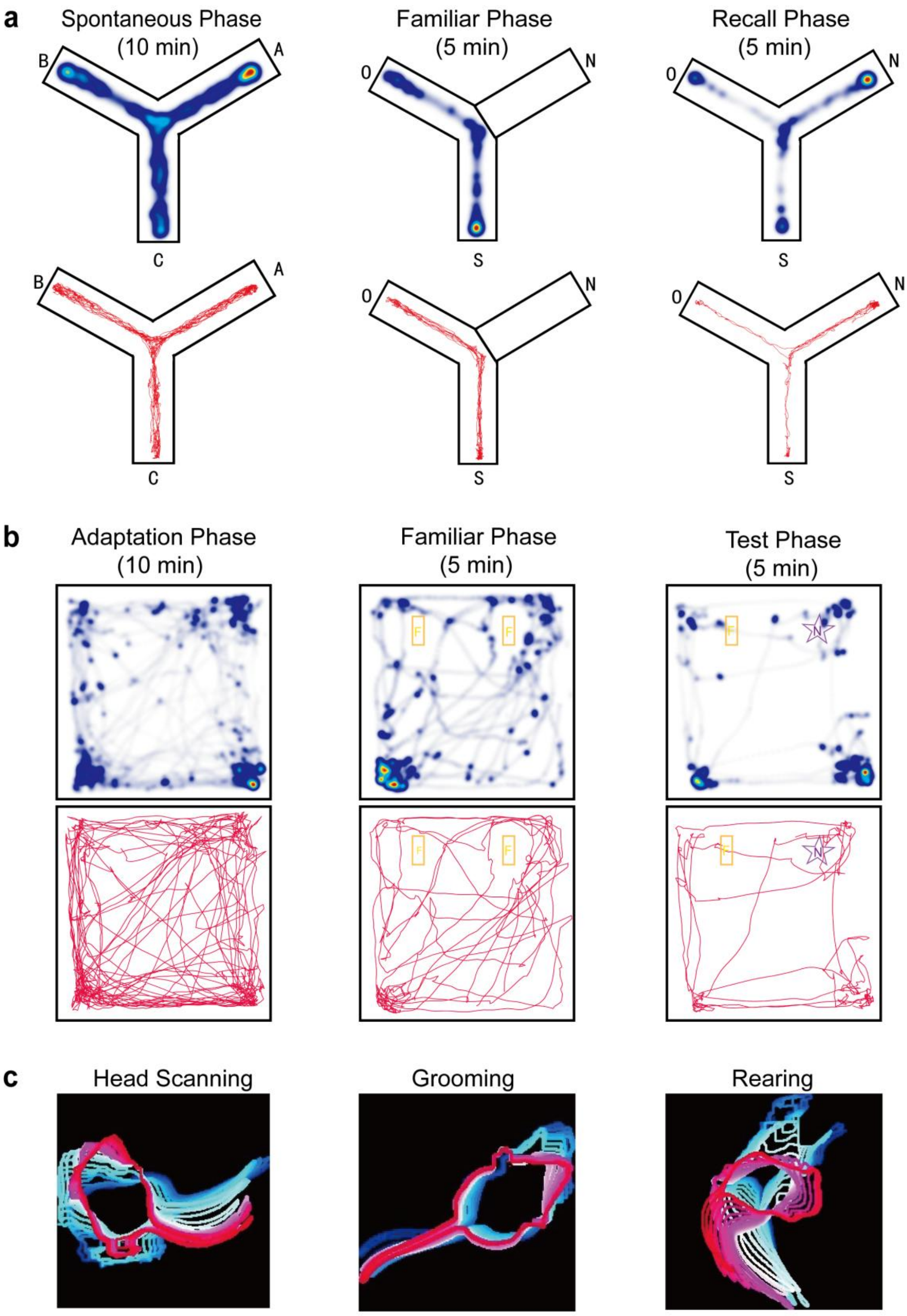


**Supplementary Fig. 10 | Representative behavioral trajectories, spatial occupancy and automated behavior features.**

**a**, Representative Y-maze behavioral data across three task phases. Spatial occupancy heatmaps are shown in the top row, and the corresponding raw movement trajectories are shown in the bottom row. **b**, Representative open-field behavioral data across three task phases, displayed as in **a**. **c**, Example LabGym pattern image used for automated behavioral quantification, showing the temporal organization of classified behavioral events.

### Supplementary Fig. 11

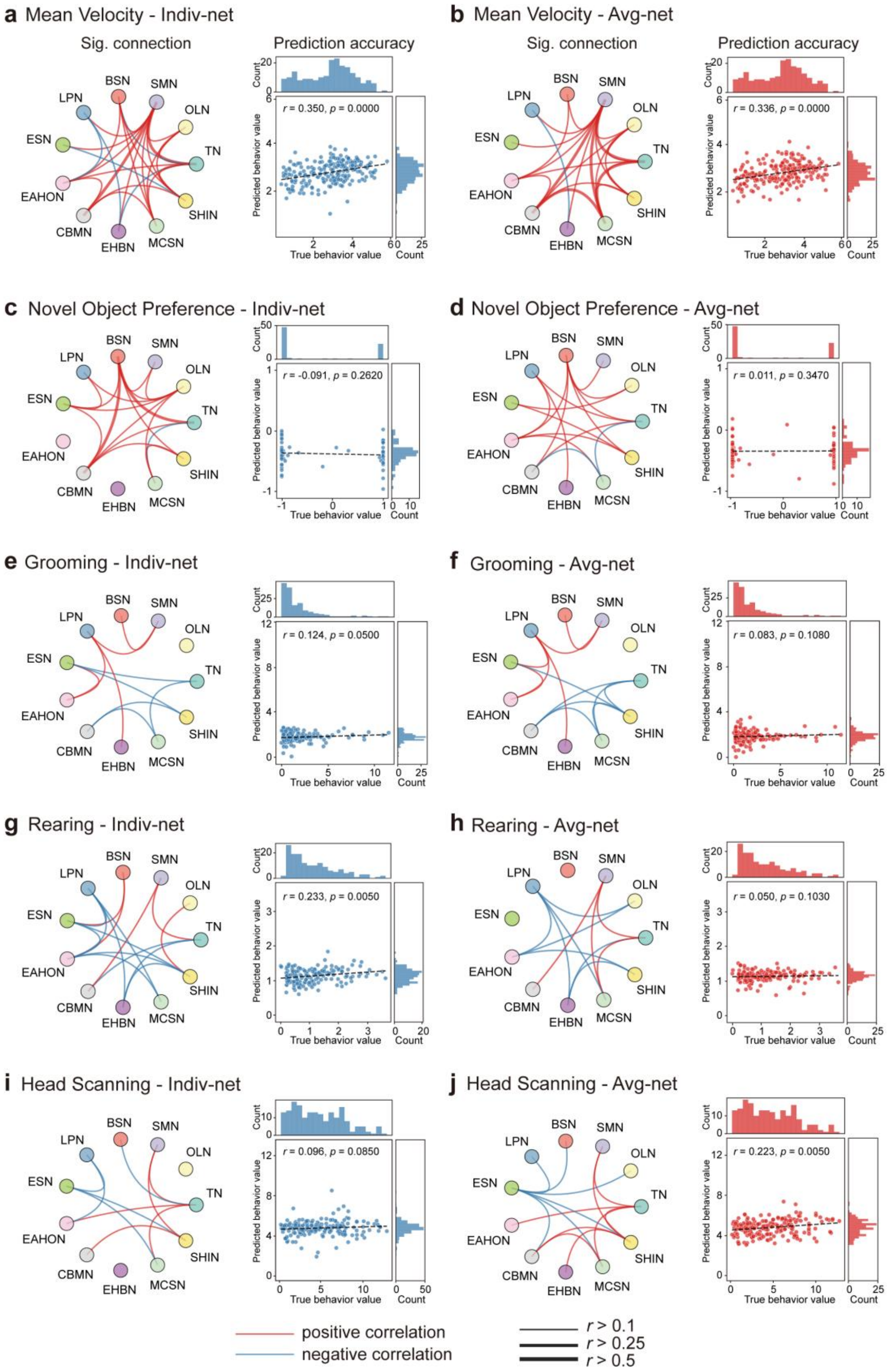


**Supplementary Fig. 11 | Additional examples of behavioral prediction using individualized and group-average network features.**

Representative prediction results for five additional behavioral outcomes, complementing Fig. 8. For each behavior, significant network connections and the correlation between observed and predicted scores are shown for Indv-Net features and Avg-Net features. **a**,**b**, Mean velocity. **c**,**d**, Novel object preference. **e**,**f**, Grooming. **g**,**h**, Rearing. **i**,**j**, Head scanning. Indv-Net, individualized parcellation-based network; Avg-Net, group-average atlas-based network.

### Supplementary Fig. 12

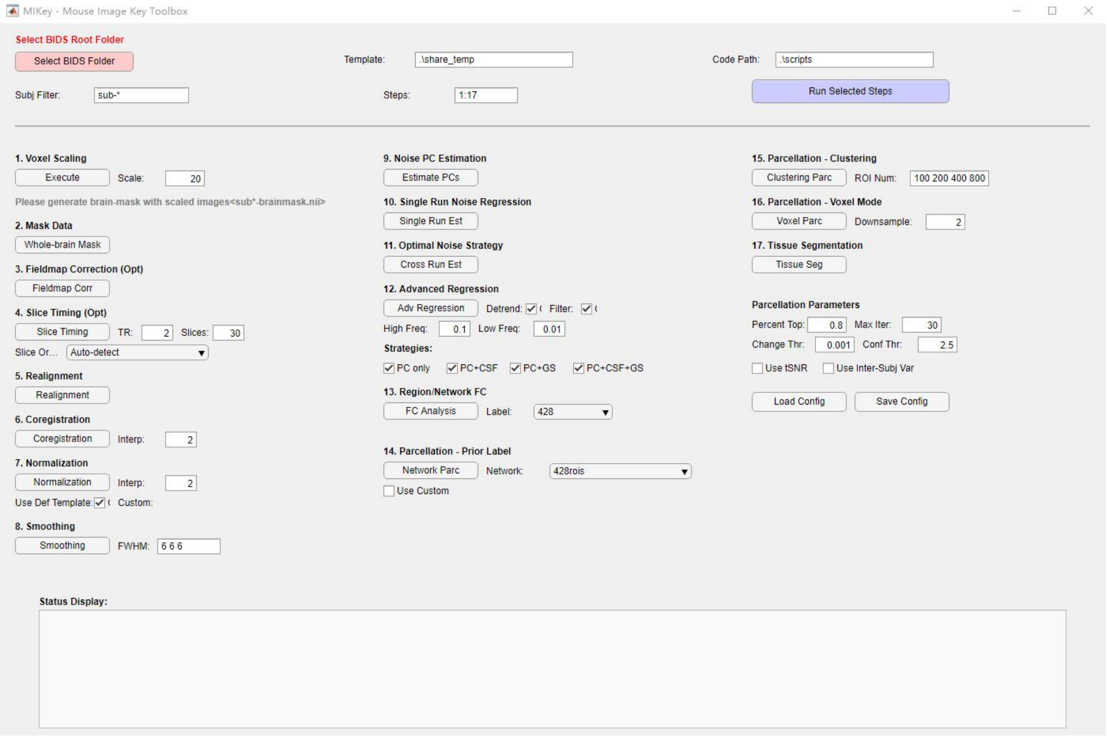


**Supplementary Fig. 12 | MIKey graphical interface for AMCP-based mouse MRI analysis.**

Representative screenshots of the MIKey toolbox graphical interface. MIKey is an open-source MATLAB/SPM12-based toolbox that integrates AMCP-derived resources, including tissue probability maps and functional network atlases, with workflows for mouse MRI preprocessing, post-processing and individualized network analysis. Users provide an initial brain mask, which can be generated using MouseBrainExtractor followed by manual refinement in ITK-SNAP. Individual modules can be executed separately, or selected preprocessing and analysis steps can be run sequentially through the graphical interface. This interface is designed to facilitate application of AMCP resources by users with different levels of programming experience.
